# Structure-function relationship of alpha-synuclein fibrillar polymorphs derived from distinct synucleinopathies

**DOI:** 10.1101/2025.10.13.682210

**Authors:** Tetiana Serdiuk, Virginie Redeker, Jimmy Savistchenko, Sandesh Neupane, Walther Haenseler, Yanick Fleischmann, Viviane Reber, Sabrina Keller, Cinzia Tiberi, Ruxandra Bachmann-Gagescu, Matthias Gstaiger, Thomas Braun, Roland Riek, Steve Gentleman, Adriano Aguzzi, Natalie de Souza, Ronald Melki, Paola Picotti

## Abstract

The aggregation of the protein alpha-synuclein (αSyn) is a common feature of multiple neurodegenerative diseases collectively called synucleinopathies, for which the pathobiology is not well understood. The different phenotypic characteristics of the synucleinopathies Parkinson’s disease (PD), Dementia with Lewy Bodies (DLB) and Multiple System Atrophy (MSA) have been proposed to originate from the distinct structures adopted by αSyn in its amyloid forms. Here, using covalent labeling and limited proteolysis coupled to mass spectrometry (LiP-MS) *in vitro* and *in situ* within neuronal cells and directly in native patient brain homogenates, we show that pathogenic αSyn from distinct synucleinopathies (PD, DLB and MSA) are structurally different. Further, we found that fibrillar structural differences are associated with different fibril interactomes and neuronal responses. We discovered disease-specific ubiquitination patterns and turnover profiles for pathogenic αSyn species, detected molecular pathways responding specifically to the uptake of different αSyn fibrillar polymorphs, and identified a subset of the involved proteins as candidate direct interactors of αSyn. In particular, components of the Ubiquitin-proteasomal System (UPS), including E3 ubiquitin ligases, chaperones, and Deubiquitinating proteins, showed disease/polymorph-specific interaction patterns, possibly accounting for different resistance of patient-derived αSyn fibrils to degradation. Genetic modulation with CRISPR-based tools showed that members of the UPS degradation pathway (three E3 ligases: UBE3A, TRIM25, HUWE1 and the AAA+ ATPase VCP) reduced αSyn inclusions, in a strain-specific manner. LiP-MS also identified sets of proteins with altered protease susceptibility in postmortem brain homogenates of PD, DLB, and MSA patients. These sets were largely disease-specific and included proteins altered in cells treated with fibrils derived from patients with the matching disease. Our findings provide insight into cellular processes involved in the accumulation and turnover of αSyn pathogenic aggregates in PD, DLB and MSA in a disease specific manner and constitutes a resource of potential novel drug targets in these synucleinopathies.

## Introduction

Alpha-synuclein (αSyn) aggregation is a characteristic feature of the neurodegenerative diseases, collectively called synucleinopathies^1-4^, that include Parkinson’s disease (PD), Dementia with Lewy Bodies (DLB), and Multiple System Atrophy (MSA). In all these diseases, αSyn aggregation is seen as a driving force of disease initiation, progression, spatial spread of pathology in the brain, and neuronal loss. As previously described for prions, αSyn aggregates were shown to spread from one neuron to another^5-7^. During this prion-like propagation, aggregated fibrillar ‘seeds’ of αSyn bind to membrane proteins, enter a healthy cell by hijacking endocytosis, escape from the endo-lysosomal pathway, and are released into the cytosol, where they form pSer129-and ubiquitin-positive inclusions that also stain with amyloid dyes upon multiplication through the recruitment of endogenous αSyn^8-13^. Multiple cellular quality control (QC) systems are thought to be involved in clearing these aggregates from neurons, but incomplete disaggregation of αSyn aggregates by molecular chaperones (e.g., DNAJB-Hsp70) may also favor propagation through fibril fragmentation and generation of small seeds with higher seeding capability^14,15^. As for prion diseases, endogenous αSyn is crucial for forming intracellular inclusions and for subsequent phenotypic effects, including disease manifestation in mouse models^12,16,17^. Notably, cell exposure to monomeric αSyn does not cause the formation of insoluble αSyn inclusions^16^.

Despite the central role of αSyn in PD, DLB, and MSA, these synucleinopathies differ strongly in their clinical and pathological phenotypes^17-25^. One possible explanation for this diversity is that αSyn forms structurally different strains that may target distinct neuronal cell populations within the brain and/or exhibit differential toxicity through interference with different cellular pathways. The concept of different strains originates from the prion field, where the different structures adopted by the prion protein (PrP) in its misfolded state were proposed to affect distinct brain regions, thus causing different disease phenotypes^26,27^. Similarly, well–characterized *in vitro* generated αSyn strains were shown to amplify in a strain-specific manner within the rat brain^28^, to cause distinct synucleinopathies in rat^8^, and to spread and target neuronal cells differentially after injection into the olfactory bulb of mice^29^. Furthermore, cryo-electron microscopy (cryo-EM) studies of Sarkosyl-insoluble αSyn fibrils isolated from post-mortem brain samples of PD, DLB, and MSA support the existence of distinct polymorphs in Lewy bodies in PD and DLB or Glial Cytoplasmic inclusions (GCI) in MSA^30,31^.

We previously showed that αSyn fibrils derived from PD, DLB, and MSA patient brains through a Protein Misfolding Cyclic Amplification (PMCA) method exhibit structural differences that translate into different pathogenic phenotypes in rats, and importantly, that amplified fibrils could faithfully capture strain-specific effects of brain homogenates themselves^32^. To understand the αSyn strain structure-pathology relationship at the molecular level, here we characterized structural differences between αSyn strains amplified from PD, DLB and MSA patient samples *in vitro*, in cell lysates, within intact cells and in iPSCs-derived cortical neurons, and compared their stability and their cellular interactomes. We further analysed proteome alterations in response to the internalization of αSyn strains derived from different synucleinopathies using limited proteolysis coupled to mass spectrometry (LiP-MS), a structural proteomic approach that we previously showed to probe cellular alterations in complex lysates^33^. LiP-MS creates structure-specific proteolytic fingerprints for every detectable protein in the sample, reporting on numerous classes of molecular events, including protein-small molecule interactions, protein-protein interactions, aggregation, post-translational modifications, allosteric regulation, and altered enzymatic activity^33-35^. It is thus an attractive approach to detect both strain-specific structural changes in αSyn itself as well as to characterize strain-specific cellular interactomes and global functional responses.

Our results demonstrate that αSyn strains derived from PD, DLB, and MSA patient brains differ structurally *in vitro*, in SH-SY5Y cell lysates and in neurons. We show that they have distinct interactomes and trigger different functional responses following their uptake by cells ranging from SH-SY5Y to iPSCs-derived cortical neurons. We further observe that αSyn strains derived from PD, DLB, and MSA patients exhibit distinct ubiquitination patterns, turnover rates, and trigger structural responses in different sets of degradation pathway components (E3 ligases, Deubiquitinating proteins (DUBs), ubiquitin-binding proteins, and chaperones) in cells and neurons. CRISPR-based genetic activation of selected candidate regulators of αSyn strain turnover validated their ability to modulate phosphorylated S129 αSyn inclusions, in a strain specific manner. Our work presents a resource of potential novel drug targets for synucleinopathies that can enhance clearance of αSyn aggregates.

## Results

### Structural differences between disease-derived αSyn strains in vitro

Partial structures of αSyn fibrillar polymorphs generated *de novo* or purified from patient brain homogenates have been solved by structural methods such as cryo-EM or solid state-NMR^30,31,36-38^, but they lack information on dynamic amino acid stretches such as the C terminus. These dynamic regions represent a substantial fraction of the surfaces of these polymorphs and may account for differences between distinct synucleinopathies. Here, we analysed the structural differences between αSyn fibrillar polymorphs derived from patients along the entire sequence of αSyn using a mass spectrometric approach. We applied two structural proteomic methods, covalent molecular painting^39-43^ and LiP-MS, to study the structures of polymorphs both in a buffer (Figure 1) and in a cellular lysate (Figure 2). Patient-derived αSyn polymorphs were obtained from brain homogenates of PD, DLB, or MSA patients (n ≥ 3 patients each)^32,36^ using an adapted version of the PMCA method we recently implemented (Methods)^32^. Brain homogenates from the cingulate cortex were used for PD and DLB cases, and from the cerebellum for MSA cases. We have previously shown that homogenates from the same brain regions originating from healthy individuals did not yield fibrillar assemblies under our experimental conditions^32^.

**Figure 1.**
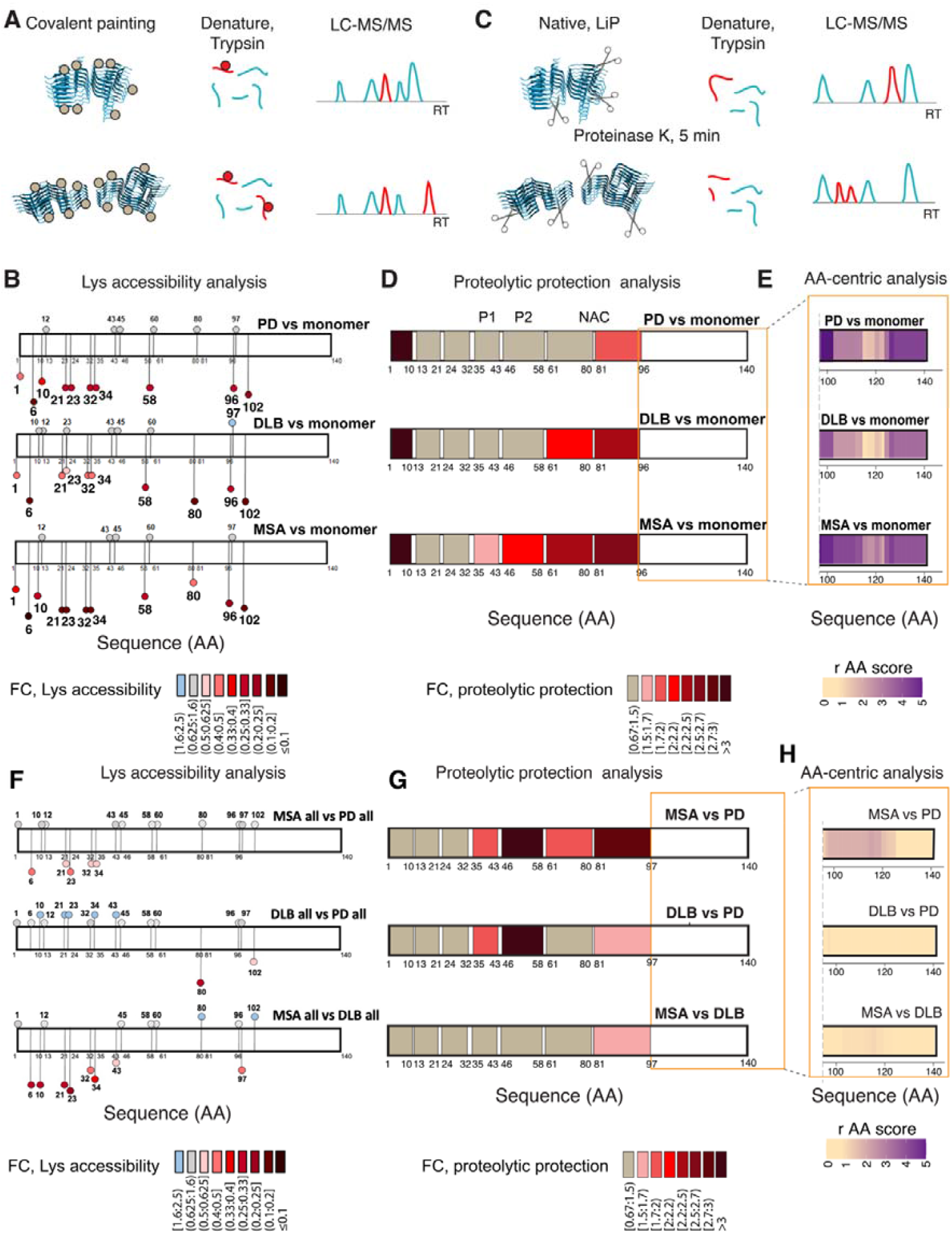
Structural differences between pure PMCA-amplified αSyn fibrillar polymorphs from PD, DLB, and MSA patients. **(A)** Schematic of covalent painting experiment. First, protein surfaces are covalently painted by NHS-biotin. Subsequently, samples are denatured and digested by trypsin for bottom-up MS analysis. **(B)** Covalent painting-based Lys accessibility analysis of PD, DLB, and MSA derived αSyn fibrillar polymorphs vs αSyn monomer. The color scale shows fold changes (FC) of lysine accessibility vs monomer for all lysine residues within αSyn primary structure; darker hues indicate decreased accessibility (n=4 patients for PD, and n=3 patients each for DLB and MSA). **(C)** Schematic of LiP-MS experiment. First, proteins or proteomes are digested in native conditions by Proteinase K. Next, samples are denatured and digested by trypsin to generate shorter peptides optimal for bottom-up MS analysis. **(D)** LiP-MS-based proteolytic protection analysis for PD, DLB, and MSA derived αSyn fibrils vs αSyn monomer. The color scale shows the fold change of proteolytic protection vs monomer along αSyn primary structure; darker hues show increased protection (n=3 patients per disease, n=4 technical replicates per sample). **(E)** LiP-MS-based amino acid-centric analysis of proteolytic patterns in αSyn C-terminal moiety of PD, DLB, and MSA derived αSyn fibrils vs αSyn monomer. The color scale shows the r score, a measure of the change in protease susceptibility per amino acid residue, plotted along the αSyn C terminal primary structure (n=3 patients per disease, n=4 technical replicates per sample). The same patient-derived fibrillar polymorphs were analysed in (B), (D), and (E), except for one additional PD patient (Supplementary Figure 2) in (B). **(F-H)** Direct comparison of the structural features of the PD, DLB, and MSA fibrillar polymorphs. **(F)** Lysine accessibility to biotinylation, **(G)** LiP-MS-based proteolytic protection analysis. The color scale shows the fold change of proteolytic protection between the strains along the αSyn sequence; red hues show increased protection, blue hues show increased accessibility, in each case in the first strain of the pair (n=3 patients per disease, n=3 (F) and n=4 (in G and H) technical replicates per sample). **(H)** LiP-MS-based amino acid-centric analysis of proteolytic patterns in αSyn C-terminal moiety of PD, DLB, and MSA derived αSyn fibrils, compared to each other. The color scale shows the r score, a measure of the change in protease susceptibility per amino acid, plotted along the αSyn C-terminal primary structure (n=3 patients per disease, n=4 technical replicates per sample). Hashed circle representations are used when the signal passes Fold Change cut-off but does not meet the p-value statistical significance cut-off.

**Figure 2.**
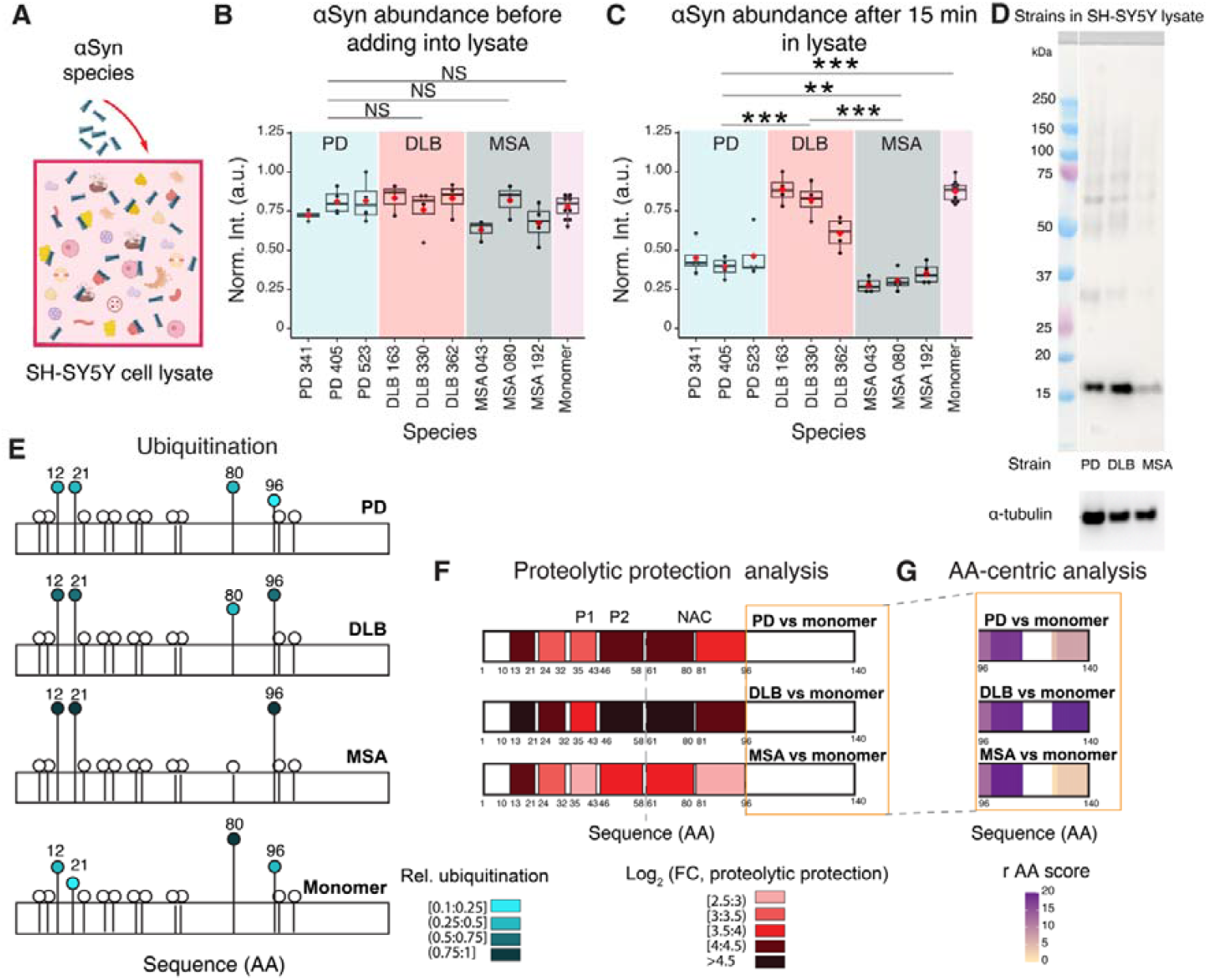
Structural differences, differential post-translational modification, and stability of PMCA-amplified αSyn fibrillar polymorphs from PD, DLB and MSA patients in a cellular lysate. **(A)** Schematic of the experimental setup to probe patient-derived αSyn fibrillar polymorphs structure and stability in a cellular context. Equal amounts of PMCA-amplified PD, DLB or MSA αSyn fibrils, or monomer, were added to separate aliquots of SH-SY5Y lysate. **(B, C)** The plots show mass spectrometric intensities of αSyn preparations before **(B)** and 15 minutes after **(C)** spike-in to the cell lysate. Boxplots represent individual patients, where the black line indicates the median, and the red dot shows the mean value. Individual replicates are depicted as black points. The significance of differential stability of the fibrillar polymorphs is indicated as follows: NS p-val≥0.5, *p-val<0.05, **p-val <0.01, ***p-val < 0.001 **(D)** Western blot analysis of αSyn (anti-αSyn antibody 5G4 (against aa 47-52)) after the same amount of αSyn (1.5 µg) of each strain was incubated for 15 min in cell lysate. SDS-PAGE was done using MES buffer. α-tubulin served as a loading control on the same blot. **(E)** Ubiquitination fingerprints of the indicated αSyn preparations after 15 min in cell lysates (n=3 patients per disease, n=4 technical replicates per sample) or monomer (n=12 technical replicates). All ubiquitination sites are shown, and the four detected sites are numbered (K12, K21, K80, and K96). The color indicates normalized intensity of the ubiquitinated peptides. Intensities are scaled separately for each ubiquitination site, with the maximum change between conformations in each case set as 1. Darker hues indicate a higher relative degree of ubiquitination. White color indicates that no ubiquitination was detected. **(F)** LiP-MS-based proteolytic protection analysis for PD, DLB, and MSA derived αSyn fibrils vs αSyn monomer in cellular lysate. The color scale shows the fold change of proteolytic protection vs monomer along the αSyn sequence; darker hues show increased protection (n=3 patients per disease, n=4 technical replicates per sample). **(G)** LiP-MS-based amino acid-centric analysis of proteolytic patterns in αSyn C-terminal moiety of PD, DLB, and MSA derived αSyn fibrils vs αSyn monomer. The color scale shows the r score, a measure of the change in protease susceptibility per amino acid, plotted along the αSyn C-terminal primary structure (n=3 patients per disease, n=4 technical replicates per sample).

First, we evaluated the structure of amplified fibrils relative to those in original patient brain material. The fibrillar nature of all PMCA-generated αSyn polymorphs was demonstrated by transmission electron microscopy (Supplementary Figure 1A). Their limited proteolysis profiles analysed by gel-electrophoresis show disease polymorph-specific patterns (Supplementary Figure 1B). In previous work, we have also shown this for amplified, *de novo* generated ‘ribbon’ and ‘fibril’ polymorphs^28,36^ as additional controls^36^. Amplified fibrils probed with the amyloid binder, Thioflavin T (ThT), showed disease-specific fluorescence signals (Supplementary Figure 1C), with trends matching the ThT intensities recorded for the patient brain homogenates used for amplification (Supplementary Figure 1D). Indeed, the DLB (n=3) patient brain homogenates and fibrils amplified from these homogenates showed the strongest ThT fluorescence signal. MSA (n=4) patient brain homogenates and fibrils amplified from these homogenates showed the lowest ThT fluorescence signal while their PD (n=4) counterparts exhibited intermediate ThT fluorescence signal. No significant increase in ThT fluorescence was observed for brain homogenates from control individuals (n=7).

Next, we analysed structural differences between the polymorphs by covalent molecular painting MS^39^ and LiP-MS. These methods are based on orthogonal principles and thus yield complementary information. Covalent biotinylation of Lys (K) residues exposed to the solvent under native conditions using NHS-biotin, or covalent painting, followed by proteolytic digestion (with trypsin and GluC) under denaturing conditions allows mapping and quantitative comparison of the accessibility of K residues in distinct biotinylated fibrillar polymorphs using classical bottom-up MS (Figure 1A). We first compared the exposure of K residues to the solvent in each fibrillar polymorph (PD, n=4; DLB, n=3; MSA n=3 patients) relative to monomeric αSyn. Molecular painting revealed protection of a subset of K residues in amplified fibrils (Figure 1B, red shades, Supplementary Data 1). Residues M1, K6, K21, K23, K32, K34, K58, K96, K102 were less exposed to the solvent in all fibrillar polymorphs compared to monomers. However, the protection level of K residues (i.e., the average fold change (FC) in biotinylated peptides between fibrillar polymorphs as compared to monomeric αSyn) in fibrillar polymorphs derived from different synucleinopathies differed significantly. Indeed, K10, 21, 23, 32, 34 showed a disease-specific accessibility pattern, with the highest and the lowest protection levels in the MSA- and DLB-derived polymorphs, respectively. This reflects structural differences within this region between MSA- and DLB-derived polymorphs. Moreover, K80 exposure to the solvent was higher in polymorphs derived from PD cases as compared to those originating from DLB and MSA patients. Assessment of the exposure of K residues in fibrillar polymorphs from individual patients (Supplementary Figure 2A) confirmed polymorph-specific biotinylation footprints, in particular the differential accessibility of K10, K21, K32, K34, and K80. It is worth noting that individual patient-specific features were also observed for K23 in PD and DLB; K43 in MSA; K45 in PD, DLB and MSA and K80 in PD and MSA (Supplementary Figure 2A).

Next, we assessed the structural features of disease-specific polymorphs with LiP-MS, which captures differences in susceptibility of a protein region to a sequence-unspecific protease such as proteinase K (PK). The LiP-MS workflow (Figure 1C) includes two digestion steps (versus one step in classical bottom-up proteomics). First, proteins or proteomes in the native state are subjected to a short (limited) proteolysis by PK. Second, proteins are denatured and proteolyzed to completion with trypsin. Structure-specific proteolytic fingerprints consisting of fully tryptic, semi-tryptic and non-tryptic peptides are generated in this workflow.

We performed LiP-MS-based proteolytic protection analysis of fully tryptic peptides for each fibrillar polymorph (PD, n=3; DLB, n=3; MSA n=3 patients) and for monomeric αSyn. An increase in peptide intensity in this setup reflects the protection of the corresponding protein region from PK cleavages. The protection pattern for each strain was plotted along the sequence of αSyn in the form of a structural barcode (Figure 1D, Supplementary Data 2). As expected, all fibrillar polymorphs exhibited increased protection compared to monomeric αSyn^34^. Furthermore, fibrillar polymorphs derived from PD, DLB, and MSA patients exhibited distinct protection patterns. On average, fibrillar polymorphs derived from PD cases showed the lowest protection in the amino acid stretch spanning residues 61-95, also named non-amyloid-β component region (NAC region), as compared to polymorphs derived from DLB and MSA patients. The latter polymorph exhibited the highest protection level in the NAC and pre-NAC regions, in agreement with the results obtained via the molecular painting strategy. The amino acid stretch spanning residues 1-10 was strongly protected in all polymorphs. These disease-specific structural barcodes could also be observed in fibrillar polymorphs from individual patients, with some inter-individual variability (Supplementary Figure 2B).

Our LiP-MS protection analysis is not suited for the αSyn C-terminal domain (spanning residues 103-140) because of the absence of trypsin cleavage sites. To circumvent this limitation and obtain structural information on the αSyn C-terminal domain, we analysed all PK-generated peptides (fully and semi tryptic). We scored structural differences between the C –termini of αSyn fibrillar polymorphs and monomer at the single amino-acid residue level^44^, using multiple overlapping peptides (n=87 peptides). This analysis yields fold changes (abs(log2(FC)) and their significance (-log10(p value)) for each amino acid residue, and we have previously demonstrated that it captures differences between de novo αSyn fibrils assembled under different experimental conditions^45^. This analysis revealed differences in the C-terminal stretches spanning residues 97-114 and 124-140 of αSyn fibrillar polymorphs and the monomeric form of the protein. No such differences were observed for the central portion of the αSyn C-terminal domain spanning residues 115-123 (Figure 1E, Supplementary Figure 2C). We conclude from these observations that the structure of the αSyn C-terminal domain in fibrillar polymorphs derived from patients is distinct from that of the monomeric form of the protein.

We next used the data generated by our approaches to directly compare the structures of fibrils of all three disease strains to each other. Covalent painting probes lysine accessibility and is therefore perfectly suited to detect changes in the lysine-rich αSyn N-terminus, while LiP-MS exhibits higher sensitivity in detecting changes in protease susceptibility in the pre-NAC and NAC regions of the protein. Indeed, the two techniques provided complementary results. Covalent painting revealed that the DLB strain was the most accessible in the N terminus, while the MSA strain was the least accessible (Figure 1F). Based on LiP-MS, the NAC and pre-NAC regions were more protected in the MSA compared to PD strains (Figure 1G). Also, DLB-derived fibrils were more protected than PD, both in the NAC region based on covalent painting (residue 80) and LiP-MS analysis, and in the pre-NAC region based on LiP-MS (Figure 1F, G). The C-terminal stretch spanning residues 97-117 showed a structural difference between MSA and PD strains; this was less pronounced between the MSA and DLB strains, and we observed no significant difference in C-terminal structure between the PD and DLB strains (Figure 1H).

Taken together, both molecular painting and LiP-MS analyses demonstrate structural differences between PD, DLB, and MSA patient-derived αSyn polymorphs. The NAC region and the 10 N-terminal amino acid residues exhibited the lowest solvent exposure level in all fibrillar polymorphs compared to the monomeric form of the protein. We identified structural differences characteristic for each synucleinopathy in the rest of the αSyn N-terminal domain as well, with the DLB strain being the most solvent-exposed in this region, and in the pre-NAC regions P1 and P2. Our results suggest that the αSyn C-terminal domain is differently structured in MSA patient-derived αSyn polymorphs compared to PD and DLB. Overall, we observe that αSyn exhibits the highest resistance to proteolysis in MSA patient-derived αSyn polymorphs.

### Stability and structural differences between patient-derived αSyn polymorphs in a cellular lysate

We next assessed the stability and structural characteristics of patient brain-derived αSyn fibrillar polymorphs in a complex cellular milieu. The stability of the different polymorphs was assessed by classic proteomic analysis measuring protein abundances before and 15 min after spiking an SH-SY5Y neuroblastoma cell lysate with a constant amount of monomeric or patient brain-derived fibrillar αSyn polymorphs (Figure 2A-C). Cells were harvested without trypsin and cell lysates were generated mechanically using a pellet pestle on ice in the absence of detergent (Methods). Lysis efficiency was evaluated by electron microscopy as well as based on PK accessibility of organellar proteomes (Supplementary Figure 3).

**Figure 3.**
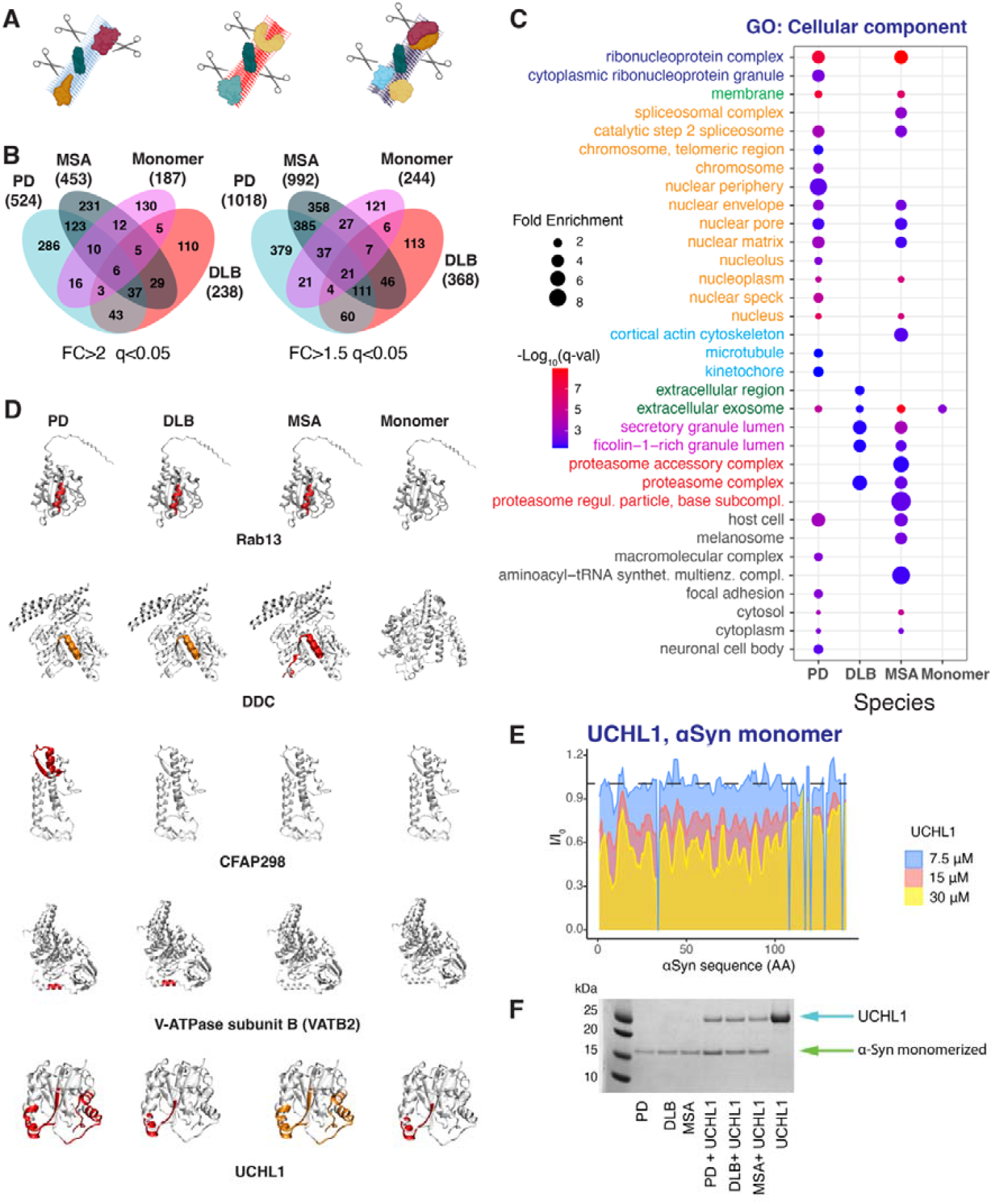
Specific interactomes of disease-derived αSyn fibrillar polymorphs. **(A)** Schematic of the experimental strategy to reveal αSyn fibrillar polymorphs-specific interactomes. First, PD, DLB, and MSA-derived αSyn fibrils or monomer were added to the SH-SY5Y cell lysate for 15 min. Then, LiP-MS and global structural analysis revealed direct and indirect interactors of the spiked-in species. **(B)** The Venn diagram shows the overlap between proteins that show changes in protease susceptibility upon spike-in of the different disease-derived αSyn fibrillar polymorphs or αSyn monomer; two significance cut-offs are shown (left: FC>2, q-val<0.05; right: FC>1.5 q-val<0.05). **(C)** Functional enrichment analysis (GO cellular component) for the set of proteins subject to changes upon spike-in of the different αSyn species. All significant enrichments are shown (q-val<0.05). **(D)** Structural models showing LiP-MS hits mapped onto experimental or αSyn predicted structures of proteins peptides (red (FC>2, q-val<0.05) or orange (FC>1.5 q-val<0.05)) subject to changes upon spike-in of the different αSyn fibrillar polymorphs or monomer. The structures used were: Rab13 (AF-P51153-F1-model), DDC (AF_P20711-F1-model), CFAP298 (AF-C9JX57-F1-model), VATB2 (AF-P21281-F1-model), and UCHL1 (2etl). (E) NMR validation of interaction between UCHL1 and αSyn monomer. Intensity ratios of αSyn peaks in the presence (I) and absence (I_0_) of UCHL1 are shown. 2D [^15^N,^1^H] HMQC NMR spectra of 15 µM ^15^N labelled αSyn were recorded in the absence or presence of increasing quantities of purified UCHL1 in PBS (Supplementary Figure 17). Peak positions and peak intensities were extracted from 2D [^15^N,^1^H] HMQC NMR spectra. (**F**) Coomassie stained SDS-PAGE of αSyn strains and UCHL1 co-sedimentation. The first three lanes correspond to the pellets of PD, DLB and MSA strains only. The next three lanes correspond to the pellets of PD, DLB, and MSA strains pre-incubated with UCHL1. The last lane corresponds to pure UCHL1. All the samples were incubated in 8M urea for 60h to disassemble αSyn fibrils.

The patient-derived fibrillar αSyn polymorphs exhibited differential stability in the SH-SY5Y lysate, since their relative abundance changed upon incubation in the lysate (Figure 2B, C). Specifically, αSyn fibrillar polymorphs derived from PD and MSA patients declined in abundance relative to the DLB fibrillar polymorph and αSyn monomers over 15 min in the lysate. We further validated this disease-specific stability using Western blotting (Figure 2D). Interestingly, this pattern was barely affected upon inhibition of serine and cysteine proteases (Supplementary Figure 4).

**Figure 4.**
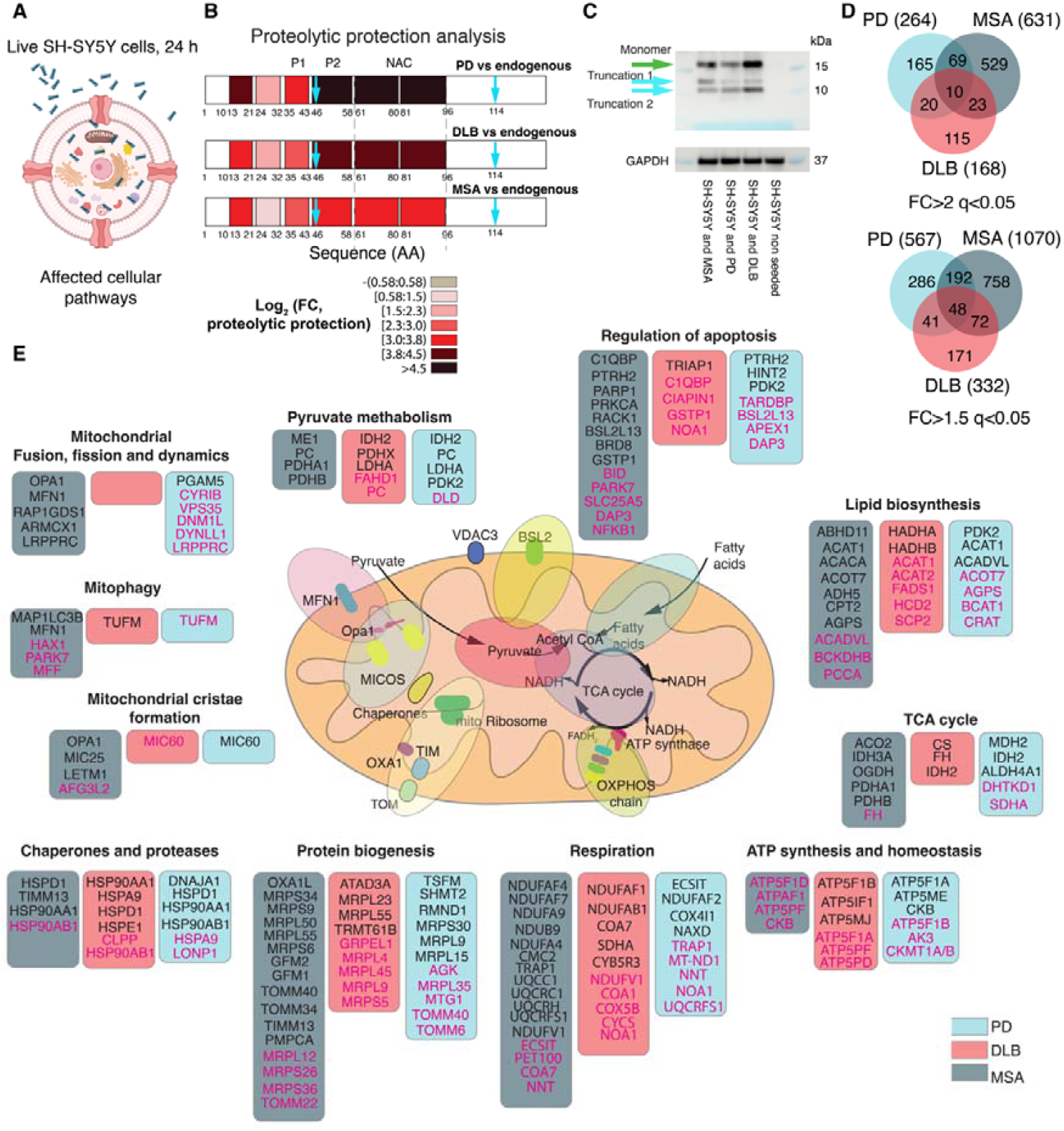
Specific cellular response to disease-derived αSyn fibrillar polymorphs uptake. **(A)** Schematic representation of the experimental strategy to identify cellular pathways affected by different disease-derived αSyn fibrillar polymorphs. PD, DLB, or MSA derived αSyn fibrillar polymorphs were added to living SH-SY5Y cells for 24h. The cells were next washed and lysed in native conditions and LiP-MS was performed to identify fibril polymorph-specific changes in the cellular proteome. **(B)** LiP-MS-based proteolytic protection analysis for αSyn in cells infected with PD, DLB, and MSA fibrillar polymorphs vs endogenous αSyn in control uninfected cells. The color scale shows fold changes in proteolytic protection vs endogenous αSyn along the αSyn primary structure; darker hues show increased protection (n=3 patients per disease, n=3 independent cell infections per sample). Blue arrows indicate sites with evidence of cleavage by endogenous proteases. **(C)** Western blot against αSyn using a mixture of two antibodies (5G4 (aa 47-52) and 42/α-Syn (aa 91-99)) on lysates of live SH-SY5Y cells that were incubated with 250 nM of each αSyn strain for 24h. Green arrow indicates the monomer. Cyan arrows indicate truncated forms. SDS-PAGE was done using MES buffer. GAPDH served as a loading control using GA1R anti-GAPDH antibodies on the same blot. **(D)** The Venn diagram shows the overlap of proteins subject to structural changes upon cell infection with the different disease-derived αSyn fibrillar polymorphs; two cut-offs of significance are shown (top panel: FC>2, q-val<0.05; bottom panel: FC>1.5 q-val<0.05). **(E)** Proteins involved in multiple mitochondrial processes respond to cellular invasion by PD, DLB, and MSA-derived αSyn fibrils. The diagram shows proteins annotated to the mitochondrial processes where at least one peptide is subject to changes upon uptake of the indicated αSyn fibrillar polymorph. Hits are colored (boxes) according to the fibrillar polymorph eliciting the response (PD, cyan, DLB, red, MSA, grey). Hits are also colored (text) based on the cut-off of significance (black: FC>2, q-val<0.05; magenta: FC>1.5, q-val<0.05).

Our MS data showed differential ubiquitination patterns for amino acid residues K12, K21, K80, and K96 for the different αSyn fibrillar polymorphs (Figure 2E, Supplementary Figures 5-9, Supplementary Data 3), which may suggest that differential ubiquitination is responsible for their differential clearance in SH-SY5Y cell lysate. We note however that this analysis does not reveal the fraction of αSyn that is ubiquitinated. Western blot analysis of the αSyn polymorphs after spiking into lysate did not show species that we could definitively attribute to ubiquitination (Supplementary Figure 10).

**Figure 5.**
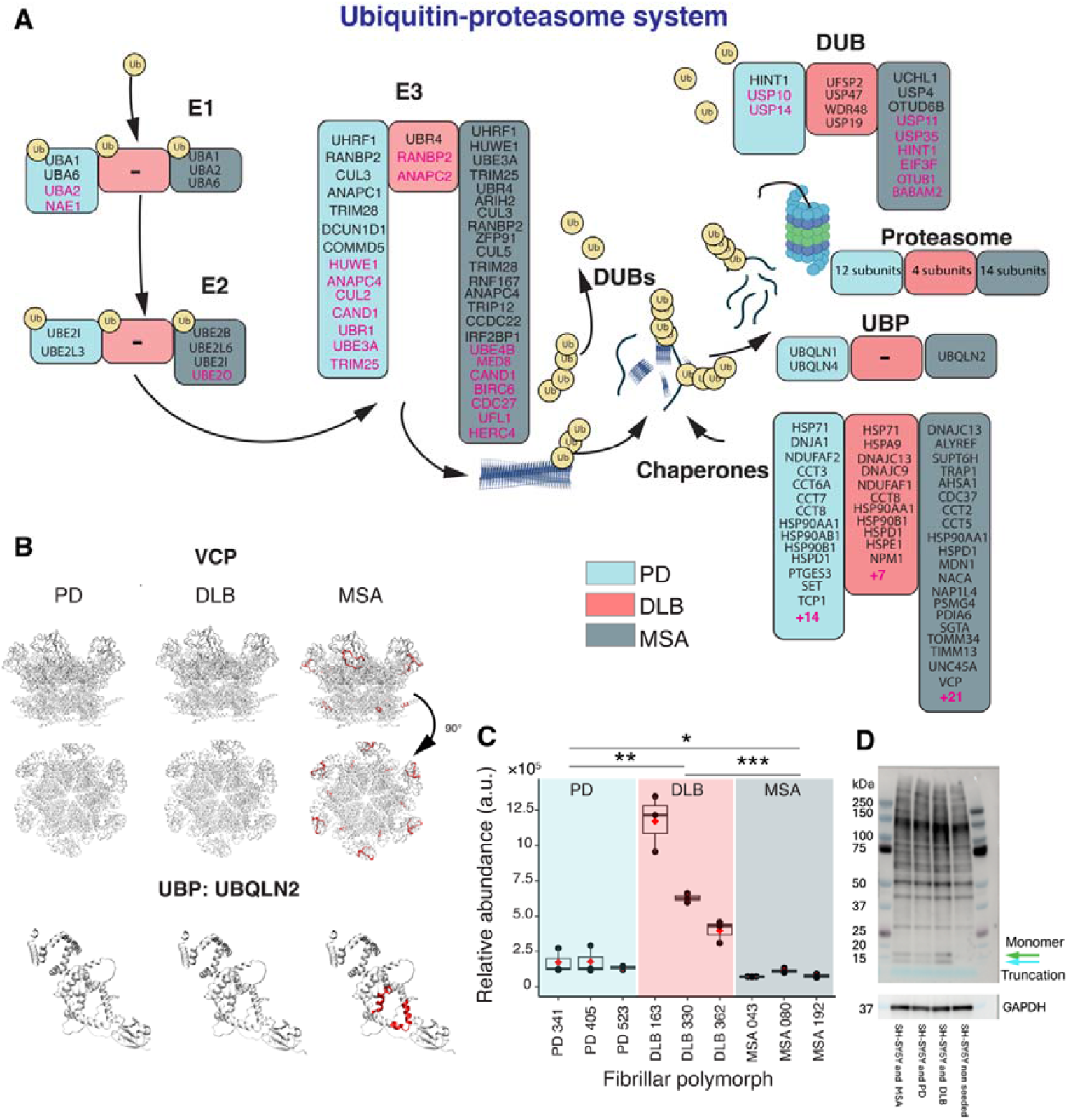
Specific degradation of disease-derived αSyn fibrillar polymorphs. **(A)** The diagram shows proteins annotated to the ubiquitin-proteasomal pathway where at least one peptide is subject to changes upon uptake of the indicated αSyn fibrillar polymorph. Hits are colored (boxes) according to the fibrillar polymorph eliciting the response (PD, cyan, DLB, red, MSA, grey). Hits are also colored (text) based on the cut-off of significance (black: FC>2, q-val<0.05; magenta: FC>1.5, q-val<0.05). **(B)** Structural models mapping the LiP-MS hit peptides (red (FC>2, q-val<0.05)) of VCP and UBQLN2 that change upon uptake of the different αSyn fibrillar polymorphs. The PDB structures used were: 7vcs (VCP, experimental), and AF-Q9UHD9-F1-model (UBQLN2, predicted). **(C)** The plots show the relative quantity of αSyn measured by mass spectrometry 24h after infection of cells with each disease derived fibrillar polymorph. Boxplots represent individual patients, where the black line indicates the median, and the red dot shows the mean value. Individual replicates are depicted as black points. The significance of differences between the fibrillar polymorphs derived from PD, DLB, and MSA is indicated as follows: *p-val<0.05, **p-val <0.01, ***p-val < 0.001. **(D)** Western blot against αSyn using a mixture of two antibodies (5G4 (aa 47-52) and 42/α- Synuclein (aa 91-99)) after living SH-SY5Y cells were incubated with 250 nM of each αSyn strain for 24h. The green arrow indicates αSyn monomer. SDS-PAGE was done using MOPS buffer. GAPDH served as a loading control using GA1R anti-GAPDH antibodies on the same blot.

In the next step, to more deeply characterize endogenous proteolysis, we assessed the cleavage pattern of αSyn strains in cell lysate (Supplementary Figure 11). To this end, we analyzed semi-tryptic peptides in the tryptic control datasets. Since these data are collected in the absence of PK treatment, these peptides identify cleavages from endogenous proteases. We detected multiple such cleavages for all three strains (PD, DLB, MSA; Supplementary Figure 11). Interestingly, the N-terminal end of αSyn (1-7 aa) was significantly less cleaved for MSA fibrils compared to DLB, while the C-terminal end (128-140 aa) was significantly more cleaved in fibrils of the MSA strain.

Next, we used LiP-MS to assess structural differences between patient-derived αSyn fibrillar polymorphs within the context of the SH-SY5Y lysate. Fully and semi-tryptic peptides from αSyn monomers and fibrillar polymorphs clustered separately. In addition, with few exceptions, peptides from fibrillar polymorphs derived from distinct synucleinopathies clustered together (Supplementary Figure 12). This analysis indicates that the structural differences among αSyn fibrillar polymorphs derived from patients with distinct synucleinopathies are also apparent in a complex cell lysate. All patient-derived fibrillar polymorphs exhibited overall increased protection compared to monomeric αSyn in the cell lysate, with the DLB-derived polymorph being most protected (Figure 2F, Supplementary Data 4), and amino acids 13-21 showing the highest protection level in all polymorphs. The amino acid stretch spanning residues 24-96 exhibited protection patterns characteristic of each synucleinopathy. The disease-specific protection barcodes we report were apparent in fibrillar polymorphs from individual patients with limited inter-individual variability (Supplementary Figure 13). Notably, the proteolytic protection patterns of fibrillar αSyn polymorphs in SH-SY5Y cell lysate differed from that in buffer, and they showed overall increased protection in lysate compared to buffer for all protein regions analyzed. The latter observation is not necessarily due to an altered ubiquitination pattern, as no protection of the corresponding fully tryptic peptides was apparent in the tryptic control data (Supplementary Figure 14).

Higher resolution (amino acid-level) analysis of all peptides that were detected in all replicates (n= 24 peptides) revealed polymorph-specific protection patterns also within αSyn C-terminal domain, spanning residues 128-140 (Figure 2G). This reflects conformational differences characteristic of each synucleinopathy within this region and/or differential interactions of this region with proteins within the SH-SY5Y cell lysate.

Next, we performed a direct comparison of proteolytic fingerprints of the different disease-derived fibrils, which showed that MSA fibrils were less protected in the pre-NAC and NAC region (aa 46-96) than PD fibrils. We observed a similar trend when we compared MSA and DLB fibrils, with the MSA strain being less protected across aa 24-96. Comparison of DLB and PD fibrils showed higher protection in specific regions of DLB fibrils (aa 21-43 of the N terminus, the pre-NAC, and aa 81-96 in the second half of the NAC region). We observed differential susceptibility to PK in αSyn C-terminus (region spanning residues 128-140), in particular for DLB vs PD and MSA vs DLB (Supplementary Figure 15).

Altogether, our results demonstrate that αSyn fibrillar polymorphs derived from PD, DLB and MSA patient brain homogenates exhibit different and polymorph-specific stability, ubiquitination, C- and N-terminal cleavages, and structural fingerprints in cell lysates.

### Different interactomes for αSyn fibrillar polymorphs derived from distinct synucleinopathies

The structural differences we report between patient-derived αSyn fibrillar polymorphs in a complex milieu, e.g., cell lysates, may in part reflect specific interactions between αSyn and partner proteins. Therefore, we analysed changes in SH-SY5Y proteome accessibility by LiP-MS upon spiking the cell lysate with monomeric or patient-derived fibrillar polymorphs individually for 15 min (Figure 3A). We recently demonstrated that cellular proteins exhibiting changes in protease-susceptibility represent candidate direct physical interactors of the added forms of αSyn^46^.

We quantified approximately 73 000 LiP peptides corresponding to 5766 proteins through our LiP-MS workflow for each spiked-in form of αSyn (Supplementary Data 5). We measured changes in protease susceptibility (FC>2, q-val<0.05) in 524, 238, 453, and 187 potential protein interactors of PD, DLB, MSA patient-derived αSyn fibrillar polymorphs and monomeric αSyn, respectively. Although there was overlap between potential interactors of patient-derived αSyn fibrillar polymorphs, strikingly, about half of the potential interactors of each disease-derived polymorph were unique (Figure 3B). These patterns also held true when we used a less stringent significance cut-off for changes in protease susceptibility (FC>1.5, q-val<0.05), allowing less prominent candidate protein interactors to be considered (Figure 3B).

Gene Ontology (GO) enrichment analysis of the candidate interactors of monomeric and patient-derived αSyn fibrillar polymorphs (Figure 3C) showed overlap in a single enriched cellular component term ‘Extracellular exosome’. Terms related to protein-RNA interactions, such as ‘ribonucleoprotein complex’, ‘spliceosomal complex’, and ‘cytoplasmic ribonucleoprotein granule’, as well as to ‘cytoskeleton’ (‘cortical actin cytoskeleton’, ‘cytoskeleton’, and ‘microtubules’) and ‘nucleus’ were enriched among interactors of αSyn fibrillar polymorphs derived from PD and MSA, but not from DLB patients or monomers. Terms pertaining to the ‘proteasome’ and ‘secretory vesicles’ were enriched among interactors of DLB and MSA-derived αSyn fibrillar polymorphs. Other terms were enriched for different combinations of αSyn patient-derived polymorphs.

We further identified domains within selected hits of interest that are most affected upon spike-in of monomeric and patient-derived αSyn fibrillar polymorphs and highlighted these domains in the corresponding protein structures (Figure 3D). Here, we highlight several representative and disease-relevant examples. A LiP peptide from Rab13, a known modulator of αSyn aggregation^47^, exhibited altered protection in the presence of all patient-derived αSyn fibrillar polymorphs but not αSyn monomer. This suggests a direct interaction between Rab13 and patient-derived αSyn fibrillar polymorphs. A similar behavior was observed for aromatic-L-amino-acid decarboxylase (DDC), a crucial enzyme in dopamine synthesis. In contrast, in the N-terminal region (amino acid stretch 8-33) of the cilia and flagella-associated protein CFAP298, proteolytic susceptibility changed significantly only in the presence of the PD-derived αSyn fibrillar polymorph. A region of the V-ATPase (subunit B), responsible for synaptic vesicle and lysosome acidification^48-50^, was differentially protected in the presence of PD and DLB patient-derived αSyn fibrillar polymorphs.

PD, DLB, and MSA patient-derived αSyn fibrillar polymorphs exhibited differential stability and ubiquitination patterns in cell lysates (Figure 2). Furthermore, different proteasome-related terms were found enriched in the GO analysis of candidate interactors of DLB and MSA patient-derived αSyn polymorphs (Figure 3C). This observation, together with the differential stability of polymorphs in lysates and their ubiquitination, prompted us to inspect the UPS and protein degradation pathways for potential interactors of distinct αSyn fibrillar polymorphs (Supplementary Figure 16; Supplementary Table 1, Methods). We selected proteins with functions annotated in Uniprot with the keywords ‘Ubiquitin conjugation’ (biological process) and ‘Chaperones’ (molecular function). We identified several E3 ligases and DUBs as potential interactors; these classes of proteins are of substantial interest as they are substrate-specific, crucial for protein degradation, and potentially druggable. Among the 272 proteins associated with the Ubiquitin conjugation process (these include E1, E2, E3 ligases, DUBs, and Ubiquitin binding proteins), 10, 4, and 7 E3 ligases and 3, 3, and 5 DUBs exhibited altered proteolytic protection in the presence of PD, DLB and MSA patient-derived αSyn fibrillar polymorphs, respectively (FC>2, q-val<0.05) (Supplementary Table 1). The proteolytic protection pattern of the E3 ligase HUWE1 changed in the presence of all αSyn patient-derived fibrillar polymorphs. That of UBE3A and UHRF1 changed only in the presence of PD- and MSA-patient derived αSyn fibrillar polymorphs. The proteolytic protection pattern of the Ubiquitin C-terminal Hydrolase L1 (UCHL1), a DUB proposed to be a genetic risk factor for Parkinson’s disease^51^, changed in the presence of all patient-derived fibrillar polymorphs and monomeric αSyn (Figure 3D). We confirmed the direct interaction of ^15^N labeled monomeric αSyn and unlabeled UCHL1 using NMR and further showed that the interaction involves the N-terminus and NAC region of αSyn (Figure 3E, Supplementary Figure 17). NMR detected no interaction between αSyn monomer and the negative control CHCHD2, which is also a PD-linked protein^52^ but was not a hit in our screen (Supplementary Figure 18), pointing to the specificity of the αSyn interaction with UCHL1. To validate the interaction between αSyn fibrillar polymorphs and UCHL1, we performed a co-sedimentation assay. Purified UCHL1 co-sedimented with all three αSyn fibrillar polymorphs after incubation for 15 min, while it remained in the supernatant in the absence of αSyn fibrils (Figure 3F), providing further evidence for a physical interaction.

In total, 19, 9, and 18 molecular chaperones exhibited protease susceptibility changes (FC>2, q-val <0.05, Supplementary Data 6) in the presence of PD, DLB, and MSA patient-derived fibrillar polymorphs, respectively. Fewer (4) molecular chaperones were detected as putative interactors for monomeric αSyn. Finally, since we saw evidence of endogenous proteolysis of spiked-in fibrils, we asked whether there were proteases among the candidate interactors of αSyn fibrils. Multiple proteins annotated as proteases in Uniprot (KeyWord Molecular function ‘Protease’) showed protease susceptibility changes, with 14 proteases responding to PD, 7 to DLB, and 12 to MSA strains (FC>2, q-val<0.05, Supplementary Table 2). These proteases are interesting candidates for follow-up study.

Altogether, we identified proteins that interact differentially with patient-derived fibrillar polymorphs and monomeric αSyn with sufficient resolution to map candidate interaction sites to the protein structure. The differential interactomes we identified may account for differential tropism of distinct disease-associated αSyn fibrillar polymorphs and for some of the phenotypes associated with distinct synucleinopathies.

### Specific cellular responses to uptake of patient-derived αSyn fibrillar polymorphs

We previously showed that LiP-MS can detect multiple functional molecular events^33,35^. In particular, LiP-MS captures differences in protease susceptibility upon protein aggregation, post-translational modification, protein-protein interactions, allosteric changes, protein-small molecule interactions and changes in enzymatic activity, and as a consequence can detect alterations in cellular pathways^33,35^. To assess cellular changes elicited by the different patient-derived fibrillar polymorphs, we made use of a well-established model of endocytic uptake of fragmented αSyn fibrils, which mimics prion-like infection of healthy cells^7,11,53^. We incubated adherent SH-SY5Y neuroblastoma cells with fragmented fibrils from each patient-derived fibrillar polymorph for 24h (Figure 4A; n=3 patients, 3 independent infections for each PD, DLB or MSA patient-derived fibrillar polymorph), lysed the cells in native conditions to preserve protein structures and interactions, and performed the LiP-MS workflow. Comparison of the profiles obtained from fibril-treated and untreated cells after normalization for changes in protein abundance (Methods) then allowed the identification of structural changes in the intracellular proteome, and thus cellular pathways, induced by each polymorph. Notably, such changes may be triggered irrespective of αSyn location within the cell (i.e., in cytosol, inside of cellular compartments or bound to the lipid membrane).

We first examined the structure of αSyn fibrils themselves in this system. The patient-derived αSyn fibrils exhibited strong protection toward proteolysis compared to endogenous αSyn from untreated cells 24 hours after uptake, with differential levels of protection for each disease (Figure 4B, Supplementary Data 7). Since some αSyn fibrils might be hidden from PK in cellular compartments (e.g., endosomes) in this seeded live cell model, we directly assessed αSyn accessibility to PK by quantifying by MS the loss of intensity of fully tryptic peptides under LiP conditions. This analysis revealed that αSyn is about 62% (PD), 70% (DLB) and 76% (MSA) cleaved at its most accessible regions, indicating that it is largely accessible to PK (Supplementary Figure 19). Further, PD, DLB, and MSA fibrils were cleaved by endogenous proteolysis after residue E114 and E46 in the C- and N-terminal regions, respectively, as detected by the quantification of semi-tryptic peptides in the absence of added PK. Notably, E46 corresponds to a PD-linked familial mutation (E46K)^54^ and E114 was recently reported as a major cleavage site during αSyn cellular processing^55^. αSyn truncations were also detected using Western blot analysis of seeded cell extracts after fibril depolymerization by urea (Figure 4C, Supplementary Figure 20). Interestingly, we observed two truncated forms in cells infected with all the strains, but MSA fibrils showed different relative levels of these two forms compared to PD and DLB, consistent with our observation that the αSyn C-terminus is structured differently in the MSA strain (Figure 1). We note that the high molecular weight (HMW) bands in this seeded cell model are largely due to background staining (Supplementary Figure 20), an effect that was much weaker in our lysate spike-in experiments, where αSyn was added at approximately an 8-fold higher concentration compared to the cell seeding model, resulting in a higher signal to noise ratio. To demonstrate this, we have spiked monomeric αSyn into cell lysates in parallel to the fibrils (Supplementary Figure 21). These data show that, in the spike-in context, the HMW background signal is much weaker than in the cell seeding model.

At the proteome level, we observed changes in protease susceptibility (FC>2, q-val<0.05) in 264, 168 and 631 proteins upon SH-SY5Y cell infection with PD, DLB, MSA patient-derived αSyn fibrillar polymorphs, respectively (Figure 4D). These changes likely represent both the effects of fibrillar uptake and of fibrils bound to the cell membrane. The lists of altered proteins common to all disease-associated polymorphs or specific to each synucleinopathy are provided as a resource (Supplementary Data 8), together with information on the specific altered protein regions. This list includes mitochondrial proteins (Figure 4E) such as ATP synthase subunits, Opa1, the MICOS complex, VDAC3, the TOM complex and many other proteins performing crucial molecular functions. These results are in line with genetic and environmental evidence suggesting that mitochondrial dysfunction plays an important role in synucleinopathies^56-60^.

We next focused our analysis on the UPS, to better understand the differential turnover of αSyn fibrillar polymorphs within living cells. The Parkin-ubiquitin proteasomal system was indeed over-represented in a pathway enrichment analysis of proteins for which we detected protease susceptibility changes upon fibril uptake (WikiPathways, Supplementary Figure 22). SH-SY5Y cell invasion (24h) by αSyn fibrillar polymorphs led to changes in protease susceptibility of numerous proteins involved in the UPS, as identified by at least one LiP specific peptide with FC>2, q-val <0.05 (black) or FC>1.5, q-val <0.05 (magenta) (Figure 5A). These included E1, E2 and E3 ubiquitin and SUMO-protein ligases, molecular chaperones, ubiquitin binding proteins (UBP), deubiquitinating enzymes (DUBs), proteasomal subunits and endogenous proteases. As expected from our global analysis, changes common to PD, DLB and MSA patient-derived fibrils or specific to each polymorph could be observed.

Our LiP-MS workflow allowed the mass spectrometric identification of 149 molecular chaperones in total (Supplementary Data 9). Changes in protease susceptibility of family members of the molecular chaperones Hsp70, Hsp40 and Hsp90, mitochondrial chaperones, CCT and the AAA+ ATPase Valosin-containing protein (VCP) were detected. VCP is of particular interest as its protease susceptibility changes only upon uptake of MSA patient-derived fibrils, has been proposed to exhibit disaggregase activity, and was demonstrated to reduce the size of tau fibrils and promote their proteasomal degradation^61^. The two most protected LiP-MS peptides we identified in VCP are highlighted in the quaternary structure of the protein (Figure 5B, top row). The peptides spanning amino acid residues 26-45 and 701-708 from VCP N-terminal domain and D2 region are important for VCP interaction with co-factors and ATPase activity, respectively.

In total, 295 ubiquitin conjugating proteins were detected and identified through the LiP-MS workflow (Supplementary Data 10). Changes in the protection patterns of multiple E3 ligases or subunits of E3 ligase complexes in response to uptake of PD patient-derived fibrils (UHRF1, RANBP2, CUL3, TRIM25, TRIM28, HUWE1, UBE3A), MSA patient-derived fibrils (UHRF1, HUWE1, ARIH2, UBE3A, CUL3, TRIM25, TRIM28, UBR4, ZFP91, RNF167, CUL5) and DLB patient-derived fibrils (UBR4, RANBP2, ANAPC2), were observed (Figure 5A, FC>1.5, q-val <0.05). A similar picture was seen for numerous DUBs (Figure 5A). Unlike what we observed upon addition of all patient-derived fibrillar polymorphs and monomeric αSyn to cell lysates (Figure 3), the protection pattern of UCHL1 was significantly affected only upon uptake of MSA patient-derived fibrils (Supplementary Figure 23).

Finally, the protection patterns of ubiquitin binding proteins, believed to recognize and deliver ubiquitinated proteins to the proteasome for degradation^62,63^, exhibited polymorph-dependent changes (Figure 5A, right). Uptake of PD patient-derived fibrillar αSyn triggered changes in UBQLN1 and UBQLN4 profiles (FC>1.5, q-val <0.05); while only that of UBQLN2 was affected by the MSA polymorph (Figure 5B) and none by the DLB fibrils (Figure 5A, right). Considering simply the number of proteins for which the protection profile changes in response to αSyn fibrillar polymorphs internalization, our analysis suggests a generally weaker response of the UPS to DLB as compared to PD or MSA patients-derived fibrillar αSyn. A similar trend was observed for proteasomal subunits (Figure 5A) and for proteins related to autophagy (Supplementary Figure 24), which also play an important role in αSyn clearance^64^. Thus, both the UPS and autophagic degradation systems exhibit changes of their component proteins characteristic of each patient-derived αSyn fibrillar polymorph.

Intrigued by the endogenous cleavage we observed within the αSyn fibrillar polymorph C-terminus (Figure 4B) we examined our LiP-MS hits for the presence of proteases that have been previously proposed to cleave the protein (Calpain, Neurosin, CtsB, CtsL)^65^. The complete list of proteases responding to the PD, DLB, and MSA strain is given in Supplementary Table 3. Among these, we identified Calpain as a LiP-MS hit for MSA fibril-infected cells, and CtsL for DLB fibril-infected cells, suggesting that these proteases may be interesting candidates in follow-up studies.

Since we had observed differential stability of the disease-derived fibrillar polymorphs in cell lysates, we next asked whether we could observe differences in the cellular levels of αSyn accumulation 24h after uptake of PD, DLB and MSA patient-derived fibrillar αSyn (Figure 5C). Consistent with our findings that DLB patient-derived fibrillar αSyn exhibits the highest stability in cell lysates (Figure 2C), classical MS-based protein abundance quantification, using 10 αSyn peptides, showed striking accumulation of αSyn in cells exposed to DLB-derived αSyn fibrils as compared to PD and MSA-derived counterparts. This finding was further supported by Western blot analysis (Figures 4C, 5D). Our data thus show that SH-SY5Y cells respond differentially to distinct patient-derived fibrillar polymorphs, with polymorph-associated disparity within cellular protein degradation systems as well as differential stability of the PD, DLB, and MSA patient-derived fibrils. Finally, we repeated the experiment with synthetic fibrils (PFFs) self-assembled in PBS at pH 7.4 (Supplementary Figure 25A), conditions that are often used to generate *in vitro* PFFs for cellular studies^66^. After normalization for protein abundance, we detected 93 proteins for which PK susceptibility changes upon treatment with PFFs (250 nM) for 24h relative to control (Supplementary Figure 25B). We observed fewer proteins structurally responding to PFFs in comparison to the disease strains. Some were common with the hits of at least one disease strain (52 proteins), and some were PFF-specific (41 proteins) hits (Supplementary Figure 25C). Interestingly, proteins involved in the ubiquitin-proteasomal system also responded structurally to synthetic PFFs, as we had observed for the disease strains (e.g., TRIM28, UCHL1, UHRF1) (Supplementary Figure 25D, E). However, the E3 ligases UBE3A, TRIM25, HUWE1, UBR4 and the chaperone VCP, which all showed structural changes in response to fibrils of at least one disease strain, did not significantly respond to PFFs. Proteins responding to PFFs were enriched for the ‘mRNA processing’ pathway, as in the DLB and MSA strains. Our data suggest that commonly used PFFs share some features of disease-specific fibrils in terms of the downstream affected pathways in SH-SY5Y cells, but that the responses they elicit are not identical.

### iPSC-derived cortical neurons differential response to patient-derived fibrillar polymorphs uptake

To further strengthen our findings and test their relevance in a more disease relevant context, we exposed iPSC-derived cortical neurons to the PD, DLB, and MSA strains and used LiP-MS to characterize their response. For these experiments, we differentiated a pool of iPSCs from three healthy individuals into cortical neurons and confirmed differentiation by fluorescence microscopy of neuron-specific markers (Supplementary Figure 26). Neurons were incubated with PD, DLB, and MSA αSyn fibrillar polymorphs for 24h, and then processed for LiP-MS (Methods). We performed 2-3 independent seedings for each patient, yielding 8-9 replicates per disease and 12 replicates of control (no exposure to fibrils, equal volume of PBS) neurons (Figure 6).

**Figure 6.**
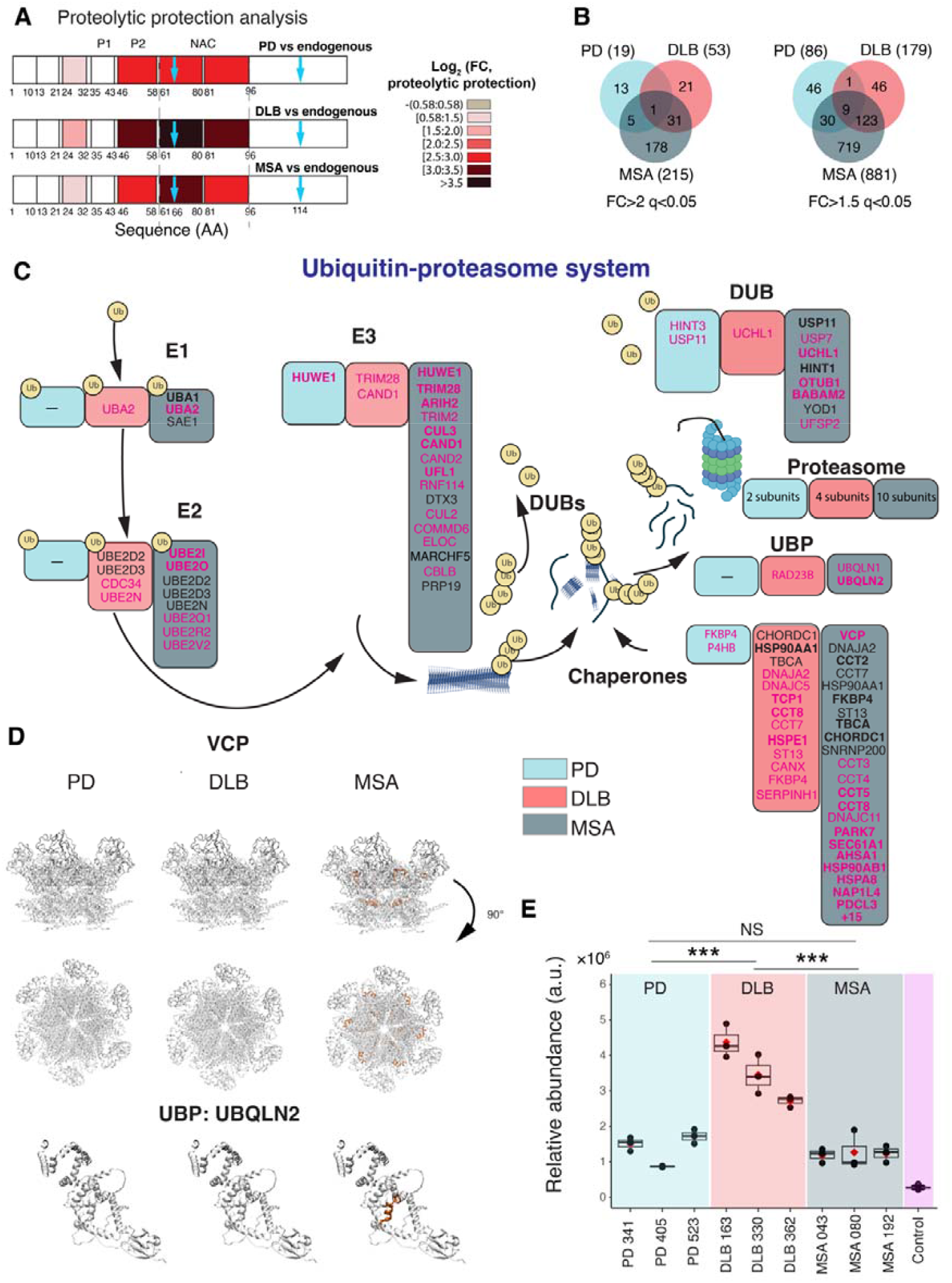
iPSCs-derived cortical neurons respond to disease-derived αSyn fibrillar polymorphs in a strain-specific manner. **(A)** LiP-MS-based proteolytic protection analysis for αSyn in iPSCs-derived cortical neurons infected with PD, DLB, and MSA fibrillar polymorphs vs endogenous αSyn in control uninfected neurons. The color scale shows fold changes in proteolytic protection vs endogenous αSyn along the αSyn primary structure; darker hues show increased protection (n=3 patients per disease, n=2-3 independent cell infections per sample). Blue arrows indicate endogenous protease cleavages. **(B)** The Venn diagram shows the overlap of proteins that undergo structural changes upon neuron infection with the different disease-derived αSyn fibrillar polymorphs; two cut-offs of significance are shown (left: FC>2, q-val<0.05; right: FC>1.5, q-val<0.05). **(C)** The diagram shows proteins annotated to the ubiquitin-proteasomal pathway where at least one peptide is subject to changes upon uptake of the indicated αSyn fibrillar polymorph. Hits are colored (boxes) according to the fibrillar polymorph eliciting the response (PD, cyan, DLB, red, MSA, grey). Hits are also colored (text) based on the significance cut-off (black: FC>2, q-val<0.05; magenta: FC>1.5, q-val<0.05). Hits are highlighted in bold if they were also observed in the corresponding experiment on SH-SY5Y cells for corresponding strain (Figure 5). **(D)** Structural models mapping the LiP-MS hit peptides (orange (FC>1.5, q val<0.05)) of VCP and UBQLN2 that change upon uptake of the different αSyn fibrillar polymorphs in neurons. The PDB structures used were: 7vcs (VCP, experimental), and AF-Q9UHD9-F1-model (UBQLN2, predicted). **(E)** The plots show the relative quantity of αSyn measured by mass spectrometry 24h after infection of cells with each disease-derived fibrillar polymorph. Boxplots represent individual patients, where the black line indicates the median, and the red dot shows the mean value. Individual replicates are depicted as black points. The significance of differences between the fibrillar polymorphs derived from PD, DLB, and MSA is indicated as follows: *p-val<0.05, **p-val <0.01, ***p-val < 0.001.

As in SH-SY5Y cells, αSyn fibrillar strains exhibited higher proteolytic resistance compared to endogenous αSyn in infected neurons, with strain-specific digestion patterns (Figure 6 A, Supplementary Data 11). We also detected cleavages from endogenous proteases, after residue 65 and 114. The UPS system was again enriched for neurons seeded with DLB and MSA strains (Supplementary tables 4-5, WikiPathways ‘proteasomal degradation’ and ‘parkin-ubiquitin proteasomal system pathway’), and we observed, as in SH-SY5Y cells, that these hits span several classes of proteins within the UPS, including DUBs, chaperones, and E1, E2 and E3 ligases (Figure 6B, C). We observed many similarities with the hits we identified in SH-SY5Y cells. This includes the E3 ligases HUWE1, Trim28, ARIH2, Cul3, CAND1, and UFL1 and the DUB UCHL1. Further, we recapitulated the MSA-specific response of VCP and UBQLN2 also in neurons and observed changes in the identical region in UBQLN2 (aa 218 to 232) in both systems (Figure 6D). Nevertheless, there were also neuron-specific effects such as enrichment of multiple pathways related to cholesterol metabolism for DLB and MSA fibrils (Supplementary Tables 4 and 5).

Importantly, we again observed a relatively higher accumulation of αSyn in neurons exposed to the DLB strain, compared to the other two strains, further corroborating this finding in a third model. Overall, we detected fewer proteins with structural changes in neurons compared to the SH-SY5Y cells (Figure 6B), but again the MSA strain showed the highest number of hits. Taken together, our findings in neurons largely support those we report using SH-SY5Y cells. They strongly suggest a disease strain-specific response to fibril infection and that the DLB strain accumulates to a higher extent compared to the PD and MSA strains.

### Global structural proteome alterations in patient brains afflicted by PD, DLB, and MSA

To determine to what extent the changes we observed upon exposure of SH-SY5Y cells and iPSC-derived cortical neurons to different patient-derived fibrillar polymorphs reflect those occurring in patient brains, we performed the LiP-MS workflow we described above directly on the brain homogenates we used to generate disease-specific αSyn polymorphs by PMCA (Figure 7). We compared the cingulate gyrus of PD and DLB patients and the cerebellum of MSA patients to the corresponding regions from control donors (n=3 each).

**Figure 7.**
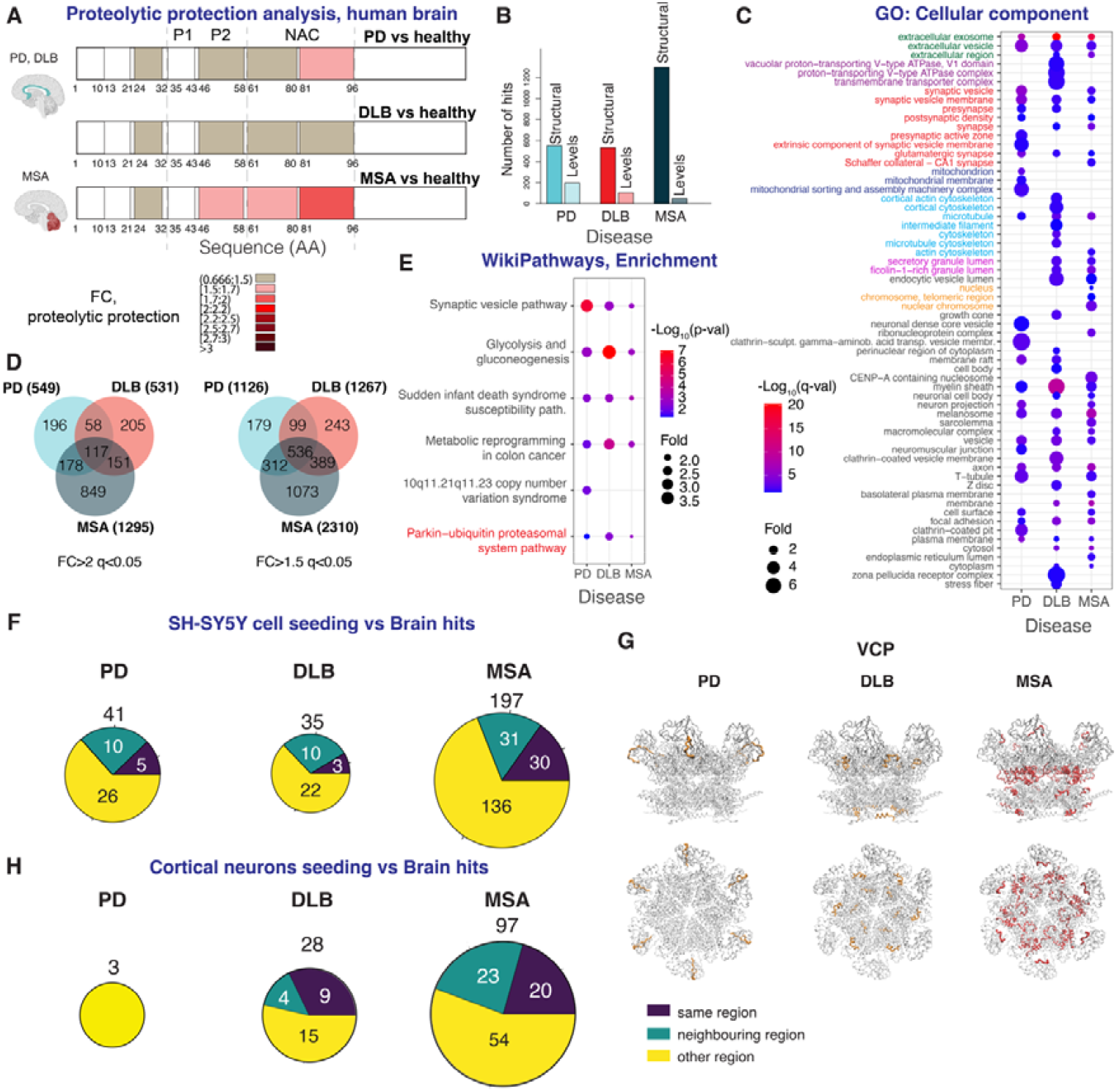
Common and disease-specific alterations in proteomes of PD, DLB, and MSA brain homogenates. **(A)** LiP-MS-based proteolytic protection analysis of αSyn in brain homogenates of patients suffering from PD, DLB, and MSA vs healthy control individuals. The color scale shows fold change of proteolytic protection (FC), along the αSyn primary structure, of PD, DLB, and MSA derived αSyn fibrillar polymorphs vs monomer; darker hues show increased protection (n=3 patients per disease and control, n=4 technical replicates per sample). Cingulate gyrus was analyzed for PD and DLB and controls, while Cerebellum was analyzed for MSA and corresponding control patients. **(B)** The plot shows the number of proteins affected by structural changes and which abundance varies for PD, DLB and MSA brain homogenates, relative to control individuals (q-val<0.05, FC>2). **(C)** Functional enrichment analysis (GO cellular component) for the set of proteins that fulfill the criteria above in PD, DLB or MSA afflicted brains. All significant enrichments are shown (q-val<0.05). **(D)** The Venn diagram shows the overlap of proteins that exhibit changes in different synucleinopathies; two cut-offs of significance are shown (left: FC>2, q-val<0.05; right: FC>1.5 q-val<0.05). **(E)** Pathways enrichment analysis (WikiPathways) for the set of proteins that show changes in patient brains afflicted by synucleinopathies. Displayed here are the top five enriched pathways for each synucleinopathy (p-val<0.05). Complete list of enriched pathways is in Supplementary Tables 6-8. **(F)** The plot shows the overlap of LiP-MS hits (i.e., proteins that show changes) between the SH-SY5Y cell seeding experiment and the comparison of brain proteomes (q-val< 0.05, FC >2). **(G)** Structural models mapping the significantly altered LiP-MS peptides (red) of VCP in the indicated disease brain homogenates (red (FC>2, q-val<0.05) or orange (FC>1.5 q-val<0.05)). VCP structure PDBID 7vcs was used. **(H)** The plot shows the overlap of LiP-MS hits (i.e., proteins that show changes) between the experiment on seeded cortical neurons and the experiment on brain proteomes (q-val< 0.05, FC >2).

First, we assessed the structural features of αSyn in native brain homogenates. This analysis provides insights into the structure of pathogenic, monomeric and any other αSyn form simultaneously present in the lysate. The αSyn amino acid stretch spanning residues 81-96 from the NAC region exhibited increased protection from proteolysis in PD compared to control patient brain homogenates. The αSyn portion exhibiting increased protection was larger in MSA patient brain homogenates, spanning the entire NAC and pre-NAC regions (P2) (FC>1.5, q-value<0.05) (Figure 7A, Supplementary Data 12). We did not observe a significant increase in αSyn protection in DLB patient brain homogenates. αSyn in brain homogenates from individual patients exhibited overall protection patterns characteristic of each synucleinopathy, with exceptions (Supplementary Figure 27). Notably, patient PD341 exhibiting the highest αSyn aggregate load based on previous quantifications^31^ showed a protection pattern similar to that of MSA patients, while patient PD405 who had the lowest pathogenic αSyn load^32^ showed a pattern similar to DLB patients (Supplementary Figure 27) albeit with a 1.4 fold increased protection within the NAC peptide 81-96, just below our cut-off set to 1.5.

We also identified changes in protease susceptibility in 549, 531 and 1295 proteins in PD, DLB and MSA patient brain homogenates (FC>2, q-val<0.05), respectively, relative to control donors, after adjusting for protein abundance (Figure 7B). We detected many fewer changes in the abundances of proteins between control and diseased patient brain homogenates, supporting our previous observations in cerebrospinal fluids^67^.

Functional enrichment analysis (GO: Cellular Component) of proteins showing changes in protease susceptibility identified terms enriched for all synucleinopathies (e.g., ‘extracellular exosomes’, ‘synaptic vesicles’ and ‘synapses’) but also disease-specific terms (e.g., ‘mitochondria-related’ in PD, ‘V-type ATPase’ and ‘cytoskeleton’ in DLB, the ‘nucleus’ in MSA) (Figure 7C, FC>2, q-val<0.05). In total, 117 proteins showed changes in protease susceptibility in all three synucleinopathies (Figure 6D, FC>2, q-val<0.05, Supplementary Data 13). These common hits were enriched for proteins associated with ‘Parkinsonism’ and ‘Epilepsy’, based on UniProt Keyword Disease enrichment analysis. Interestingly, a pathway enrichment analysis of proteins with altered protease susceptibility (FC>2, q-val<0.05; Figure 7E and Supplementary Table 6-8, Wikipathways) again revealed that the ‘Parkin-Ubiquitin proteasomal system’ pathway was enriched in all diseased relative to control brains.

We compared the lists of proteins that show changes in protease susceptibility in PD, DLB and MSA patients brain homogenates to that in SH-SY5Y cells exposed to patient-derived fibrillar polymorphs. This identified 41, 35 and 197 overlapping proteins for PD, DLB and MSA, respectively (Figure 7F, FC>2, q-val <0.05, Supplementary Data 13). Mapping the LiP peptides to the respective protein structures showed that a significant number (15, 13 and 61, for PD, DLB and MSA, respectively) were located in the same or neighboring regions in cells exposed to patient-derived fibrillar polymorphs and in the brain lysates they originated from (Figure 6F). We observed when examining UPS proteins in particular, overlapping protease susceptibility changes in E3 ligases (2 for PD, 1 for DLB, and 8 for MSA) and DUBs (1 for PD and 4 for MSA) (Supplementary Table 9, FC>1.5, q-val<0.05). Notably, multiple proteins exhibited altered protease susceptibility in a manner specific to the disease across both sample types. For instance, changes in protease susceptibility in UBR4 were observed in DLB and MSA but not in PD brain homogenates, and in cells exposed to DLB- and MSA-but not PD-patient derived fibrillar polymorphs. Similarly, changes in protease susceptibility of VCP were observed in MSA only patient brain homogenates, and in cells exposed to fibrils derived from those same homogenates (Figure 7G, q-val <0.05, FC >2). Lowering the threshold stringency (q-val<0.05, FC>1.5) led to detecting changes in susceptibility to protease in VCP in samples originated from PD and DLB patients. Several other UPS proteins, including HUWE1, TRIM25, TRIM28, RanBP2, HINT1, CUL3, CAND1 exhibited changes in both experimental models.

Similarly, we identified 3, 28 and 97 proteins that show protease susceptibility changes in both PD, DLB and MSA patient brain homogenates and in iPSCs-derived cortical neurons exposed to patient-derived fibrillar polymorphs (Figure 7H, Supplementary Data 14, q-val<0.05, FC>2). Interestingly, for 9 of the 28 shared hits for DLB (YWHAZ, HSP90A, JPT1, Creatine kinase B-type (CKB), PRDX1, Transketolase (TKT), TPI1, TPM3, and TPM4), the changing regions of the hit proteins were the same in both models. In the case of MSA, 20 proteins out of the 97 shared hits (among them, YWHAQ, YWHAZ, COPE, CPNE1, ENOG, EPN1, FBL, FKBP4, FLNA) showed the change in the same protein region. The three shared hits for PD were Clathrin heavy chain 1 (CLH1), serine/threonine-protein kinase PRKDC, and Receptor-type tyrosine-protein phosphatase zeta (PTPRZ). Importantly, the E3 ligases HUWE1, TRIM28, TRIM2, CUL2, CAND1 and DUBs UCHL1, OTUB1, and HINT1 and the AAA-ATPase VCP were altered in both neurons exposed to the disease fibrils and in brain homogenates derived from patients with the corresponding disease (q-val<0.05, FC>1.5). Taken together, our data reveal differential functional interaction networks for both pathogenic αSyn in PD-MSA- and DLB-patient brain homogenates and fibrillar polymorphs derived from those homogenates after uptake by SH-SY5Y cells and neurons. The differences we report point to disease-relevant pathways in distinct synucleinopathies.

### Different clearance of αSyn fibrillar polymorphs derived from distinct synucleinopathies

We show above that αSyn structures within brain homogenates from patients affected by PD, DLB or MSA exhibit structural properties characteristic of each synucleinopathy. We further demonstrated in vitro and in SH-SY5Y cell lysates that αSyn fibrillar polymorphs derived from those homogenates present distinct structural features. Our data also suggest disease-specific ubiquitination patterns and degradation pathways for αSyn fibrillar polymorphs derived from different synucleinopathies in cell lysates and in infected cells. Finally, we identified members of the UPS among fibrillar αSyn polymorph interactors in cell lysates and infected cells and showed them to be polymorph specific. All together our data suggest that αSyn fibrillar polymorph degradation is differentially regulated.

We therefore asked whether modulation of the identified degradation pathways affects cellular levels of pathogenic, aggregated αSyn. We focused on a subset of proteins that showed altered protease susceptibility in the presence of at least one αSyn fibrillar polymorph, in at least two out of our four orthogonal assays, and where polymorph-specificity was conserved, namely, TRIM25, UBE3A, HUWE1, UBR4 and VCP (Figure 4-7, Supplementary Tables 1 and 9). Except for VCP, the proteins we selected are all E3 ligases, and several are known to be involved in neurological disorders. The set of structurally changing proteins upon disease-specific fibril infection includes additional members of pathways associated with these specific E3 ligases (Figure 8A), in particular E1 and E2 ligases known to be upstream of UBE3A^68,69^, HUWE1^70,71^, TRIM25^72,73^, and UBR4^74-76^ (Figure 8A). Our LiP-MS analysis showed that the protease susceptibility of UBE3A, TRIM25, and HUWE1 was specifically influenced by PD- and MSA-derived αSyn fibrillar polymorphs. The protease susceptibility of UBR4 was dependent on fibrillar polymorphs originated from DLB and MSA.

**Figure 8.**
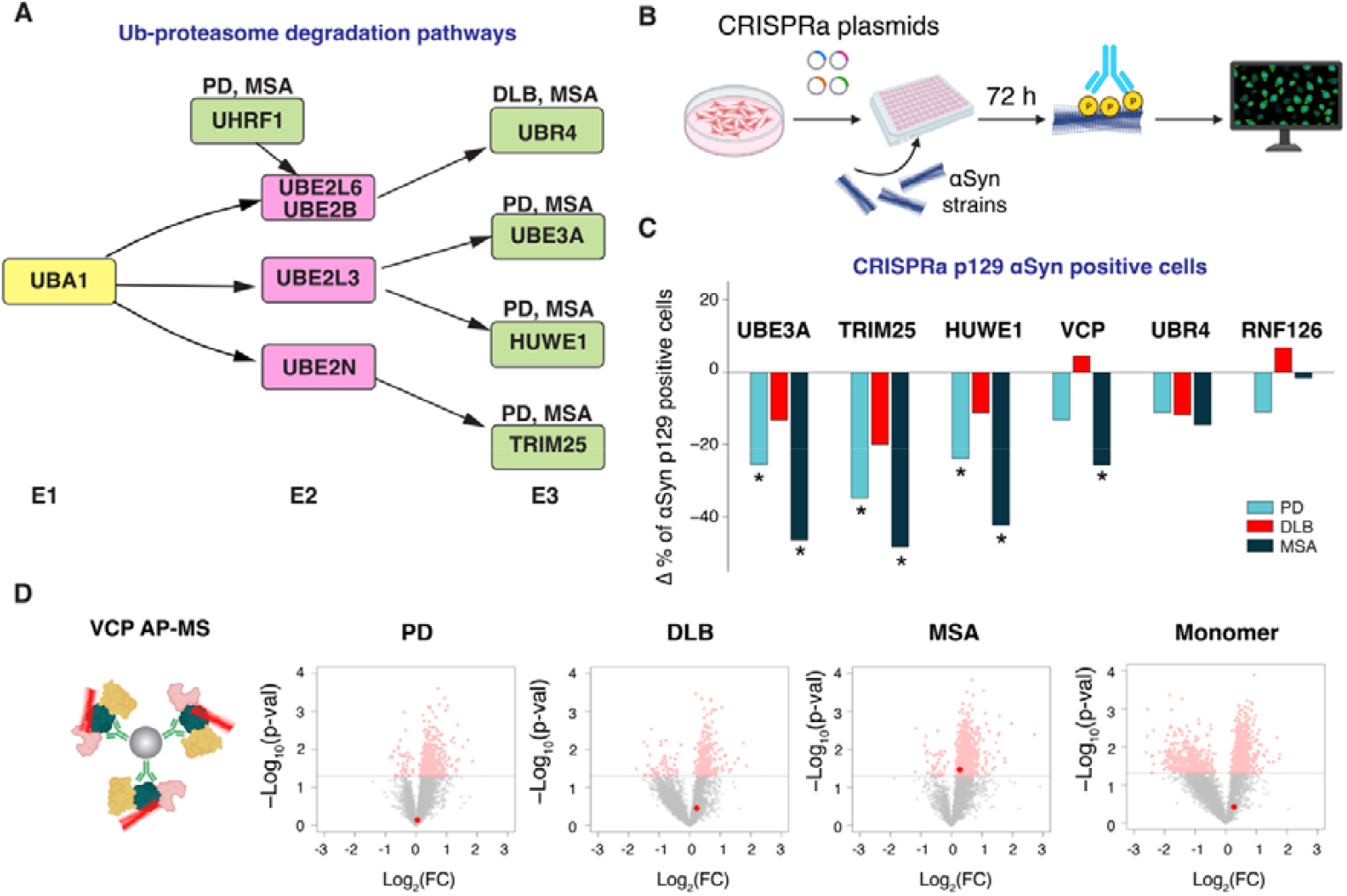
Validation of selected protein hits in disease-derived αSyn fibrillar polymorphs turnover. **(A)** The schematic shows components of cellular degradation pathways (E1 and E2 ligases and ubiquitin binding proteins) we showed to be altered through our experiments of cellular invasion by or cell lysate spike-in with αSyn fibrillar polymorphs and that are known to be upstream of the selected E3 ligases. Proposed disease-specific pathways are indicated, based on known interactions^68-76^. **(B)** Schematic of the CRISPR gene modulation experiment to test the effect of gene overexpression on accumulation of aggregated αSyn. The fraction of cells containing p129 αSyn was quantified upon cell infection with individual fibrillar polymorphs. **(C)** The plot shows the percentage of change of the fraction of p129 αSyn positive cells, upon exposure to the indicated fibrillar polymorph (PD in blue, DLB in red, MSA in grey) and activation of the indicated gene. As a control, non-activated cells infected with the same concentration of the fibrillar polymorphs were used. Asterisks indicate significance (p-val<0.05) relative to control based on a two-tailed t-test. **(D)** AP-MS validation of VCP interaction with the MSA patient-derived fibrillar polymorph. VCP was immunoprecipitated from SH-SY5Y lysate that was pre-incubated with each of the αSyn fibrillar polymorph or the monomer. The volcano plots show proteins that change upon anti-VCP immunoprecipitation versus isotype-specific IgG control; the red dot indicates αSyn and the significance threshold is indicated (p-val< 0.05).

We used a HEK293 cell line overexpressing αSyn^10,77,78^ to determine whether the five selected proteins quantitatively modulate pathogenic αSyn inclusions^10,79,80^ in living cells upon uptake of PD-, DLB- and MSA-derived αSyn fibrils. The expression of the genes encoding each of the selected proteins was upregulated with CRISPR, validated on mRNA and protein level (Supplementary Table 10) and the fraction of cells exhibiting pathogenic phospho S129-αSyn (P-αSyn) deposits was assessed after 72h incubation with patient-derived fibrillar αSyn from each synucleinopathy (Figure 8B, Supplementary Figures 28-30). E3 ligase RNF126 served as a negative control, as it did not exhibit changes in susceptibility to protease in the presence of αSyn fibrillar polymorphs despite being detected by LiP-MS; upregulation of RNF126 showed no significant effect on the fraction of P-αSyn positive cells compared to non-targeting control. Overexpression of TRIM25, UBE3A, and HUWE1 in cells exposed to PD and MSA-derived fibrillar αSyn, significantly reduced the fraction of P-αSyn positive cells as compared to control cells (Figure 8C). In contrast, and in agreement with our LiP-MS data (Figure 5A), the overexpression of the above proteins had no effect on the fraction of P-αSyn positive cells exposed to DLB-derived fibrillar αSyn, compared to control. UBR4 overexpression did not significantly affect the proportion of P-αSyn positive cells in our experimental set-up. Overexpression of VCP, which showed altered protease susceptibility in MSA brain homogenates and upon cell infection by αSyn fibrils derived from those homogenates, significantly reduced the fraction of P-αSyn positive cells after exposure specifically to MSA-derived fibrillar αSyn, compared to control. Affinity purification mass spectrometry (AP-MS) showed that VCP physically interacts with MSA-derived fibrillar αSyn, thus demonstrating the specificity of this interaction (Figure 8D). Taken together, we provide evidence for specific regulation of αSyn fibrillar polymorphs turnover by the cellular degradation machinery and identify various other pathways specific to αSyn fibrils derived from distinct synucleinopathies.

## Discussion

The 3D structure of a protein defines its function and cellular interactome. Amyloids are diverse in their protein composition and are associated with distinct diseases. Recent evidence suggests that polymorphism can result from the aggregation of one given protein into structurally diverse amyloid fibrils. However, the relationship between the structural diversity/polymorphism of amyloids and their cellular interactome, which may account for pathological diversity, is yet unknown. Here, we document the structural features, interaction partners, and cellular effects of αSyn fibrillar polymorphs derived from PD, DLB, and MSA patient brain homogenates. We showed that PD, DLB, and MSA αSyn fibrillar polymorphs differ in structure, interactomes and are differentially processed within the cellular milieu.

For our analysis we have chosen αSyn fibrils amplified directly from native patient brain homogenates. We have previously shown that our amplification procedure works with high fidelity for synthetic fibrils^36^, and importantly, that patient-derived amplified preparations faithfully capture disease-specific effects of the brain homogenates themselves in animal models^32^. This suggests that the structure-pathology relationship of αSyn fibrils is preserved in the amplification process. Further, our data showed disease-specific ThT signals on both amplified fibrils and brain homogenates, and recapitulated previous work showing that ThT emission is lower when bound to fibrils of the MSA vs PD strain^81,82^. The main alternative fibril preparation method is based on Sarkosyl precipitation of the fibrils from the brain. There are as yet no definitive data on whether fractionated αSyn fibrils in the presence of detergent or fibrils amplified from patient brain are the most relevant model of fibrils in vivo. The current literature uses αSyn fibrils prepared with both methods^30-32,81,83-85^. Direct comparison of amplified, Sarkosyl-precipitated, and in vivo αSyn fibrils within human brain tissue remains necessary. Emerging in situ EM techniques, such as cryo-electron tomography of thin patient brain sections, are expected to enable such studies at the required resolution.

We assessed the characteristic interactome and the cellular response to the uptake of structurally distinct, patient specific fibrillar αSyn polymorphs at a proteome-wide scale. We show that αSyn aggregates from patients with distinct synucleinopathies interact with different protein partners, exhibit disease-specific ubiquitination patterns, differential turnover, and differentially regulated cellular pathways in affected cells. Restoring or boosting the function of key protein partners may hold therapeutic potential.

Two complementary MS-based techniques allowed us to identify structural differences between αSyn fibrillar polymorphs derived from PD, DLB, and MSA patient’s brain homogenates, including differences in the αSyn C-terminal domain. This is of particular importance as classical structural biology methods, such as cryo-EM, are poorly suited for probing dynamic domains^36^.

LiP-MS can identify protease susceptibility differences reflecting differences in structure and/or interactions with cellular partners in complex cellular backgrounds such as cell lysates or tissue homogenates^33,35^. We applied LiP-MS to cells exposed to fragmented αSyn fibrillar polymorphs derived from PD, DLB, and MSA patient brain homogenates and identified the cellular response to disease-derived αSyn strain uptake. Remarkably, we observed that αSyn fibrillar polymorphs derived from PD, DLB, and MSA patient brain homogenates differentially affected the mitochondrial proteome, in particular proteins involved in functional maintenance of the organelle, e.g., fusion, fission, dynamics, cristae formation and turnover (Figure 4). This is in agreement with macroscopic changes we reported recently upon the accumulation of pathogenic αSyn in human neurons treated with synthetic αSyn fibrils^86^. The polymorph-dependent differences we observed, in particular those involved in fusion, fission and dynamics, suggest that mitochondrial maintenance is differentially affected by DLB as compared to PD and MSA patient-derived αSyn fibrils. This result is in line with previous evidence suggesting that mitochondrial dysfunction plays an important role in synucleinopathies^56-60,86,87^

We further applied LiP-MS to native brain homogenates and measured the average conformational state of αSyn for PD, DLB, and MSA patients and control individuals. Without applying any enrichment steps, we were able to detect an increased protection of αSyn NAC or NAC and pre-NAC regions for PD and MSA patients, respectively, compared to healthy donors’ brain homogenates. Cells exposed to the different patient-derived fibrillar polymorphs cleared those pathogenic αSyn aggregates to different extents and exhibited distinct responses as assessed by the changes in the proteome we describe. αSyn fibrillar polymorphs derived from MSA and DLB patients were cleared to the highest and lowest extent, correspondingly. Similar results were obtained upon dilution of αSyn fibrillar polymorphs into cell lysates, which suggests that clearance rather than uptake or other upstream cellular events is responsible for differential accumulation of pathogenic αSyn. Furthermore, we validated these results in iPSCs-derived cortical neurons. Interestingly, we observed different responses of members of the UPS, and autophagy processes, with the highest number of alterations in protein structures or interactions for MSA patient-derived fibrillar polymorph in all tested models. Finally, our CRISPR-based modulation of a set of candidate regulators indicated strain-specific and generally stronger effects on αSyn pathological inclusions caused by the PD and MSA strain compared to DLB (Figure 8).

In contrast to SH-SY5Y cells, the iPSCs-derived cortical neurons showed a strong response to MSA and DLB fibrillar polymorphs, while they responded less to PD fibrils at least in terms of the number of altered proteins. This could hint at selective vulnerability of different cell types to different fibrillar polymorphs. Further investigations are needed to test this hypothesis.

The VCP ATPase appears to interact specifically with fibrillar polymorphs derived from brains of MSA patients and showed altered protease susceptibility in brains of patients with the same disease. Whether VCP activity is beneficial or not in neurodegenerative diseases remains poorly understood. VCP mutations were linked to proteinopathies^88^, and VCP is believed to fragment long Tau protein fibrils in the cell^61^. While the fragmentation of fibrillar assemblies may favor proteasomal degradation, it yields shorter fibrils with increased seeding propensity. Thus, fibril fragmentation may be deleterious as we recently showed^89,90^. Further studies are needed to understand how VCP activity affects pathogenic fibrillar polymorphs characteristic of MSA and the role this protein plays in MSA progression.

Pathogenic αSyn within patient brains or fibrillar polymorphs derived from those patients diluted in cell lysates appear either to interact directly with or to trigger changes in several components of the UPS pathway, including the E3 ligases HUWE1, UBE3A, TRIM25, and UBR4. A mutation in TRIM25 was shown to cause early-onset autosomal dominant dementia with amyloid load and parkinsonism^91^. UBE3A is associated with the Angelman Syndrome^92^, and its maternal loss leads to motor deficits and nigrostriatal dysfunction in a Ube3a (m-/p+) mouse model. In line with these findings, upregulation of UBE3A and TRIM25 with a CRISPR approach affected the seeding propensity of PD and MSA patient-derived fibrillar αSyn. UBR4 is highly expressed in the central nervous system and plays an important role in neurogenesis and neuronal signaling^93^. Nonetheless, upregulation of UBR4 expression did not significantly impact p129 αSyn accumulation in our experimental setup despite the changes we observed within this protein in the presence of pathogenic αSyn within DLB and/or MSA patients’ brains or fibrillar polymorphs derived from those patients. HUWE1 is important for neurodevelopment and mitophagy^94^, is strongly expressed in the adult brain, and is enriched in the olfactory bulb, superficial layers of the cortex, hippocampus, and cerebellum. In addition, mutations within HUWE1 are linked to familial idiopathic intellectual disability^95^. Our data suggest that HUWE1 plays an important role in synucleinopathies as it appears involved in PD and MSA patient-derived fibrillar αSyn turnover.

Changes in the susceptibility to proteases of the E3 ligase UHRF1 were also observed in cells and cell lysates exposed to PD and MSA fibrillar polymorphs (Figure 3, 5). UHRF1 plays an important role in neurodevelopment and neurogenesis. Indeed, deletion of UHRF1 in the developing cerebral cortex led to postnatal neurodegeneration^96^. UHRF1 is a nuclear protein bridging DNA methylation and histone modifications^97^. Thus, it could regulate pathogenic αSyn aggregate clearance via epigenetic mechanisms. Indeed, UHRF1 was shown to modulate the methylation of the E2 ligase UBE2L6 promoter^76^, and thereby its expression. It is worth noting that UBE2L6 cooperates with the E3 ligase UBR4 (Figure 8). The changes in the susceptibility to proteases of the ubiquitin ligases UHRF1, UBE2L6, and UBR4 we report may suggest they are involved in clearance pathway specific to pathogenic αSyn aggregates (Figure 8A).

Bridging cellular models with the pathology of neurodegenerative disease is challenging because of the inherent limitations of every cell model and given the timeframe of the experiments (hours, days) compared to authentic disease progression (years). In addition, brain tissues heavily affected by αSyn pathology may not exhibit changes occurring at the early stages of the disease. Nonetheless, we identify alterations that occurred both in cell infection models and in patient brains, allowing prioritization of disease-relevant pathways in follow-up studies.

Altogether, our work demonstrates the structure-pathology relationship in distinct synucleinopathies. It also establishes differential ubiquitination patterns, turnover, interactomes, and downstream effects for pathogenic αSyn within patient brains and for fibrillar polymorphs derived from those patients. We have identified new pathogenic αSyn modulators and present a comprehensive resource of disease-relevant proteins for distinct synucleinopathies. These proteins directly or functionally interact with pathogenic αSyn in a structure-dependent manner. Our results reduce the landscape of potential drug targets to several hundreds of candidate proteins that can be prioritized in follow-up functional studies.

## Materials and methods

### Human brain tissue homogenates

Human brain tissue was obtained post-mortem from patients with PD, MSA and DLB through the Parkinson’s UK tissue bank (Imperial College London, UK). The clinical and neuropathological description of the four PD (PD258, PD341, PD405, PD523), three DLB (DLB163, DLB330, DLB362), and four MSA (MSA043, MSA080, MSA192, MSA363) patients was reported previously^32^. Patient’s brain homogenates were obtained by dilution at 20% (weight:volume) and sonication in Protein Misfolding Cyclic Amplification (PMCA) buffer (150mM KCl, 50mM Tris-HCl pH7.5). Quantification of aggregated αSyn was performed using both the Cisbio fluorescence resonance energy transfer (FRET) assay (Cisbio, France, cat # 6FASYPEG) and immunodetection with the P-129 αSyn antibody (mouse 11A5, provided by Elan Pharmaceuticals, Inc., Dublin, Ireland) as described previously^32^. Brain homogenates were aliquoted, flash frozen in liquid nitrogen, and stored at −80 °C until further use.

### Preparation of αSyn fibrils PMCA-amplified from PD, MSA and DLB brain tissues

PMCA-elongation of patients-derived αSyn seeds was performed as described previously^32,98^. Briefly, brain homogenates were diluted to a final concentration of 2% (W:V) in PMCA buffer (150 mM KCl, 50 mM Tris–HCl, pH 7.5) containing recombinant monomeric αSyn (100 μM) expressed and purified as previously described^28,99,100^. Three cycles of PMCA amplification were performed for each patient in exactly the same conditions as the protocol previously optimized and described^32^. Amplification was monitored by measuring Thioflavin T fluorescence. Cycle 2 to 3 were performed using 1% of the preceding cycle reaction as seeds for PD and DLB cases, 5% for MSA cases and with the same sonication and assembly program. PMCA-amplified reaction of cycle 4, was performed by mixing 40 µL of sonicated cycle 3 PMCA-amplified samples to 760 µL of monomeric αSyn (at 100 µM in 50mM Tris/HCl pH 7.5, 150mM KCl) and elongation was performed in an Eppendorf Thermomixer set at 37°C and 600rpm until completion. PMCA-amplified assemblies were sedimented (50 000g, 30min, 30°C) and monomeric αSyn concentration in the supernatant was determined spectrophotometrically and subtracted from the total recombinant monomeric αSyn added for amplification. Each four-round pelleted PMCA-amplified αSyn sample, from each PD, DLB and MSA sample, was resuspended at 100µM in phosphate-buffered saline (PBS) buffer, before sonication for 20 minutes in a vial sonicator (Vial Tweeter UIS250v, 250 W, 2.4 kHz; Hielscher Ultrasonic, Teltow, Germany), at 75% amplitude with the following sonication cycle: 0.5 sec On/0.5 sec Off. Aliquots were immediately liquid nitrogen frozen and stored at −80°C before use. Morphology of PMCA-derived αSyn assemblies was assessed by negative stain Transmission Electron Microscopy (TEM) imaging and limited proteolysis with proteinase K (Supplementary Figure 1) as described previously^32,101^. Thioflavin T measurements were performed as described previously ^31^.

### Covalent surface painting of αSyn monomers and PMCA-amplified patients’ seeds

#### Covalent biotinylation

Buffer of αSyn samples was changed to 40 mM Hepes/KOH pH 7.5, 75 mM KCl (labeling buffer) using a NAP-5 desalting column containing Sephadex G-25 resin for monomeric αSyn and sedimentation of PMCA-assembled αSyn fibrils by ultracentrifugation (30 min, 50 000 rpm, 20°C, S120AT3 rotor) for resuspension of the pelleted patient’s derived αSyn fibrils in the labeling buffer. Concentration of monomeric αSyn was determined spectrophotometrically in the NAP-5 desalted αSyn fractions and in the supernatant after sedimentation of PMCA-amplified αSyn fibrils using a molar extinction coefficient at 280 nm of 5960 M^-1^.cm^-1^. Concentration of αSyn samples (monomers and Patient’s PMCA-derived fibrils) was adjusted to 100 µM (monomeric aSyn equivalent) with labeling buffer.

αSyn samples (monomeric and PMCA-derived fibrils at 100µM) were labelled with N-hydroxysulfosuccinimide biotin (EZ-Link Sulfo-NHS-Biotin, Thermo Scientific) at a molar ratio of 1:5 (αSyn/NHS-Biotine) during 10 min at room temperature. Stock solution of Sulfo-NHS-Biotin diluted at 100mM in DMSO was further diluted in the labelling buffer. Biotinylation reaction was stopped by addition of 1M Tris HCl pH7.5 to reach 50 mM final concentration. Unbound biotin was removed from to αSyn samples using a NAP-5 desalting column for monomeric αSyn and by ultracentrifugation (30 min, 50 000 rpm, 20 °C, S120AT3 rotor) for PMCA-derived αSyn fibrils. Pelleted fibrils were resuspended in 50µl of proteolysis buffer (50 mM Tris-HCl, pH 7.5) and finally dried using a speed vacuum for further dissociation of αSyn sample fibrils by addition of pure HFIP (hexafluoroisopropanol). After evaporation of HFIP under a hood, dried samples were resuspended in 50µl of ultrapure milliQ water. Unlabeled sample was incubated in the same conditions and used as control. The average incorporation of biotin was between 1 and 3 for all αSyn samples, as determined by MALDI-MS in the same conditions as the one described previously^39^.

#### Covalent painting analysis

After dilution at 25µM in proteolysis buffer, αSyn samples were submitted to a total in-solution digestion using trypsin Gold (Promega) or Glu-C endoprotease sequencing grade (Merck) during 16h under 350rpm agitation with an enzyme/substrate (w/w) ratio of 1/20 and a temperature of 37°C or 25°C for trypsin and Glu-C, respectively. Proteolytic peptide samples were stored at −20°C until use. Proteolytic peptides resulting from trypsin and Glu-C were mixed at a 1:1 ratio (vol:vol) and three technical replicates for each αSyn sample were analysed by nanoLC-MS/MS using a Triple-TOF 4600 mass spectrometer (ABSciex) as previously described^39^. Raw data were converted into mgf data files using the MS Data Converter software (version 1.3) included in the PeakView software (version 1.2, AB Sciex).

Peptides identifications were performed using the Mascot search engine (Matrix Science, London, UK; version 2.4.1) against the human wild-type αSyn sequence, including a decoy database search. Peptides were identified using specific digestion with Trypsin and Glu-C with up to five missed-cleavage and following variable modifications: oxidation of methionine (+15.99 Da); biotinylation of N-ter, Lys, Ser, Thr, Tyr (+226.08 Da). Mass tolerances were set to 40 ppm and 0.05 Da for precursors and fragments respectively. False discovery rate (FDR) was set to 1%.

A list of peptides identified with a peptide mascot score above 20 and in at least two replicates out of three was used to extract the ion intensity of each peptide from the more intense mass-to-charge ratio for each replicate data set. The extracted ion chromatogram (XIC) manager plugin of the PeakView® Software (version 1.2, ABSCIEX) was used for automatic extraction of peak intensities from the chromatograms with a mass tolerance of 0.1 Da and a retention time window of 0.3 min. The accordance between the MASCOT peptide identification and the MS2 spectrum of the extracted ion intensity was checked manually by both detection of signature ions of biotinylation (at m/z=227.08 for biotin and 310.5 for biotinylated Lysine) and comparison of the MS and MS/MS spectra.

Peptide ion intensities were normalized to the total amount of αSyn in each sample, using the non-labeled reference peptide 132-GYQDYEPEA-140 generated repeatedly by Glu-C digestion of αSyn (according to Caroux *et al*.^39^) resulting from a systematic cleavage at E131, absence of cleavage at D135 and E137, and only 5% of cleavage occurrence at E139. Fold change (F_c PMCA-F/M_) between PMCA-derived αSyn fibrils (*PMCA-F*) and monomeric (*M*) αSyn sample, for a given peptide, was calculated as the ratio of the averaged normalized peptide intensity for the PMCA-amplified fibrils to the averaged normalized peptide intensity for the monomers. Statistical significance of the F_c_ was assessed by a non-parametric Kruskal-Wallis test (p-value < 0.050).

### Cell culture and fibril infection

SH-SY5Y neuroblastoma cell line (ATCC) was cultured in T75 tissue culture flasks in DMEM/F-12 Glutamax (#10565018, ThermoFisher Scientific) + 10% FBS, and 1% Penicillin/Streptomycin. For αSyn fibrils infection experiments, cells were grown in 6-well plates to medium confluence. The cells were incubated with 250 nM of αSyn fibrils in culture media for 24h. Treated cells and mock controls were grown in parallel in the same incubator and shared 6 well-plates. After incubation, cells were washed twice with ice-cold PBS, harvested, pelleted down in 15 ml falcon tubes by centrifugation, snap-frozen, and stored at −80°C.

#### Native cell lysis

Pellets of SH-SY5Y cells were resuspended in 200 μl of LiP buffer (100 mM HEPES, 150 mM KCl, 1 mM MgCl_2_, pH 7.4). Next, 10 cycles of 10 douncing steps were performed using a pellet pestle on ice. The sample was cooled on ice for one minute between each cycle of douncing steps. Concentrations of proteins were assessed by bicinchoninic acid assay (BCA) assay. After native lysis and concentration measurements the samples were directly processed by LiP-MS.

### iPSC culture and neuronal differentiation

The derivation of the three healthy control iPSC lines from dermal fibroblasts we use in this study was described previously^102,103^. The three lines are now also deposited and distributed by EBISC under alternative names SFC840-03-03 (STBCi026-A (RRID:CVCL_RB85)), SFC854-03-02(STBCi066-A (RRID:CVCL_RC86)) and SFC856-03-04(STBCi063-A (RRID:CVCL_RC81)). The iPSCs were cultured in Essential 8 medium (#A1517001, Thermo Fisher) on Geltrex (#A1413302, Thermo Fisher) coated 6 well tissue culture plates and passaged as previously described with 0.5 mM EDTA^104^.

Cortical neurons were differentiated following a previously described dual SMAD inhibition protocol^105^, with minor modifications. Stocks of neuronal cells were frozen at day 27 after induction. Neurons were cultured in neuronal maintenance medium (NMM) consisting of 50% Neurobasal medium (#21331-020, ThermoFisher) and 50% DMEM:F12 medium (#1103049, ThermoFisher) supplemented with 0.5x N2 (#17502048, ThermoFisher), 0.5x B27 (#17504044, ThermoFisher), 0.5x NEAA (#11140035, ThermoFisher), 2.5 µg/ml Insulin (#I9278, Sigma), 50 µM Sodium Pyruvate (#S8636, Sigma), 200 µM GlutaMax (#17504044, ThermoFisher) and 50 µM 2-Mercaptoethanol (#31350010, ThermoFisher).

After thawing the neurons NMM was supplemented with 10 µM Rho kinase inhibitor (#MCE-HY-10583, MedChemExpress) and 20 ng/ml bFGF (#100-18B, PeproTech) on Geltrex coated tissue culture plates. The following day medium was replaced with NMM only. On day 35 neurons of the 3 iPSC lines were pooled at equal ratio and plated on Polyornithine (#P4957, Sigma) and Laminin 521 (#A29249, Thermo Fisher) coated 6 well tissue culture plates for the Mass spectrometry experiments, respectively as single lines and pooled neurons on 96 well imaging plates (#89621, ibidi) for quality control with immunofluorescent staining in NMM supplemented with 10 µM Rho kinase inhibitor. The next day medium was replaced with NMM only. On day 42 and day 50 medium was supplemented with 10 µM DAPT (#MCE-HY-13027, MedChemExpress) to enrich for postmitotic neurons.

#### Immunofluorescent staining of neurons

Neurons cultured in 96 well µ ibidi plates were washed with PBS, fixed with 4 % pFA for 20 minutes followed by pFA quenching with 100 mM glycine.

Samples were blocked with PBS +0.3 % TritonX +10 % normal donkey serum (NDS). Primary antibodies against MAP2 (Millipore, MAB378, 1:300) and TUBB3 (BioLegend, 801202, 1:300) were applied overnight in PBS +0.1 %TritonX +2 % NDS. Neurons were washed 3x with PBS +0.3 % TritonX and secondary antibodies AF 488, Goat anti-mouse IgG1(A21121, 1:500) and AF 546, Goat anti-mouse IgG2a (A21133, 1:500) were applied for 90 minutes in PBS +0.1% Triton X +2 % NDS. After two washes with PBS + 0.3 % TritonX, the neuronswere incubated with PBS+ 0.3 % TritonX +Hoechst for 10 minutes, washed with PBS and mounted with ibidi mounting solution. Images were acquired as z-stacks with a BC43 confocal spinning disc microscope (Andor) using a 60x oil immersion objective.

### Proteomics analysis

#### LiP-MS

The concentrations of cell lysates or brain homogenates were generally similar between replicates, but fine differences were adjusted with LiP buffer. All samples were diluted with LiP buffer to a final protein concentration 2mg/ml and a total volume 50 μl per sample. For interactome experiments, the same freshly prepared SH-SY5Y lysate was split and incubated with 2 μg of αSyn strains or monomer for 15 min at 37°C. For experiments with pure proteins, 2μg of αSyn was used for each replicate. For experiments with brain homogenates or seeded cells, all samples were thawed in parallel and directly exposed to LiP-MS workflow.

LiP-MS experiments were performed as described previously with minor modifications^33,106^. Briefly, Proteinase K (PK) was added with a multichannel pipette at 1:100 enzyme-to-substrate ratio in a total volume of 5 μl. Samples were briefly mixed by gently pipetting up and down five times. Next, the samples were incubated at 37°C for exactly 5 min in a Biometra TRIO thermocycler. Subsequently, PK digestion was stopped by incubating samples at 99°C for 5 min, followed by a 5 min incubation on ice. Next, 55 μl of freshly prepared 10% sodium deoxycholate (DOC) was added.

#### Trypsin / LysC digestion

The samples were reduced with tris(2-carboxyethyl)phosphine hydrochloride (TCEP) to a final concentration of 5 mM at 37 °C for 40 minutes under 800 rpm shaking in thermomixer (Eppendorf). Subsequently, the samples were alkylated by adding iodoacetamide (IAA) at a final concentration of 40 mM and incubating for 30 minutes at room temperature in the dark. To decrease the concentration of DOC to 1%, the samples were diluted using 100 mM ammonium bicarbonate (Ambic). Lysyl endopeptidase LysC (Wako Chemicals) and sequencing-grade trypsin (Promega) were added to the samples at an enzyme-to-substrate ratio of 1:100, and digestion took place in a thermomixer at 37 °C with continuous agitation at 800 rpm overnight. After incubation time passed, the digestion was stopped by adding 100% formic acid (FA) (Carl Roth GmbH) to achieve a final concentration of 3%. As a result, DOC precipitated. Finally, the DOC was eliminated by three cycles of 20-min centrifugation at 21000 rcf, collecting the supernatant each time. The supernatants were desalted on Sep-Pak tC18 (Waters). Samples were eluted with 50% acetonitrile, 0.1% formic acid and dried in a vacuum centrifuge.

#### AP-MS

SH-SY5Y cell pellets were natively lysed in IP buffer (LiP buffer + 1x complete protease inhibitor, 1xPhosSTOP). Lysate was split to the requisite number of aliquots prior to spiking in alpha-synuclein (5 μg) or vehicle control for 1h at RT(end-to-end rotation). VCP was immunoprecipitated from SH-SY5Y lysate with the anti-VCP (10736-1-AP, proteintech) antibody for 2h at room temperature under constant end-to-end rotation using protein A conjugated magnetic beads (Thermo, 88845). As a negative control, an isotype-specific IgG control was used with magnetic protein A beads. Beads were collected on a DynaMag-2 magnetic rack, washed 6x with LiP buffer. Proteins were eluted in 120 μl 8 M urea + 100 mM Ambic for 30 min at 37 °C (1500 rpm, Eppendorf shaker) and snap-frozen. Subsequently, samples were digested with Trypsin and LysC as described previously.

### LC-MS/MS data acquisition

#### Liquid chromatography

LiP-MS and AP-MS samples resuspended in a buffer containing 0.1 % formic acid (FA) were measured on Orbitrap Exploris and Orbitrap Eclipse mass spectrometer (ThermoFisher Scientific), respectively. The mass spectrometer was connected to a Nanoflex electrospray source for nanoelectro spray ionization (nESI). A nano-flow LC system (Easy-nLC 1200, Thermo Fisher Scientific) and self-packed 40 cm x 0.75 mm columns (New Objective) containing 1.9 μm or 3 μm C18 beads (Dr. Maisch Reprosil-Pur 120) were used for peptide separation. Specifically, a linear gradient of lc-buffer A (5 % ACN, 0.1 % FA, Carl Roth GmbH) and lc-buffer B (95 % ACN, 0.1 % FA, Carl Roth GmbH) increasing from 3 % to 30 % lc buffer-B for 120 minutes with a flow rate of 300 nl/minute was used.

#### Data-independent acquisition

Data-independent acquisition (DIA)^107^ scans were performed in 41 variable-width isolation windows. Precursor ions were isolated using a quadrupole. For MS1 survey scans, a mass range of 350-1150 m/z was used and the orbitrap resolution was set to 120’000. A normalized automated gain control (AGC) target of 200 % was applied. High-energy collision induced dissociation (HCD) was employed to fragment precursor ions. DIA-MS/MS spectra were recorded using an orbitrap with a resolution of 30’000 and a mass range of 350-1150 m/z. The maximum injection time was set to 66 ms.

#### Data-dependent acquisition

To generate hybrid spectral libraries, we measured samples in Data-dependent mode (DDA) in parallel with DIA acquisition. Precursor ions were isolated using a quadrupole. For MS1 survey scans, a mass range of 350-1150 m/z was used. HCD was employed to fragment precursor ions. MS/MS spectra were acquired on an orbitrap, at a resolution of 30’000 and AGC target of 200%. The maximum injection time was set to 54 ms.

### Data search, statistical data analysis and visualisation

Hybrid libraries were generated by Pulsar search in Spectronaut, using data from both Data-Independent Acquisition (DIA) and Data-Dependent Acquisition modes. Compared to the default settings, the digestion type was set to ‘semi-specific’. To identify ubiquitinated residues, we additionally searched for GlyGly(K) modification. For analysis, we required that ubiquitinated peptide is detected in >50% of technical replicates.

The peptide precursor abundance comparison was performed using a two-tailed Welch t-test with Benjamini-Hochberg adjustment after median normalisation. Adjusted for multiple comparisons p-values were labelled as q-val. The protein group quantity was taken for protein abundance comparison. Two cut-offs of significance were applied: stringent (FC>2, q-val<0.05) and mild (FC>1.5, q-val<0.05). The statistical data analysis was performed in R, using following R packages: protti^108^, VennDiagram, and limma. For relevant experiments (e.g. when proteins abundance changes might occur), normalisation to tryptic control was performed by dividing peptide intensities on corresponding control protein abundance per peptide, prior to peptide abundance comparison. The protein abundance was identified in the same lysates that were digested only with trypsin, in parallel to LiP-MS workflow. The same sample processing protocol was applied for tryptic control samples, except addition of PK. For amino acid centric analysis^44^, scores were designated to each detected peptide, calculated as a product of the −log_10_(p-value) and |log_2_(FC)|. Subsequently, we calculated the mean value of this score for each amino acid based on overlapping peptides.

Venn Diagrams were plotted with VennDiagram R package. GO enrichments analysis was performed in David^109,110^. GO terms Cellular component and WikiPathways were plotted using R package ggplot2. The set of identified proteins was used as a background for each experiment. To select proteins that belong to Ubiquitin proteasomal system, we manually took protein hits that correspond to Uniprot Key Words Biological process ‘Ubiquitin conjugation’. For chaperones, we have taken hits falling into a Molecular Function ‘Chaperones’. We classified mitochondrial proteins accordingly to biological processes based on Uniprot annotation. Visualisation and mapping LiP-MS hits on protein structures was done in Visual Molecular Dynamics (VMD). Illustrations were performed using Illustrator and Biorender.

### Analysis of αSyn accessibility to PK

To estimate the fraction of αSyn accessible to PK upon cell seeding, we made use of the fully tryptic (FT) peptides that are generated both under LiP conditions (PK digestion in native conditions followed by trypsin digestion in denaturing conditions) and in the tryptic control (TC) samples (only trypsin digestion in denaturing conditions). In the tryptic controls, proteolytic digestion goes to completion and cleavage occurs at defined positions (after K/R residues). Thus, each peptide will be produced at a molarity corresponding to the protein molarity. In contrast, in the LiP samples, any FT peptide that includes a PK cleavage site will show a decrease in intensity corresponding to the extent of cleavage. Comparing the intensities of FT peptides in the two conditions is a measure of the PK accessibility of the corresponding protein region. We calculated the value (TC-LiP)/TC*100% for each fully tryptic peptide, describing its accessibility to PK.

### NMR sample preparation and acquisition

All *in vitro* samples were prepared using phosphate buffer saline (PBS). ^15^N αSyn and UCHL1 were added from stock solutions. To remove oligomeric species, αSyn was filtered with a 100-kDa molecular weight cut off concentrator (Amicon) before being used for NMR sample preparation. The concentrations were determined spectrophotometrically as described previously. Each sample was supplemented with 5% D_2_O and adjusted to a final volume of 100µl in a 3-mm (diameter). The sample contained 15μM ^15^N αSyn and 0, 7.5, 15, or 30μM UCHL1. The samples were measured immediately after preparation. All NMR experiments were performed on a Bruker 600MHz Avance III HD spectrometer with a cryogenic proton-optimized ^1^H(^13^C/^15^N) TCI probe. The pulse sequence employed was a 2D [^15^N,^1^H] SOFAST HMQC NMR experiment. The following parameters were used: 256 x 2048 complex points, 256 scans, 0.5 seconds interscan delay and a spectral width of 11ppm of ^1^H and 30ppm for ^15^N at 10°C. All NMR spectra were processed with TopSpin 4. Visualization and data analysis were carried out in SPARKY.

### Co-sedimentation assay

Purified UCHL1 (BPS Bioscience, 80351) was incubated with each of the three αSyn fibrillar polymorphs for 15 min. The samples were centrifugated at 50’ 000 rpm for 30 min and the pellets were washed twice with PBS.

### Electron microscopy

SH-SY5Y cells or lysates were fixed in fixative solution (2% paraformaldehyde (PFA) and 2.5 % glutaraldehyde (GA)). Fixed cells and lysates were impregnated in 2% low melting point agarose and then transferred on ice for 20 min to solidify agarose. Solid agarose pellet was carefully taken out of the tube and trimmed into small pieces with a razor blade. Cubes containing sample material were then transferred into a small glass vial, washed thoughtfully with 0.1M Cacodylate buffer, post-fixed in 1% buffered osmium tetroxide for 1 h at 4°C, rinsed in distilled water and *en block* stained with aqueous Uranyl Acetate for 1hr at 4°C in the dark. Samples were dehydrated in ascending series of ethanol, and finally, embedded in Epon 812 resin. Ultrathin sections (70 nm) cut by ultramicrotome (Leica EM UC7, Leica, Austria) were contrasted with uranyl acetate and lead citrate and examined under a FEI Tecnai G2 Spirit Transmission Electron Microscope (TEM) operating at 80kV. Images were acquired using anEMSIS Veleta camera (top, side-mounted). The Camera operates using RADIUS software from EMSIS.

### CRISPR Activation Assay

#### Cell Culture for CRISPR

HEK293 QBI cell line stably overexpressing wild-type (WT)αSyn^10^ was cultured in T75 tissue culture flasks (TPP, Trasadingen, Switzerland) in DMEM (#31053-036, ThermoFisher Scientific) + 10% FBS (Hyclone Heat inactivated, #SV30160.03HI, GE Healthcare BioSciences Austria GmbH), 1% GlutaMax (Gibco), and 1% Penicillin/Streptomycin (Gibco) supplemented with 0.4 mg/ml geneticin (#10131035, ThermoFisher Scientific). Cells were transfected with the dCas9VPR plasmid pXPR_120 (#96917, Plasmid Addgene) using Lipofectamine 2000 (Thermo Fisher Scientific) according to the manufacturer’s instructions and selected with blasticidin at a concentration of 10 μg/mL to obtain stable transfected cells and the resulting cells are denoted as dCas9-aSyn-HEK Cells and were continuously maintained under antibiotic selection (geneticin and blasticidin).

#### CRISPR Activation and aSyn Phosphorylation Assay

All the CRISPR activation gRNA plasmids with four non-overlapping guide RNAs (gRNAs) per tested gene candidate were obtained from an in-house genome-wide CRISPR activation library (T. gonfio library)^111^. The gRNA plasmids were miniprepped using the QIAprep Spin Miniprep Kit (Qiagen). 3000 dCas9-aSyn-HEK cells were seeded in a PDL coated 384-well plate (#6055500, CellCarrier Ultra microplates). For the delivery of the gRNA plasmids, transfection was performed using the ViaFect™ Transfection Reagent, (#E4981, Promega). Following transfection, the cells were selected with Puromycin. 48hrs post-selection patient brain-amplified fibrils (PD5405, DLB163 and MSA043) at a concentration of 7.5μg/ml^53^ were complexed with the TransIT^®^-LT1 Transfection Reagent (#MIR 2306) and treated for 72 hours. The experiment was conducted with 4 technical replicates (n=4).

#### Immunostaining, Imaging and data analysis

Cells were washed with Tris-Buffered Saline (TBS) and fixed with 4% PFA (Thermo Fisher Scientific), permeabilized with 0.1% Triton X-100 in TBS, and blocked with 4% BSA in TBS for 1 hour followed by the incubation with primary antibody, P-syn/81A Antibody (#825701, BioLegend) diluted in 0.5% BSA in TBS at a dilution of 1:2500. The secondary antibody used was Goat anti-Mouse IgG (H+L) Alexa Fluor™ 488 (#A-11001, Thermo Fisher Scientific), diluted in 0.5% BSA in TBS at a dilution of 1:400. For labeling the entire cell, HCS Cell Mask Stain at a dilution of 1:5000 in PBS (#H32721, Thermo Fisher Scientific) was used, and the nucleus was stained with DAPI at a dilution of 1:10000 (#D9542, Sigma). Imaging was performed using the Widefield - GE IN Cell Analyzer 2500HS with a 10X or 20X objective. The total number of cells and the percentage of cells containing pSyn inclusions were counted using ilastik 1.4.0 and CellProfiler 4.2.1. We used unpaired two-tailed Student’s t-test to identify genes that significantly modulate P-αSyn positive inclusion in comparison to non-targeting controls.

#### qPCR analysis

Cells were washed twice with PBS and lysed using RLT lysis buffer (Qiagen). RNA was extracted using the RNeasy Mini Kit (Qiagen) following the manufacturer’s instructions and eluted in ddH2O. Quality assessment and concentration measurement of RNA was done using a NanoDrop spectrophotometer (Thermo Fisher Scientific). For each sample, 1000ng of total RNA was reverse transcribed to cDNA using the Quantitect Reverse Transcription Kit (Qiagen) according to the manufacturer’s guidelines. The cDNAs corresponding to the genes of interest were individually quantified using qPCR analyses based on the SYBR Green Mastermix (Roche). The ViiA 7 Real-Time PCR System (Thermo Fisher Scientific) was employed for data acquisition. Actin-beta was used as the housekeeping gene for normalization across samples. Primer pairs for the genes of interest are listed in Table 2.

**Table 2.**
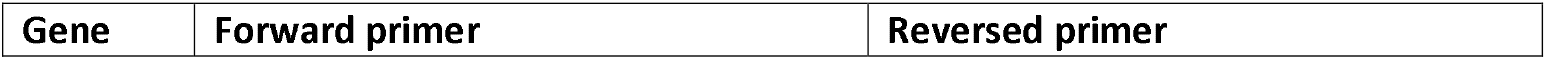
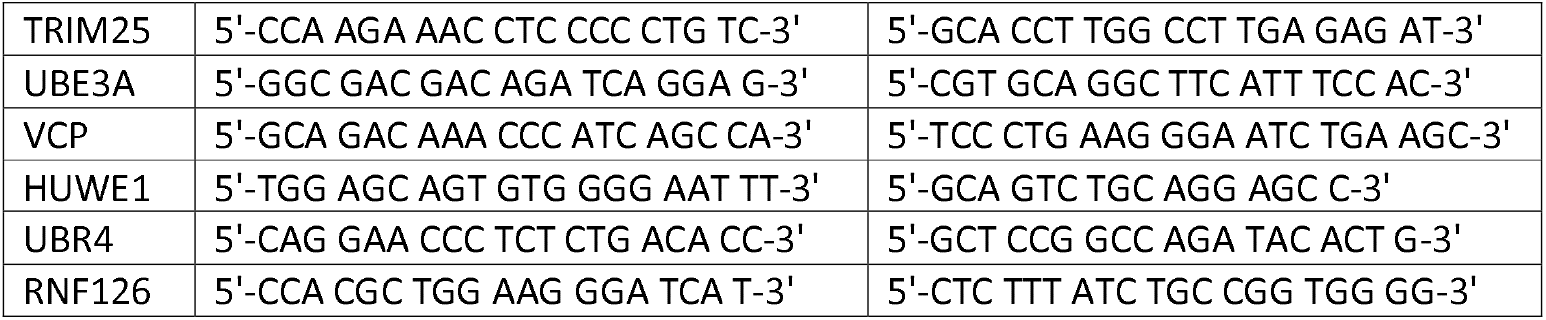
Primer pairs for the genes of interest used for qPCR.

## Ethics statement

The lines SFC840-03-03, SFC854-03-02 and SFC856-03-04 were derived from dermal fibroblasts from healthy donors trough the Oxford Parkinson’s Disease Centre (SF840, SF856, SF854): participants were recruited to this study having given signed informed consent, which included derivation of hiPSC lines from skin biopsies (Ethics Committee that specifically approved this part of the study: for control donors, National Health Service, Health Research Authority, NRES Committee South Central, Berkshire, UK, REC 10/H0505/71, all experiments were performed in accordance with UK guidelines and regulations and as set out in the REC.

## Acknowledgements

We thank Alexis Fenyi for his involvement in setting up the *ex vivo* amplification reactions of pathogenic αSyn. We thank Markus Britschgi (Roche), Norbert Volkmar (DISCO Pharmaceuticals), Ludovic Gillet (IMSB ETHZ), Viviane Reber (IMSB ETHZ), and Christian Dörig (Roche) for fruitful discussions. We thank Sally Cowley (Oxford Parkinson’s Disease Center) for providing the iPSC lines used in this study. We thank BioEM lab of University of Basel for performing EM experiments. We thank the Parkinson’s UK Tissue Bank at Imperial College London for their assistance, and we express our deepest appreciation to the brain tissue donors and their families.

RM was supported by France Parkinson, the Fondation pour la Recherche Médicale (ALZ201912009776) and the ERA PerMed Era-Net Cofund (project DEEPEN-iRBD, ANR-22-PERM-0006). VR was supported by INSERM. P.P. was funded by a Personalized Health and Related Technologies (PHRT) grant (PHRT-506), a Sinergia grant from the Swiss National Science Foundation (SNSF grant CRSII5_177195), the Peter Bockhoff Stiftung and the ETH Zurich foundation, Parkinson Schweiz, the EMPIRIS Foundation grant (2022-FS-353), the Synapsis Foundation - Alzheimer Research Switzerland ARS, The European Research Council (866004), TS was funded by a Gelu foundation grant and the Synapsis Foundation-Alzheimer Research Switzerland ARS grant (No. 2023-CDA02). S.K. and M.G. have received support from the EU/EFPIA/OICR/McGill/KTH/Diamond Innovative Medicines Initiative 2 Joint Undertaking (EUbOPEN grant no. 875510). WH and RBG are supported by the URPP AdaBD (Adaptive Brain Circuits in Development and Learning) of the University of Zurich.

MS analysis of covalent painting has benefited from access to the facilities of the I2BC proteomics platform (Proteomique-Gif, SICaPS, supported by IBiSA, Ile de France Region, Plan Cancer, CNRS and Paris-Saclay University).

## Code availability statement

The data analysis was performed using R packages and other software described in the Methods section, which were reported previously. Further code for plots and other analyses is available upon request.

## Author contributions

T.S. conceived the project, designed and performed the experiments, and analyzed the data. V.R. and R.M. conceived and designed experiments related to molecular painting-based structural characterization of the strains. V.R. optimized and supervised molecular painting experiments and analyzed the data. J.S. amplified the fibrils and contributed to molecular painting experiments. R.M. designed and performed experiments on strain characterization, brain homogenate preparation, and PTMs detection in human brain homogenates. S.N. performed CRISPR activation experiments. W.J.H. differentiated iPSCs into neurons and performed IF staining. Y.F. performed NMR experiments. C.T. performed EM experiments on cells. T.B. contributed to the data analysis. S.K. and M.G. contributed to the Western blot and AP-MS experimental set up. S.G. provided postmortem brain tissue samples. R. B.-G. supervised iPSCs differentiation into neurons. A.A. supervised CRISPR activation experiments. R.R. supervised NMR experiments. N.d.S. and T.B. contributed to experimental set up. T.S., V.R., N.d.S, P.P., and R.M. wrote and revised the manuscript with contributions from all authors. P.P. and R.M. supervised the project.

