## Supplementary Information for "Structure-function relationship of alpha-synuclein fibrillar polymorphs derived from distinct synucleinopathies"

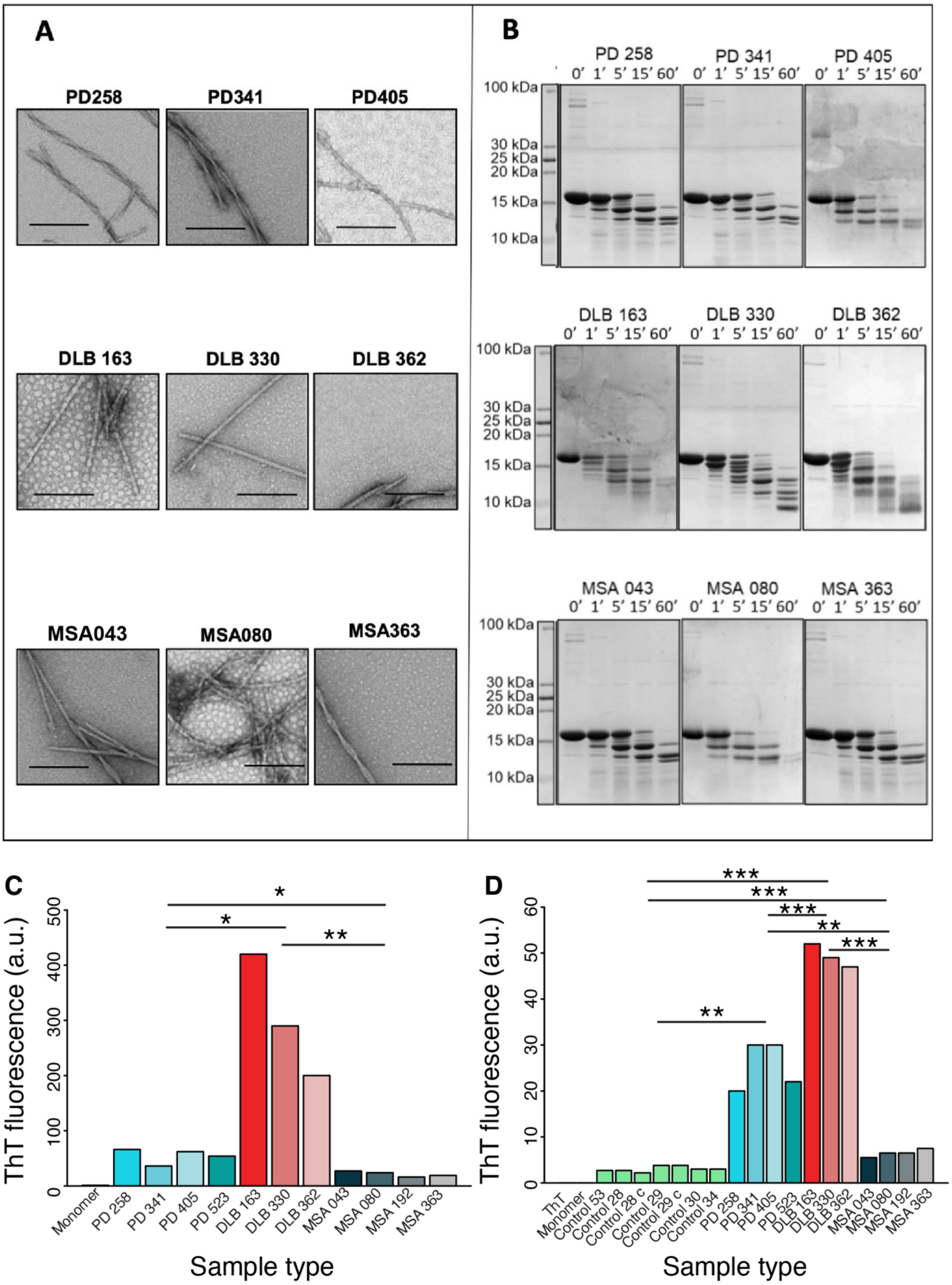


**Supplementary Figure 1. Characterization of αSyn fibrils PMCA-amplified from PD, MSA and DLB patient brains.** Transmission electron microscopy of patient derived αSyn fibrils (**A**). Scale bar = 200 nm. Limited proteolytic profiles of patient derived αSyn fibrils (**B**). Digestion of αSyn samples (100 µM monomeric concentration) in presence of Proteinase K (3.8 µg/ml) was monitored over time at 37°C and stopped after 0, 1, 5, 15, 60 min digestion by addition of 100 µM PMSF. Limited proteolysis products were dissolved by Hexafluoroisopropanol before denaturation, SDS-PAGE separation and Coomassie staining. Time (min), molecular weight markers (MW, kDa) are shown on the left side and the top of the gels. (**C, D**) Thioflavin T fluorescence of amplified strains (**C**) and of brain homogenates used as starting material for amplification. (**D**). Stars indicate statistical significance (*p-val<0.05, **p-val<0.01, ***p-val<0.001).

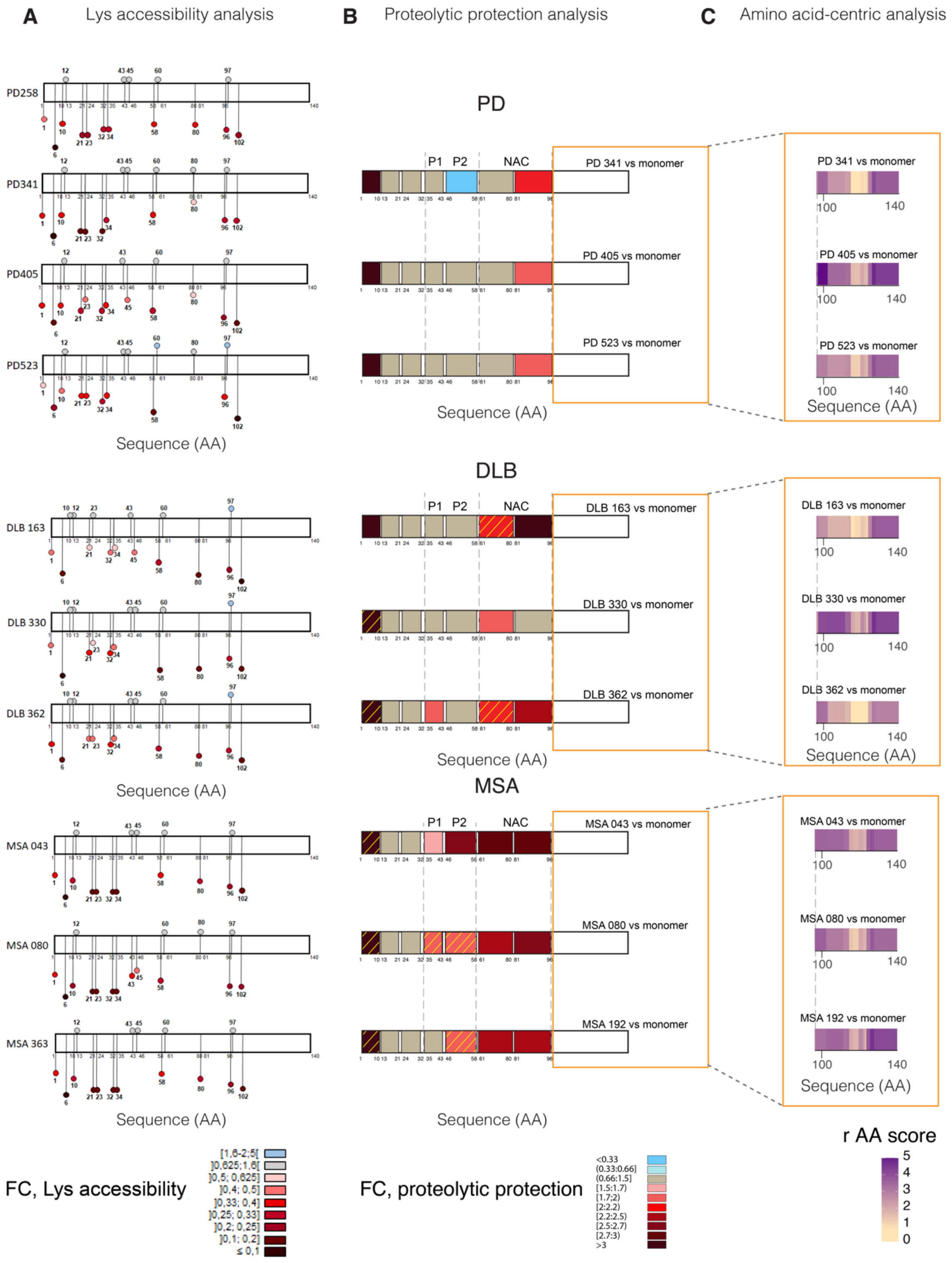


**Supplementary Figure 2. Disease-specific structural features of PMCA-amplified fibrils from individual patients.** Lys accessibility (**A**), proteolytic protection (**B**) and amino acid-centric analysis of C-terminus proteolysis (**C**) are shown. (**A**) Covalent painting-based Lys accessibility analysis of PD, DLB and MSA derived αSyn fibrils for individual patients vs αSyn monomer. The color scale shows fold change of lysine accessibility vs monomer at all lysines within αSyn primary structure; darker hues indicate decreased accessibility (n=4 patients for PD, and n=3 patients each for DLB and MSA). (**B**) LiP-MS-based proteolytic protection analysis for PD, DLB, and MSA fibrils vs αSyn monomer for individual patients. The color scale shows fold change of proteolytic protection vs monomer along αSyn primary structure; darker hues show increased protection (n=3 patients per disease, n=4 technical replicates per sample). The hatched area indicates the regions exhibiting FC>1.5 that did not pass the significance cut-off (p-val<0.05), and corresponding statistically significant changes were recorded for the same region for at least one other patient. (**C**) LiP-MS-based amino acid-centric analysis of proteolytic patterns in αSyn C-terminal moiety of PD, DLB and MSA fibrils for individual patients vs αSyn monomer. The color scale shows the r score, a measure of the change in protease accessibility per amino acid, plotted along the αSyn C terminal primary structure. Non-significantly different regions are colored in beige. Significantly changing regions are colored in violet. Darker violet color indicates stronger difference. N=3 patients per disease, n=4 technical replicates per sample.

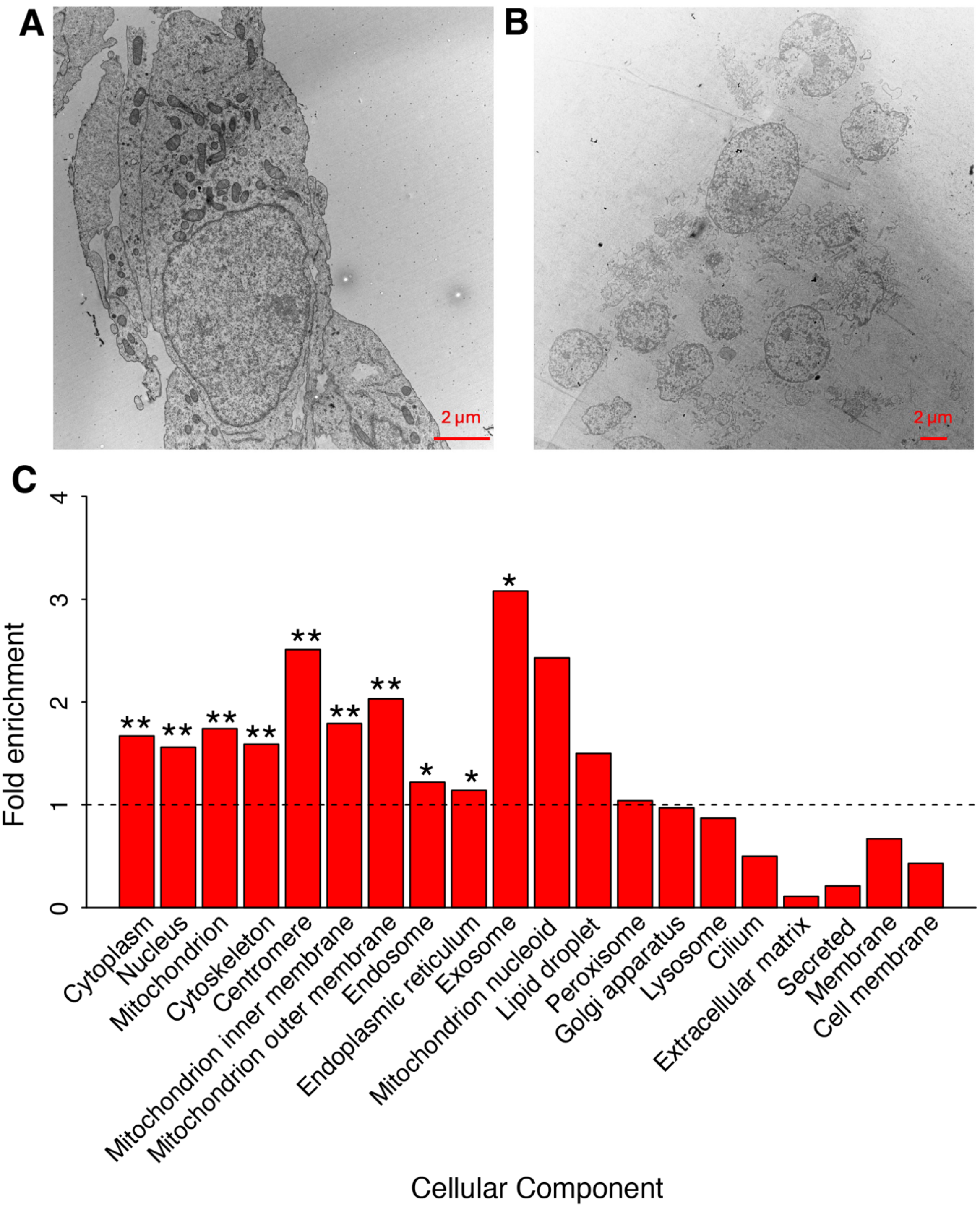


**Supplementary Figure 3. Evaluation of cell lysis efficiency.** (**A**, **B**) Electron microscopy of intact (A) and lysed (B) SH-SY5Y cells. (**C**) Enrichment analysis (Uniport Keywords cellular component) of proteins represented by semi-tryptic peptides (i.e., those produced by PK cleavage and detected in SH-SY5Y lysate). Stars indicate statistical significance (*q-val <0.05, **q-val<0.01)

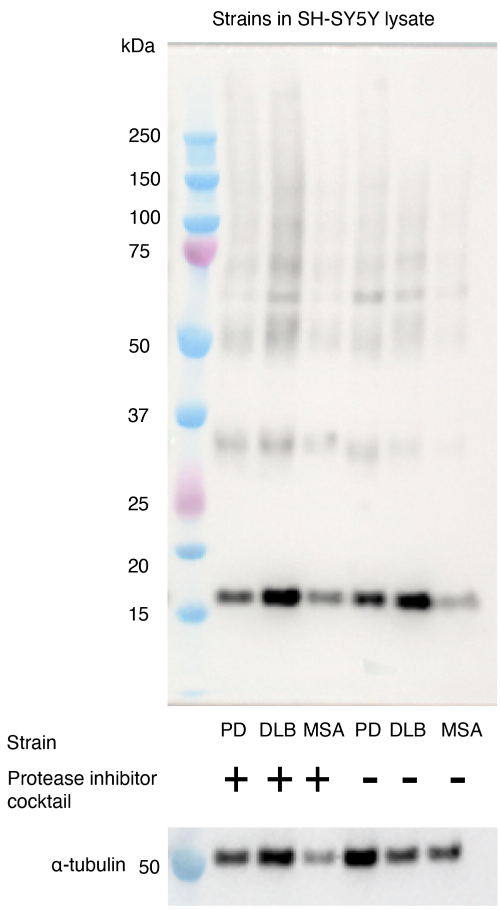


**Supplementary Figure 4. Effect of protease inhibitors on αSyn cleavage in SH-SY5Y lysate**. Western blot against αSyn (stained with the 5G4 anti-αSyn antibody, epitope aa 46-53) after incubation of the same amount of αSyn fibrillar strains (1.5 µg) in the presence or absence of a cOmplete protease inhibitors cocktail (Roche) for 15 min in cell lysate. α-tubulin served as a loading control.

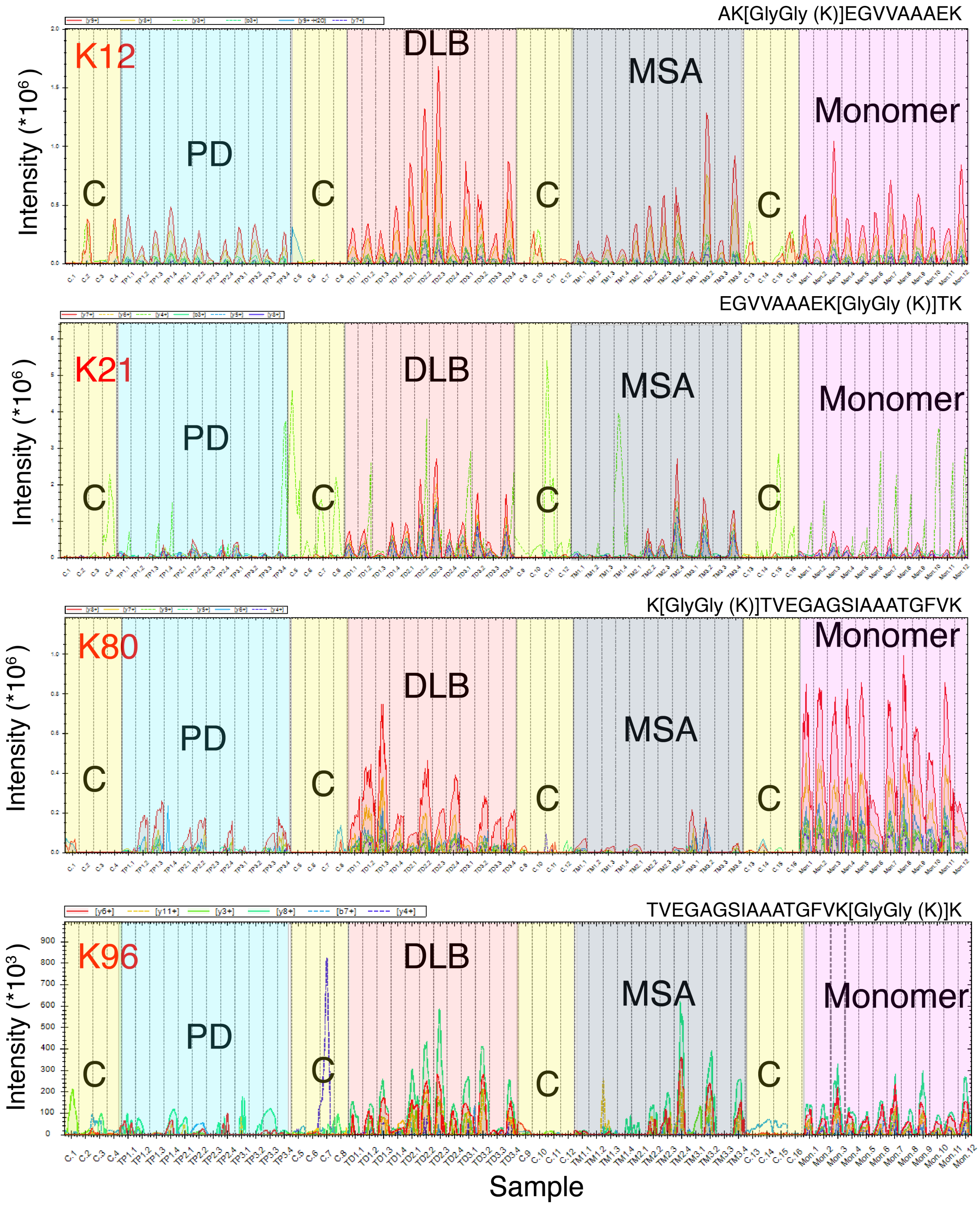


**Supplementary Figure 5. Quality control of the identification of ubiquitinated peptides.** Extracted ion chromatograms of fragment ions corresponding to identified ubiquitinated peptides of αSyn upon spiking into SH-SY5Y lysate for 15 min. Different colors represent different fragments. Dashed peaks correspond to contaminants. Conditions are labeled as follows: PD, DLB, MSA, Monomer for spiked αSyn, C- control SH-SY5Y lysate. Normalization on protein abundance was performed at the later data analysis stage.

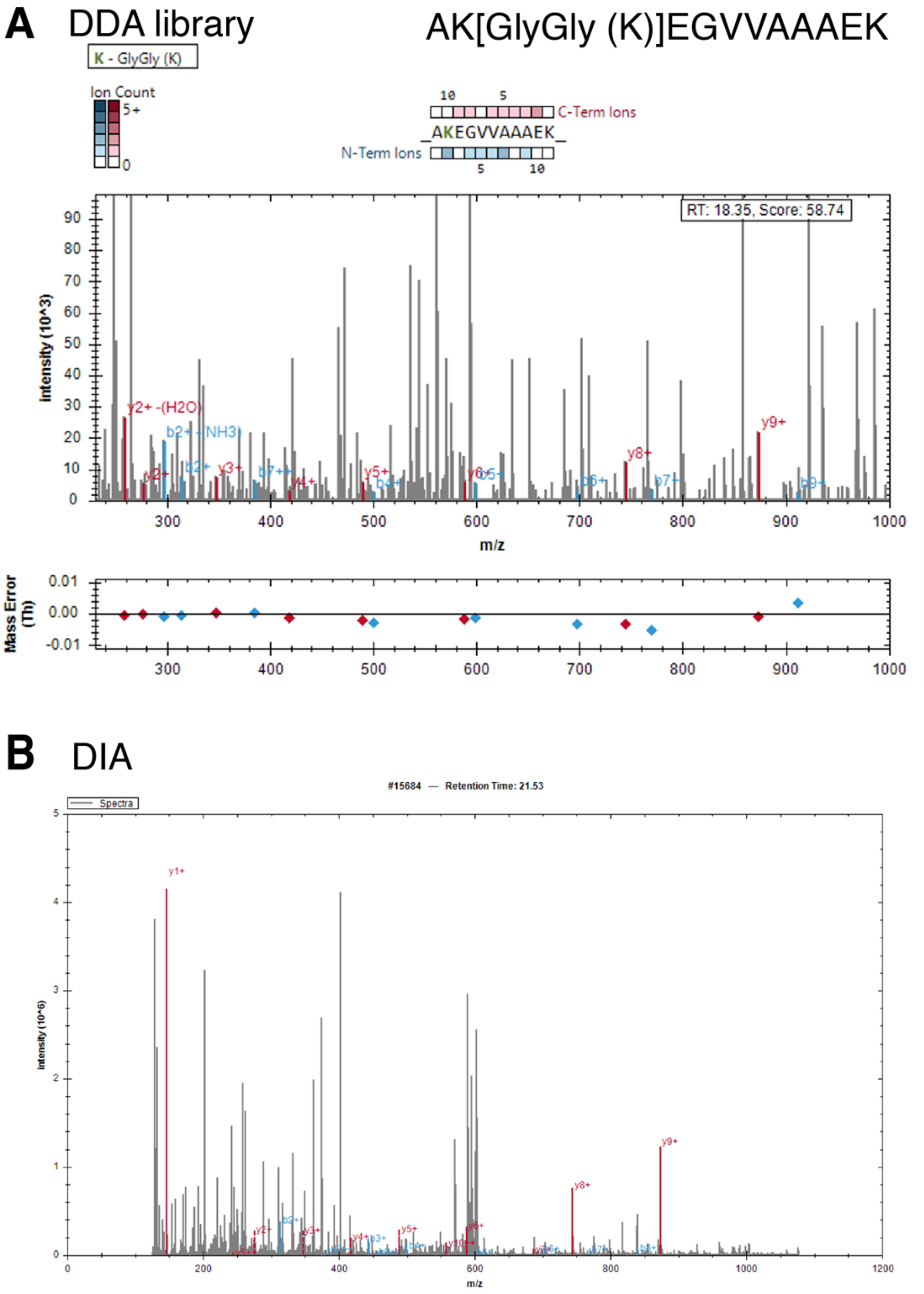


**Supplementary Figure 6. Quality control of the identification of the peptide AK[GlyGly(K)]EGVVAAAEK.** MS2 spectra corresponding to identified ubiquitinated peptides of αSyn upon spiking into SH-SY5Y lysate acquired in (**A**) DDA (for library) and (**B**) DIA mode.

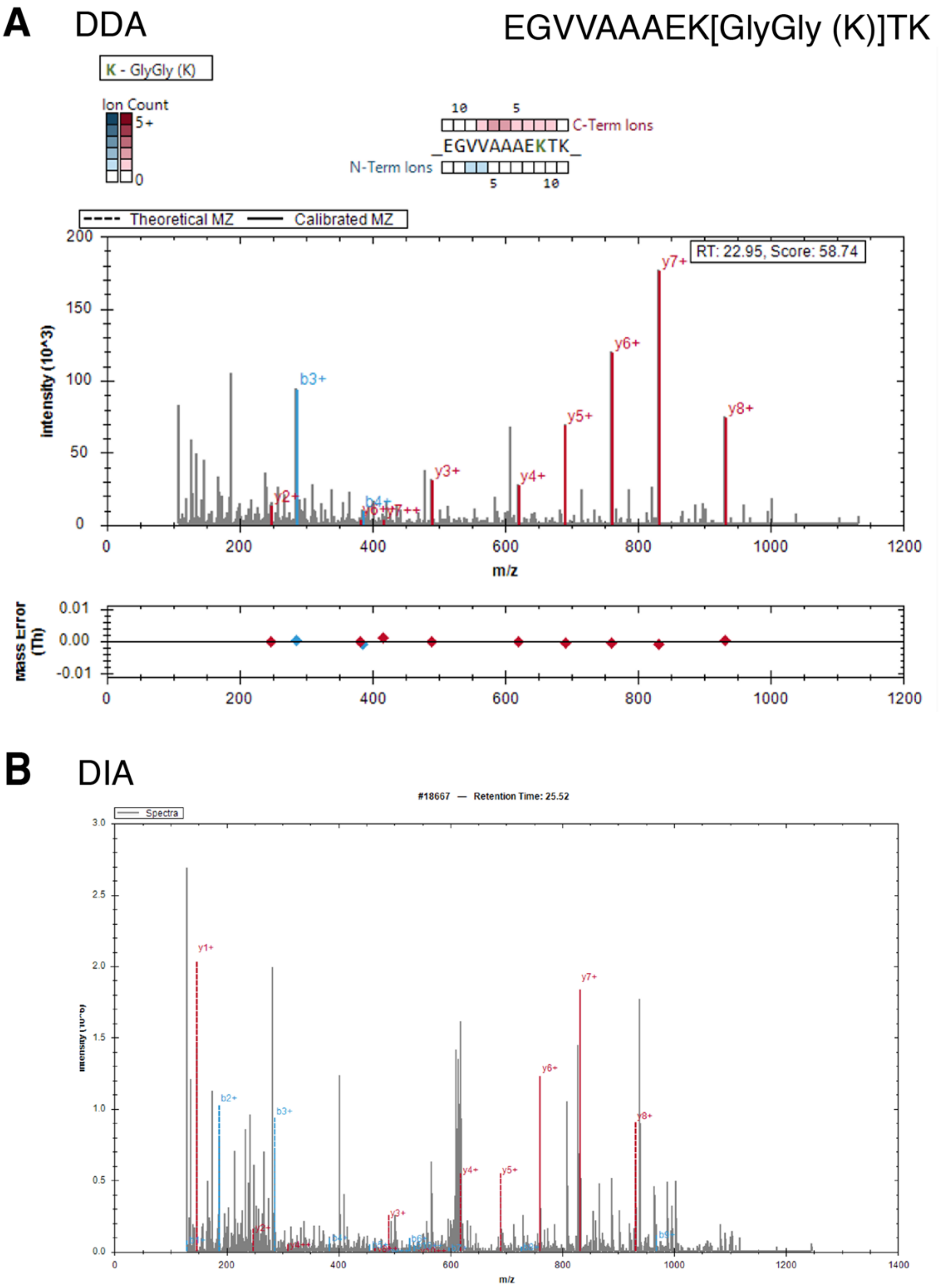


**Supplementary Figure 7. Quality control of the identification of the peptide EGVVAAAEK[GlyGly(K)]TK.** MS2 spectra corresponding to identified ubiquitinated peptides of αSyn upon spiking into SH-SY5Y lysate acquired in (**A**) DDA (for library) and (**B**) DIA mode.

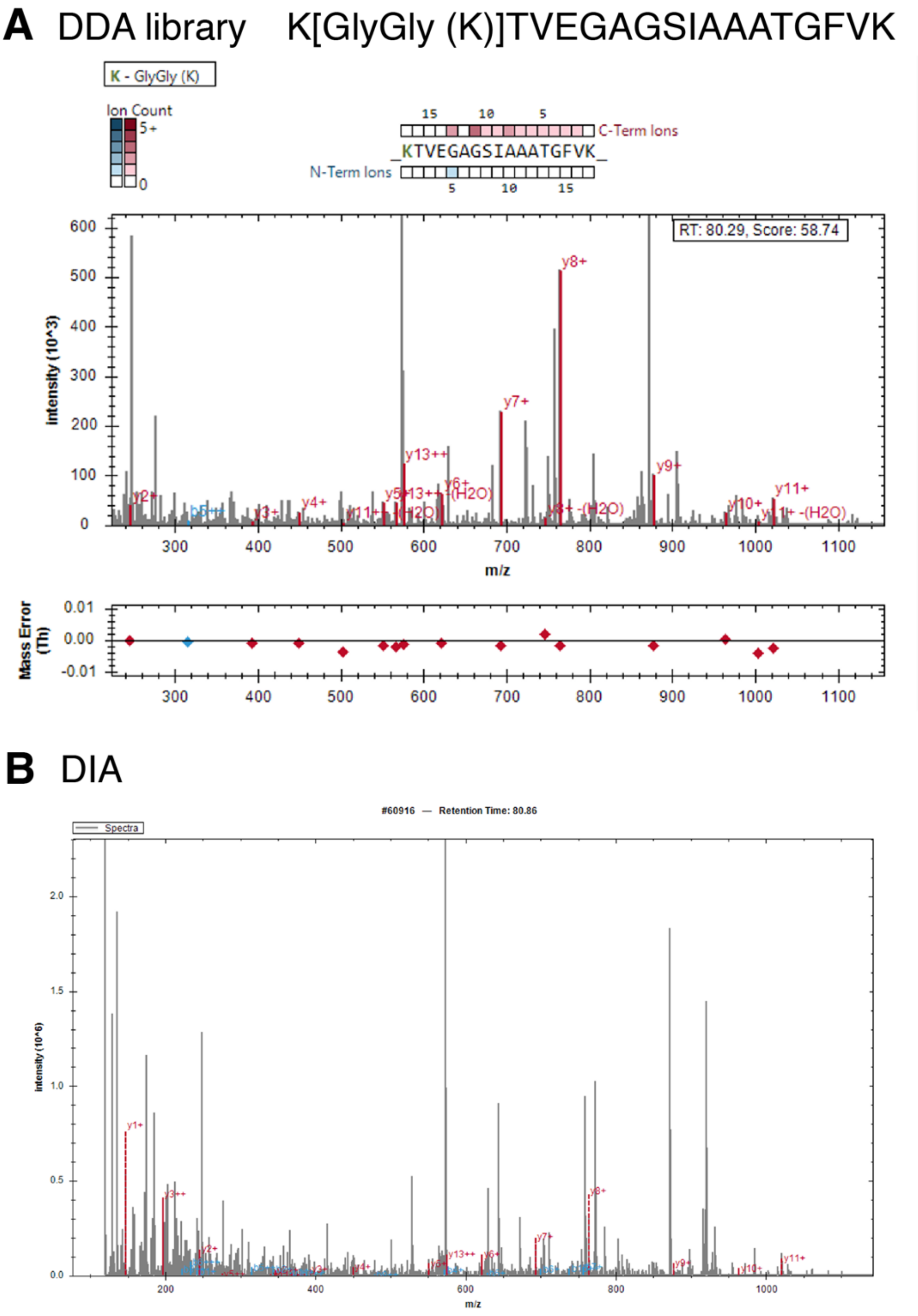


**Supplementary Figure 8. Quality control of the identification of the peptide K[GlyGly(K)]TVEGAGSIAAATGFVK.** MS2 spectra corresponding to identified ubiquitinated peptides of αSyn upon spiking into SH-SY5Y lysate acquired in (**A**) DDA (for library) and (**B**) DIA mode.

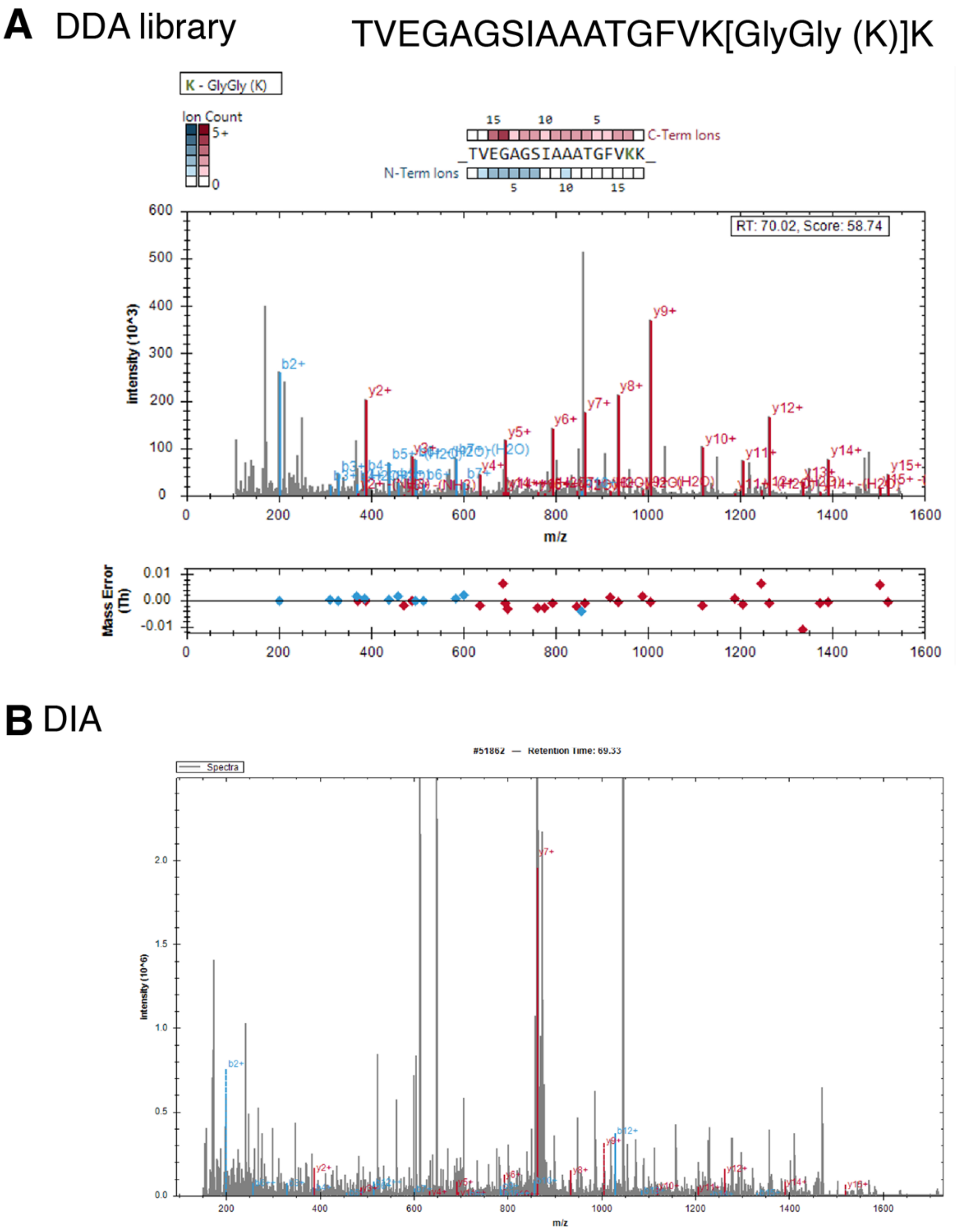


**Supplementary Figure 9. Quality control of the identification of the peptide TVEGAGSIAAATGFVK[GlyGly(K)]K.** MS2 spectra corresponding to identified ubiquitinated peptides of αSyn upon spiking into SH-SY5Y lysate acquired in (**A**) DDA (for library) and (**B**) DIA mode.

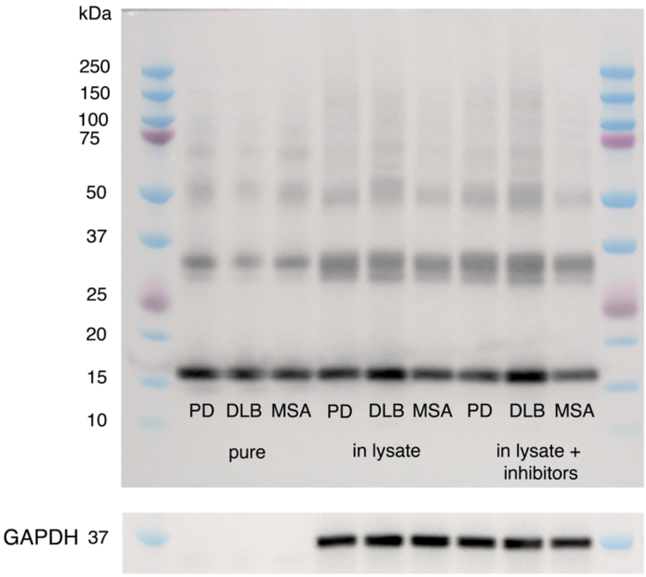


**Supplementary Figure 10. Western blot analysis of αSyn fibrillar polymorphs in pure form and after incubation in cell lysate in the absence or presence of proteases inhibitors**. The Western blot against αSyn (stained with 5G4 anti-αSyn antibodies) for amplified pure PD, DLB and MSA strains (first three lanes). Western blot against αSyn after incubation of the same amount of αSyn (1.5 μg) for 15 min in SH-SY5Y cell lysate in the absence or the presence of a cOmplete protease inhibitors cocktail (Roche). GAPDH was used as a loading control (anti-GAPDH antibody GA1R). The samples were incubated for 60h in 8M urea to disassemble the fibrils.

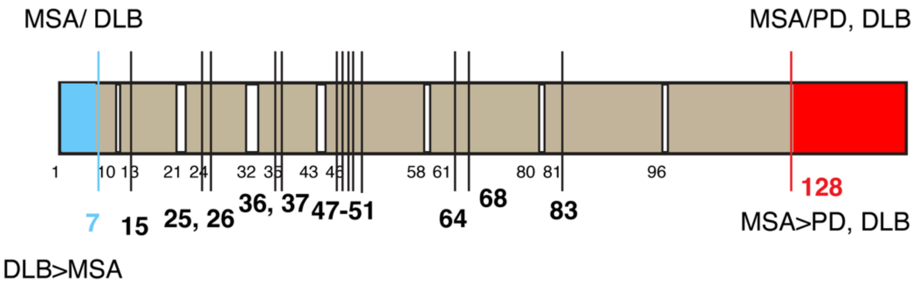


**Supplementary Figure 11. Mapping cell lysate endogenous protease cleavages on the sequence of αSyn**. Vertical lines show endogenous cleavage sites, detected based on semi-tryptic peptides found in tryptic control data (i.e., where no PK is added). Cleavage products that are significantly different between strains are labeled in blue and red.

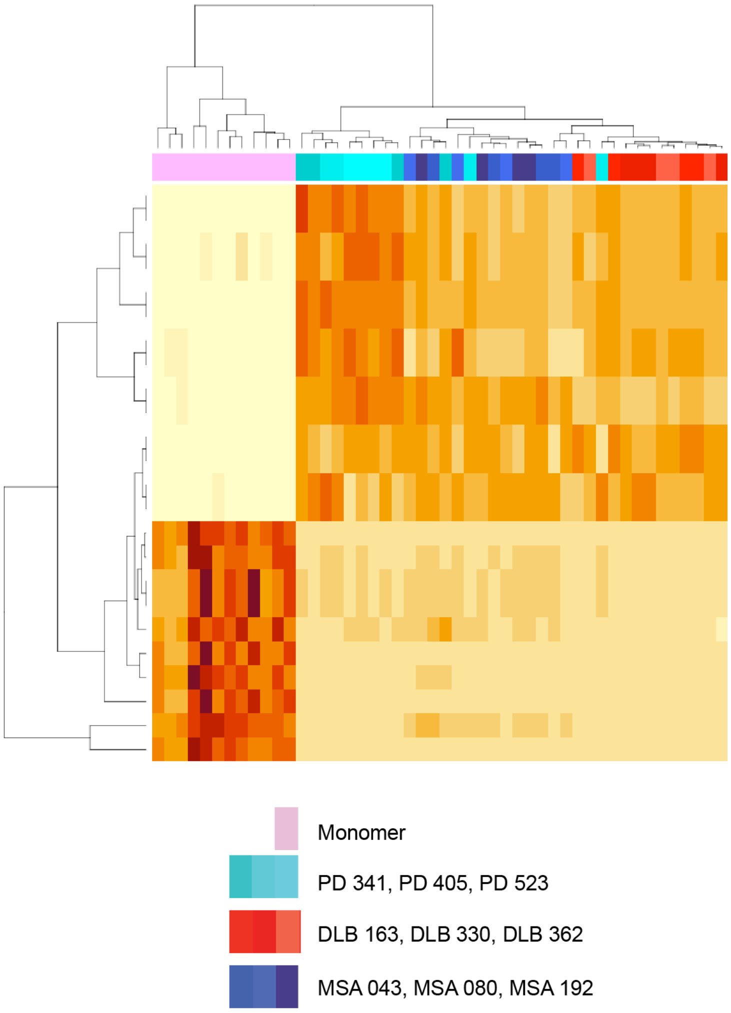


**Supplementary Figure 12**. **Conformation-specific peptide patterns of αSyn species.** Clustering analysis of all peptides for monomeric αSyn (pink) and PD (cyan), DLB (red) and MSA (dark blue) fibrils. Individual patients are shown with different shades of each colour. N=3 patients per disease, n=4 technical replicates per sample.

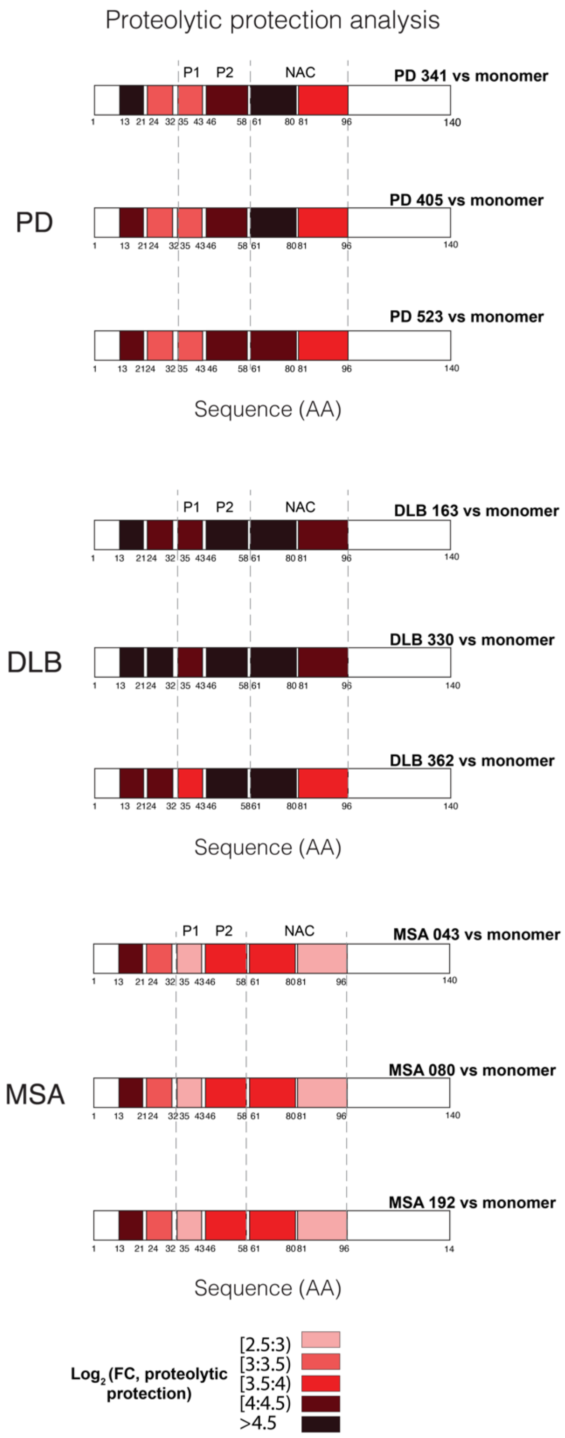


**Supplementary Figure 13. Disease-specific structural features of PMCA-amplified fibrils from individual patients *in situ* (lysate).** LiP-MS-based proteolytic protection analysis for PD, DLB, and MSA fibrils vs αSyn monomer for individual patients. The colour scale shows fold change of proteolytic protection vs monomer along the αSyn primary structure; darker hues show increased protection (n=3 patients per disease, n=4 technical replicates per sample).

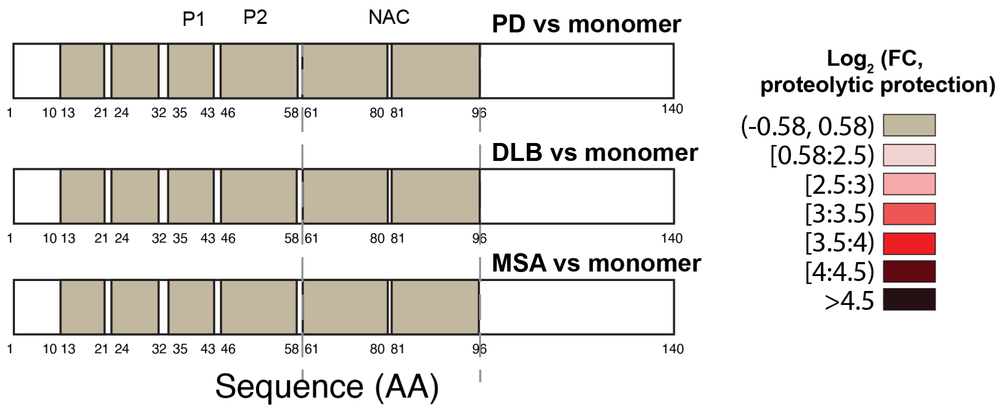


**Supplementary Figure 14. Tryptic control data shows no protection in fully tryptic peptides of αSyn**. Grey bars correspond to non-significantly different peptides between the fibrillar polymorphs and the monomer in trypsin only digestion condition. This data supports that additional protection of fully peptides in LiP condition is due to difference in structure.

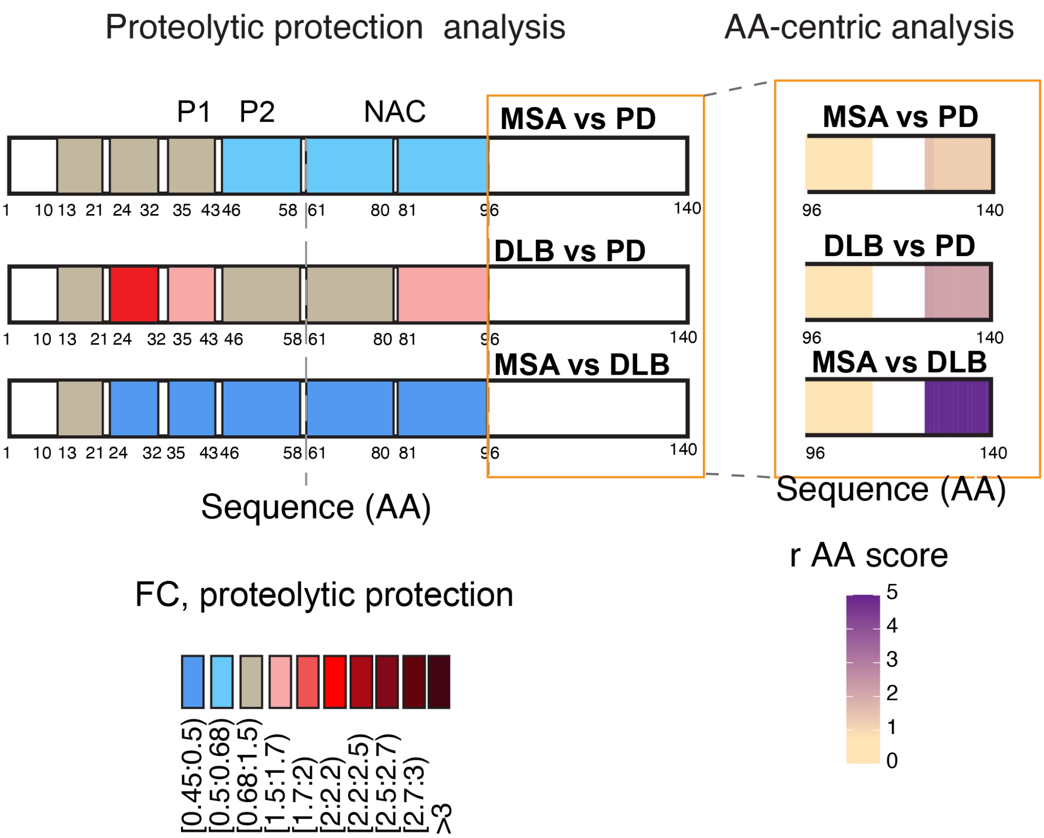


**Supplementary Figure 15. Direct structural comparison of PD, DLB and MSA strains in SH-SY5Y cellular lysate.** LiP-MS-based proteolytic protection analysis for PD, DLB, and MSA derived αSyn fibrils vs each other in cellular lysate (left). The color scale shows the fold change of proteolytic protection vs each other along the αSyn sequence; darker red hues show increased protection, and darker blue hues indicate decreased protection (n=3 patients per disease, n=4 technical replicates per sample). LiP-MS-based amino acid-centric analysis of proteolytic patterns in αSyn C-terminal moiety of PD, DLB, and MSA derived αSyn fibrils vs each other (right). The color scale shows the r score, a measure of the change in protease susceptibility per amino acid, plotted along the αSyn C-terminal primary structure (n=3 patients per disease, n=4 technical replicates per sample).

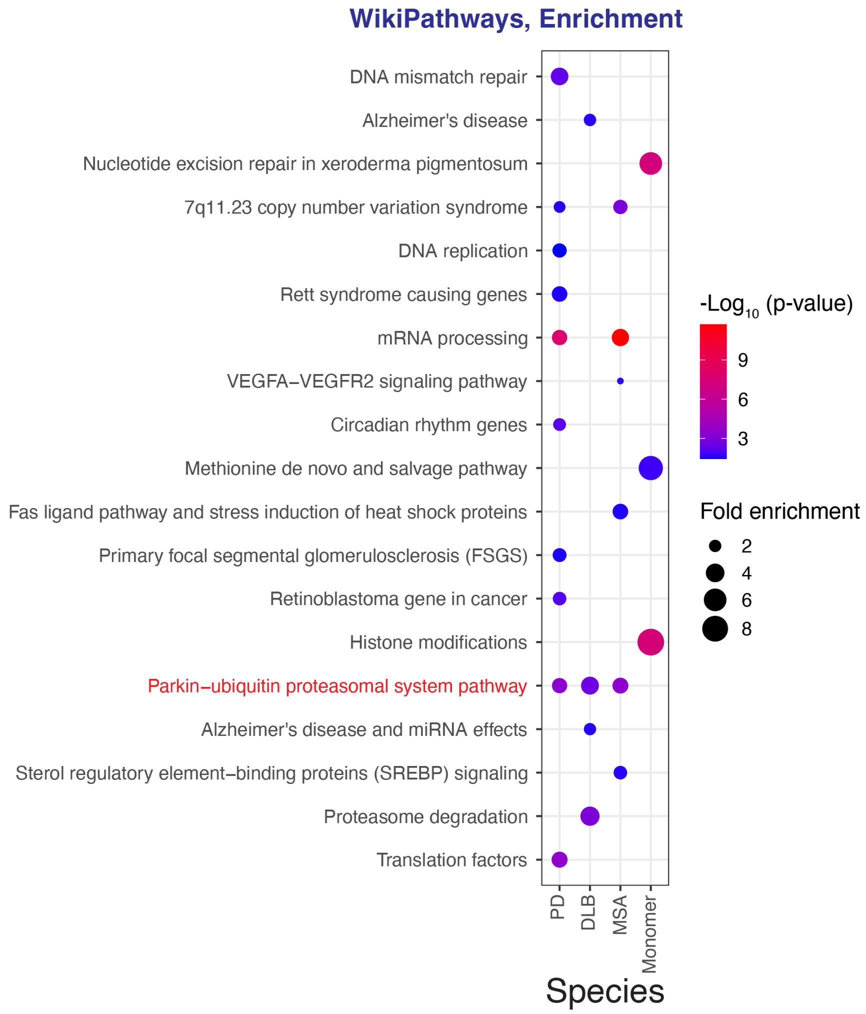


**Supplementary Figure 16. Pathways enrichment analysis for interactors of the different αSyn species.** Functional enrichment analysis (WikiPathways) for the set of proteins that show structural changes upon spiking of the different αSyn species into SH-SY5Y lysate. All significant enrichments are shown (p-val<0.05).

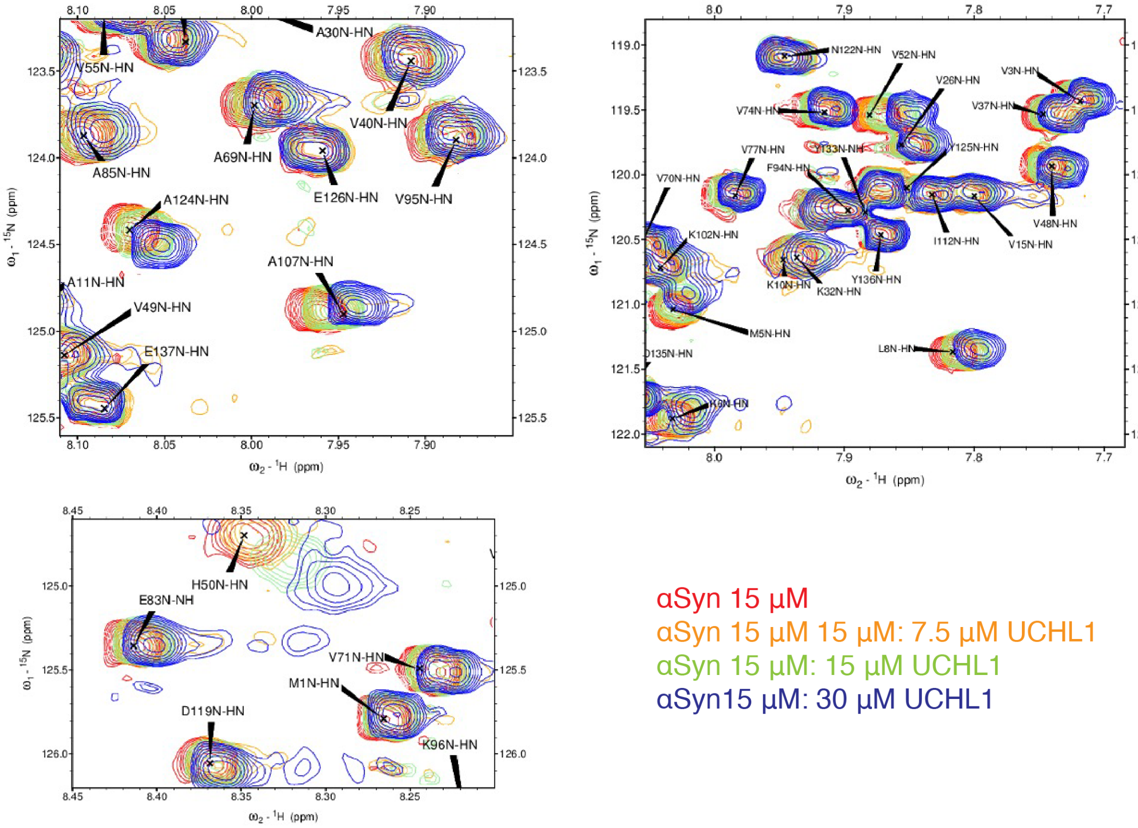


**Supplementary Figure 17. NMR validation of interaction between UCHL1 and αSyn monomer.** 2D [$^{15}$N,$^{1}$H] HMQC NMR spectra of 15 μM $^{15}$N labelled αSyn in the absence (red) or presence of increasing quantities of purified UCHL1 (yellow, green, and magenta) in PBS. Black arrowheads are used to label the peaks.

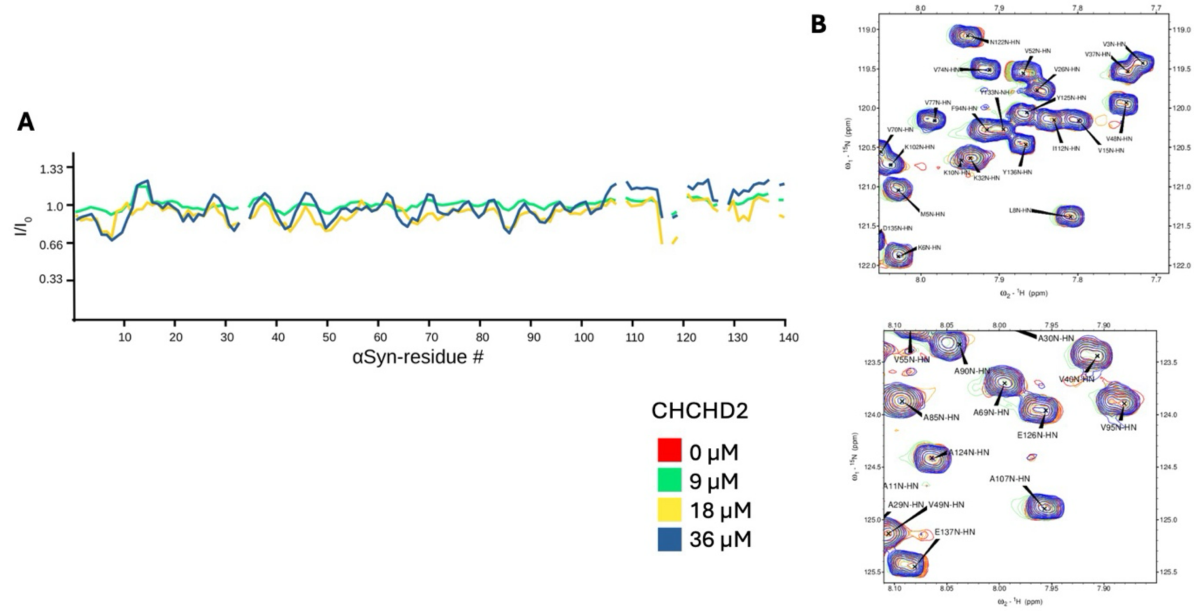


**Supplementary Figure 18. Control of αSyn interaction specificity measured by NMR.** No direct interaction between αSyn with CHCHD2 measured on $^{15}$N-labeled αSyn. 2D [$^{15}$N,$^{1}$H] HMQC NMR of 18μM wild-type αSyn in PBS at pH 7.4 with varying concentrations of human CHCHD2. (**A**) Peak intensity ratios of wild-type αSyn. (**B**) Chemical shift perturbations (CSPs) of wild-type αSyn with subsections of the HMQC spectrum with peak labels (lower panel).

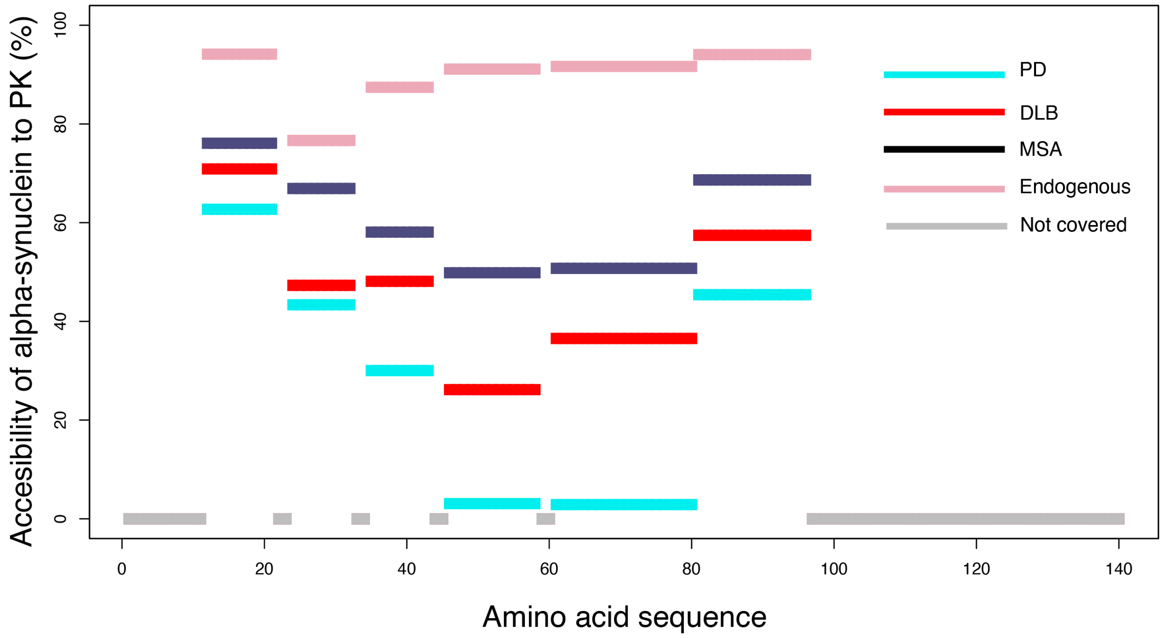


**Supplementary Figure 19. Accessibility of αSyn to PK in SH-SY5Y cell seeding model.** Percentage of the loss of intensity of fully tryptic peptides under PK digestion of αSyn in the cell seeding model (accessibility to PK). The accessibility of endogenous αSyn (without seeding) in SH-SY5Y cells is given in pink. Peptides are mapped along αSyn sequence. The αSyn sequence regions that are not covered by fully tryptic peptides are shown in grey.

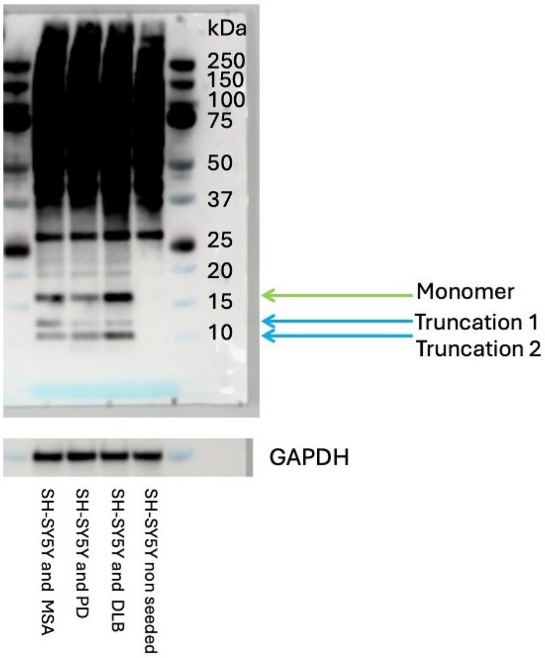


**Supplementary Figure 20. Two truncated species of αSyn revealed after strains uptake by live SH-SY5Y cells.** Western blot analysis of αSyn strains using a mixture of two antibodies (5G4 (aa 47-52) and 42/α-Synuclein (aa 91-99)) in lysates of living SH-SY5Y cells that were incubated with 250 nM of each αSyn strain for 24h. Green arrow indicates the monomer. Cyan arrows indicate truncated forms. SDS-PAGE was done using MES buffer. The higher molecular weight signal is primarily due to antibody nonspecific binding in the SH-SY5Y lysate. GAPDH served as a loading control using GA1R anti-GAPDH antibody on the same blot.

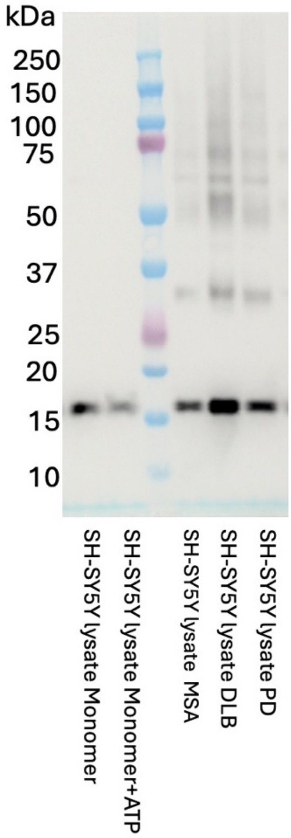


**Supplementary Figure 21.** Western blot analysis of αSyn monomer and depolymerized αSyn fibril strains spiked into SH-SY5Y lysate using a mixture of two antibodies 5G4 (epitope aa 47-52) and 42/α-Synuclein (aa 91-99) antibody.

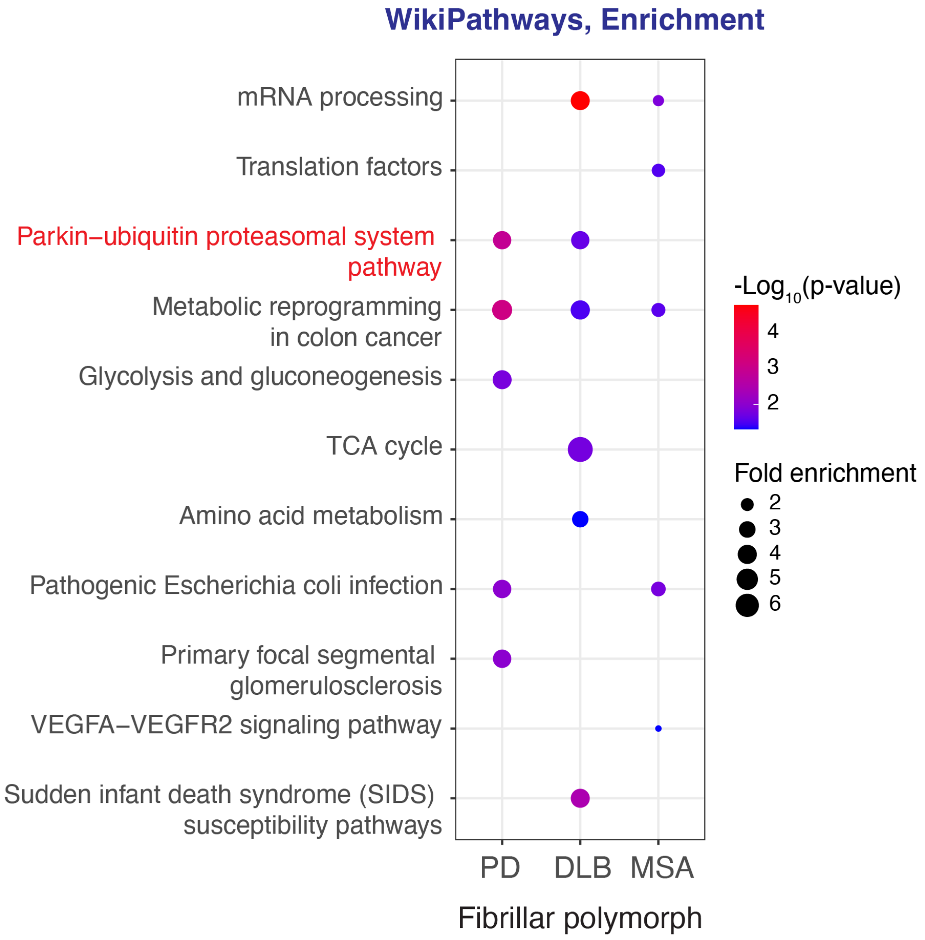


**Supplementary Figure 22. Pathway enrichment analysis for the set of proteins that structurally respond to the different αSyn fibrillar polymorphs in living SH-SY5Y cells.** Functional enrichment analysis (WikiPathways) for the set of proteins that show structural changes upon cell infection with the different αSyn fibrillar polymorphs. All significant enrichments are shown (p-val<0.05).

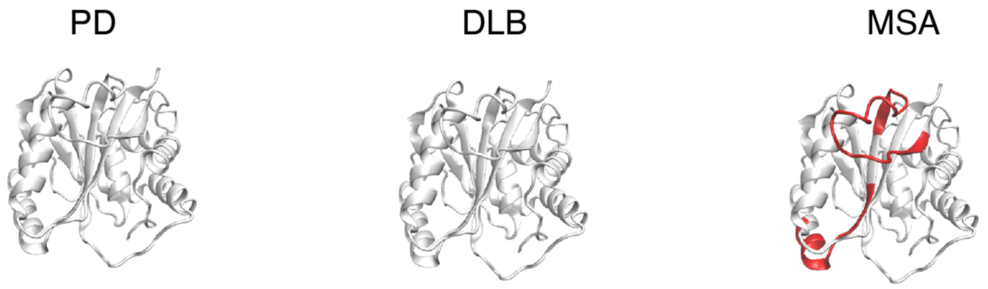


**Supplementary Figure 23. UCHL1 response to the uptake of αSyn PD, DLB, and MSA fibrillar polymorphs in SH-SY5Y.** Structural models show LiP-MS hit peptides mapped onto UCHL1 (PDBID 2etl) structure that change upon uptake of the different αSyn fibrillar polymorphs (red: FC>2, q-val<0.05).

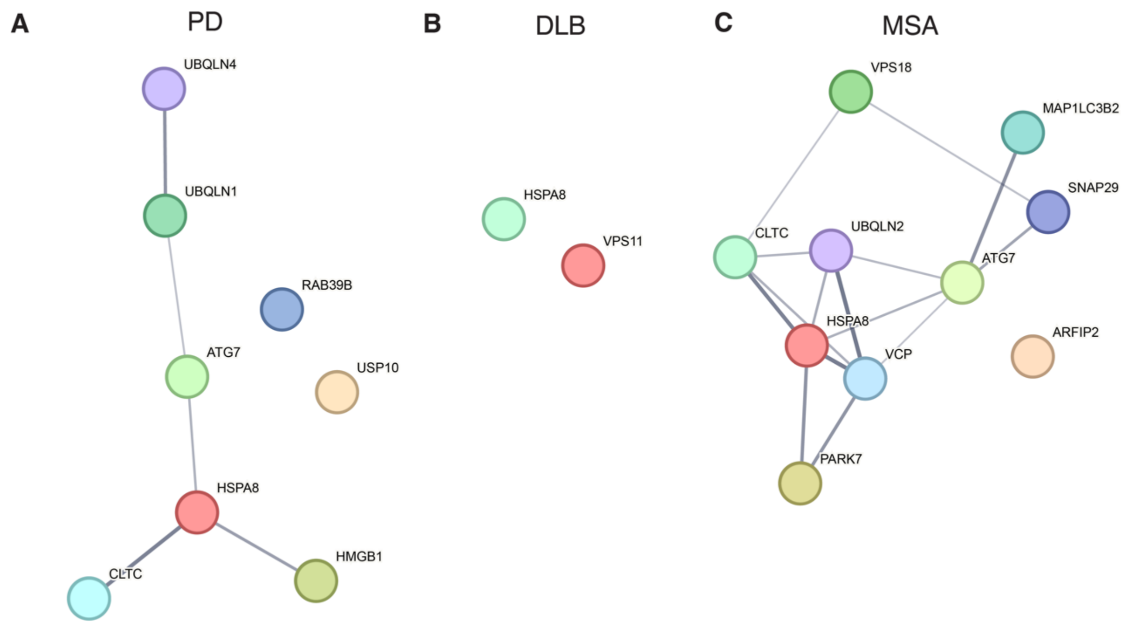


**Supplementary Figure 24. Autophagy-related proteins respond differentially to αSyn fibrils of the three disease strains.** Hit proteins responding to αSyn fibril uptake (FC>1.5, q-val<0.05) and involved in autophagy (based on annotation in Uniprot) were analyzed in String to reveal possible protein networks. Resulting plots are shown for PD (**A**), DLB (**B**), and MSA (**C**) strains. Nodes indicate proteins, edges indicate physical or functional interactions based on Experiments, Databases, and Textmining. Line thickness indicates the strength of the data support.

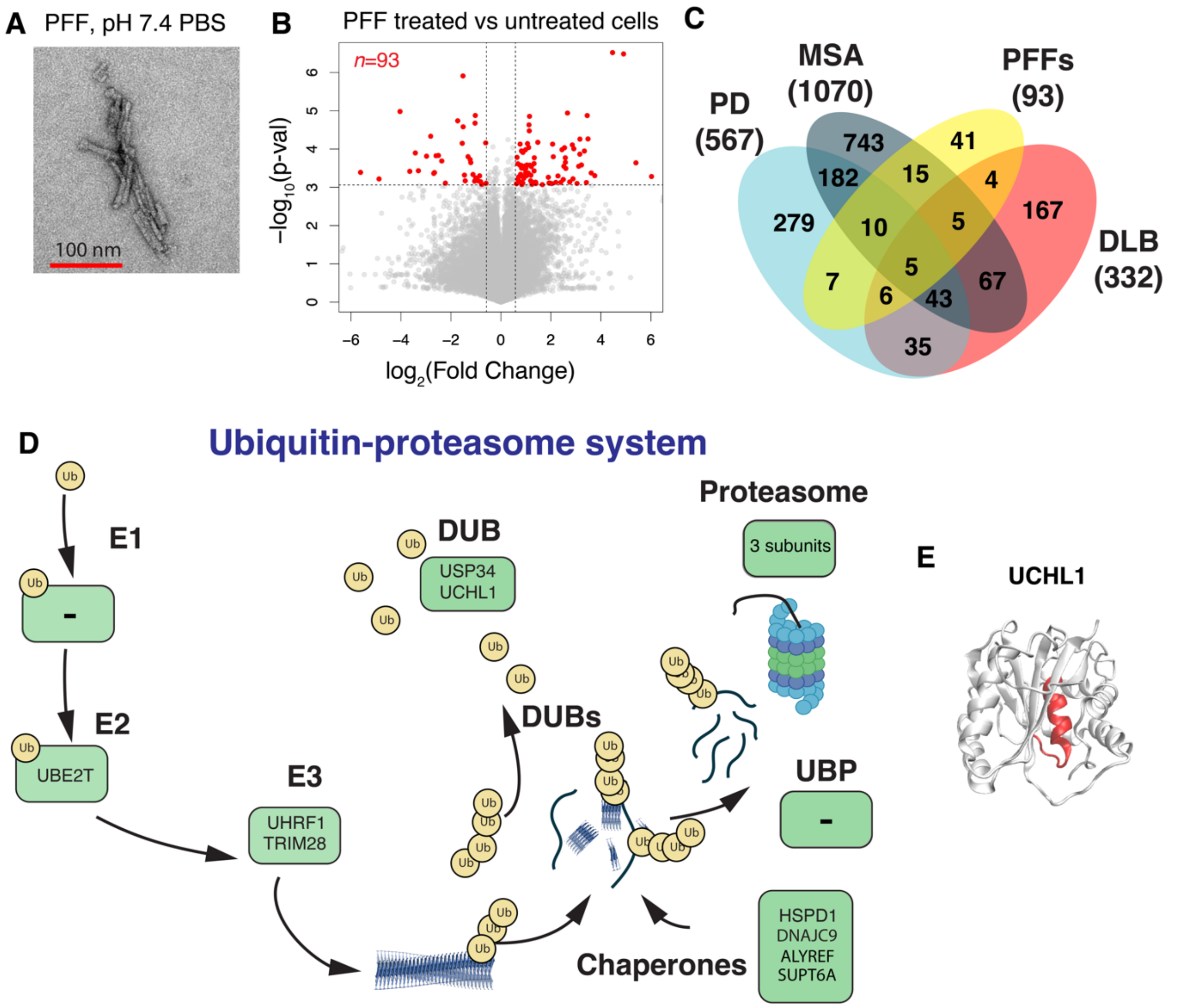


**Supplementary Figure 25. SH-SY5Y cells differentially respond to αSyn PFFs, PD, DLB, and MSA polymorphs.** (**A**) TEM image of fragmented PFFs assembled at pH 7.4 in PBS. (**B**) Volcano plot highlights peptides for which susceptibility to PK changes significantly upon treatment with PFFs. (**C**) The Venn diagram shows the overlap of proteins undergoing protease susceptibility changes upon cell infection with the different disease-derived αSyn fibrillar polymorphs and PFFs (FC>1.5 q-val<0.05). (**D**) The diagram shows proteins annotated to the ubiquitin-proteasomal pathway where at least one peptide is subject to changes upon uptake of αSyn PFFs. (**E**) Structural model shows LiP-MS hit peptide mapped onto UCHL1 (PDBID 2etl) structure that change upon uptake of the αSyn PFFs (red: FC>2, q-val<0.05).

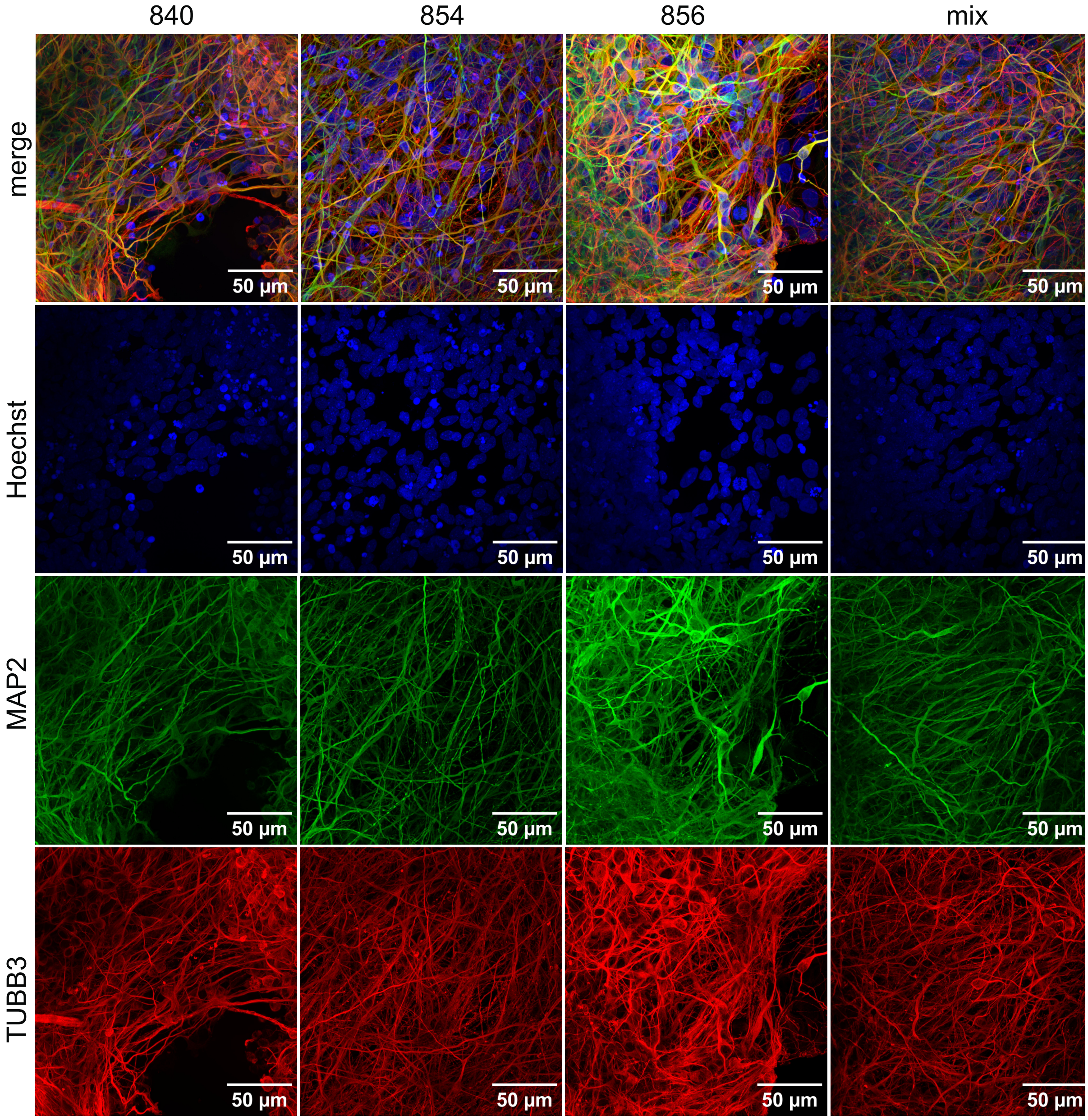


**Supplementary Figure 26. Fluorescence microscopy analysis of iPSCs-derived neurons.** A mix of neurons derived from three patients (840, 854, and 856) was differentiated for 56 days and immunostained with Hoechst, anti-MAP2 (Millipore, MAB378, 1:300) and anti-beta 3 tubulin (TUBB3, BioLegend, 801202, 1:300) antibodies.

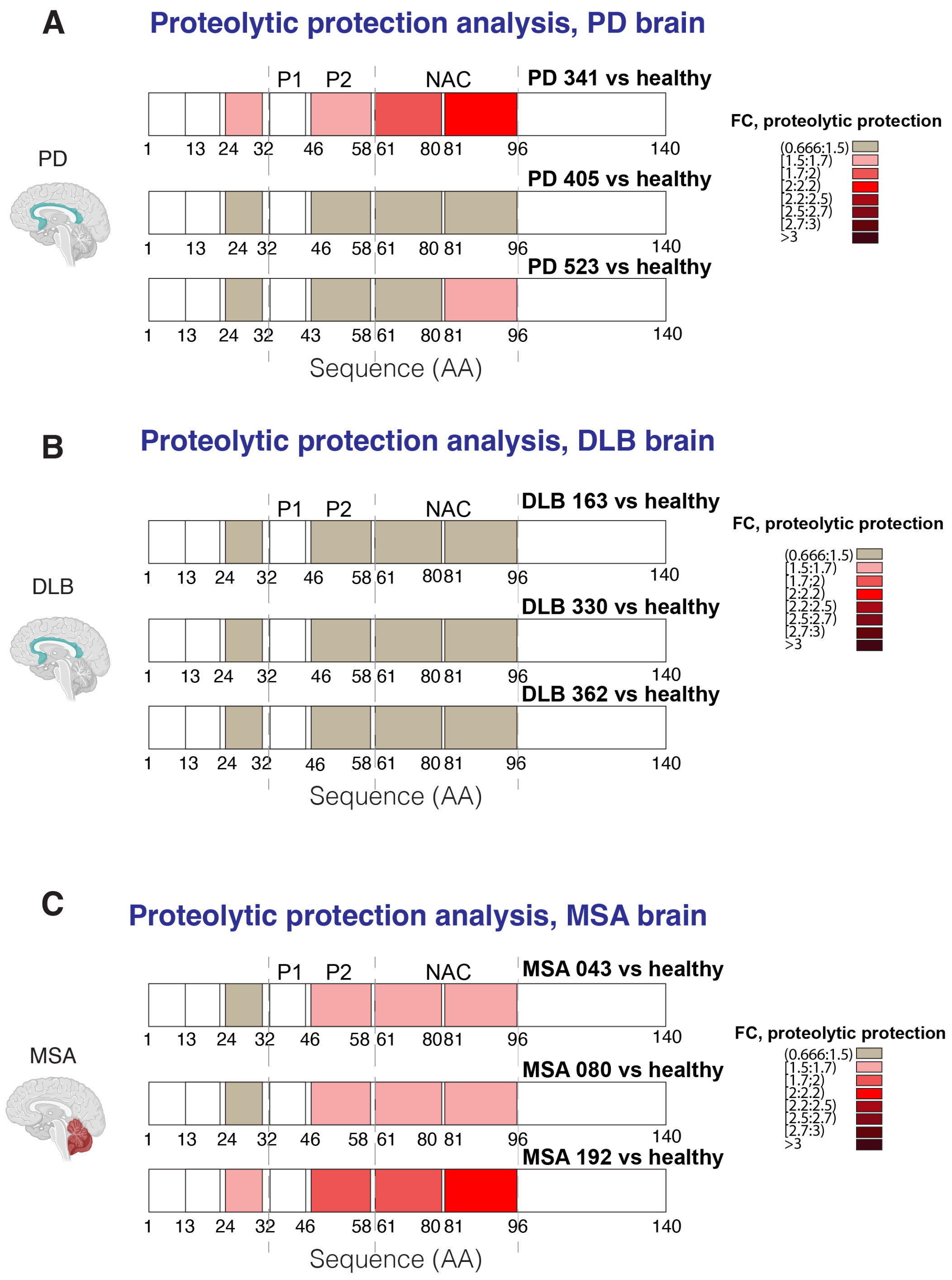


**Supplementary Figure 27. Disease-specific structures of αSyn in brain lysates from individual patients.** LiP-MS-based proteolytic protection analysis of αSyn in PD, DLB, and MSA brain homogenates vs αSyn in healthy group, shown for individual patients. The colour scale shows fold change of proteolytic protection along the αSyn sequence; darker hues show increased protection (n=3 patients per disease and control, n=4 technical replicates per sample). We note that, for patient PD405, peptide 81-96 shows a fold change (1.42) just below that used for our significance cutoff (1.5).

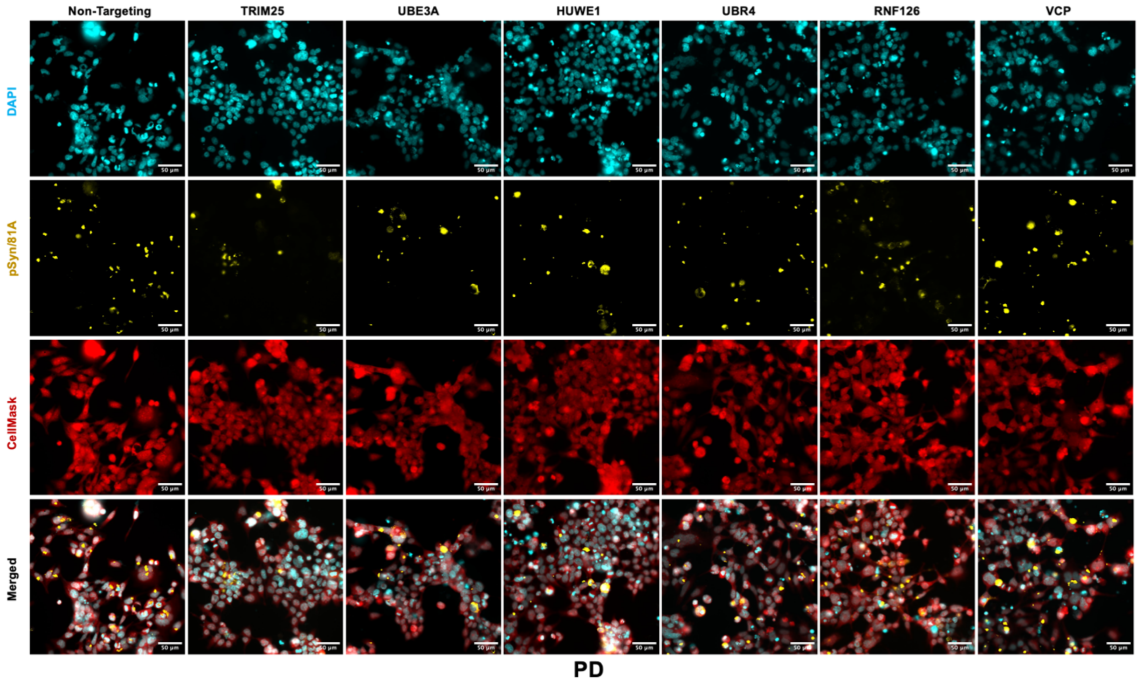


**Supplementary Figure 28. Functional testing of the effect of LiP-MS hits on αSyn accumulation upon uptake of PD fibrils.** The images show cells in which the indicated putative αSyn regulatory factor was genetically upregulated using CRISPR, followed by incubation for 72h with αSyn PD fibrils. The fluorescence stains show pSer129 αSyn inclusions (pSer129/81A yellow), nuclei (DAPI, blue), and cytoplasm and nucleus (whole cell stain, red). NTG (Non-targeting control) indicates plasmids with a scrambled gRNA sequence. Scale bar 50 µm.

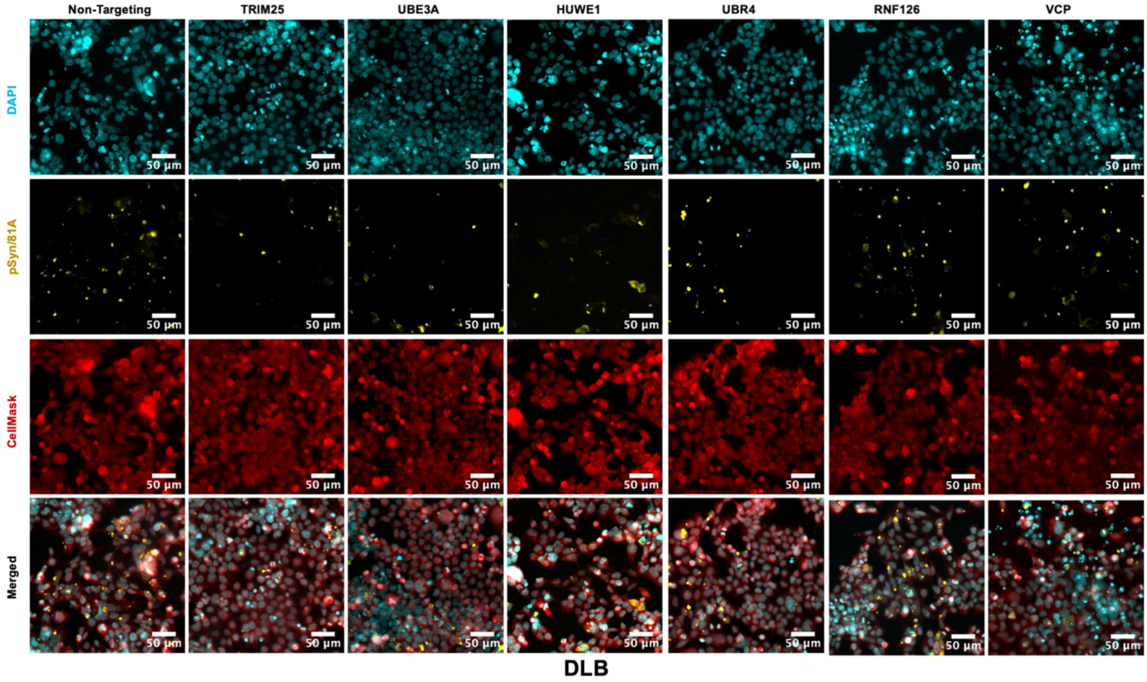


**Supplementary Figure 29. Functional testing of the effect of LiP-MS hits on αSyn accumulation upon uptake of DLB fibrils.** The images show cells in which a single putative αSyn regulatory factor was genetically upregulated using CRISPR, followed by incubation for 72h with αSyn DLB fibrils. The fluorescence stains show pSer129 αSyn inclusions (pSer129/81A yellow), nuclei (DAPI, blue), and cytoplasm and nucleus (whole cell stain, red). NTG (Non-targeting control) indicates plasmids with a scrambled gRNA sequence. Scale bar 50 µm.

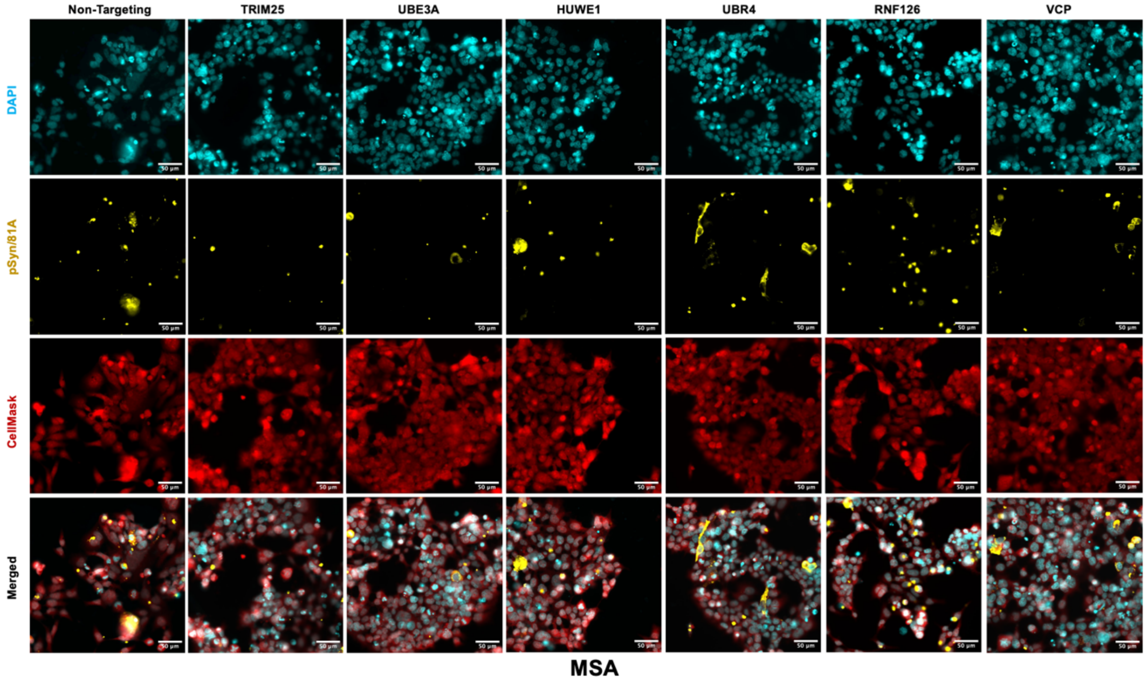


**Supplementary Figure 30. Functional testing of the effect of LiP-MS hits on αSyn accumulation upon uptake of MSA fibrils.** The images show cells in which a single putative αSyn regulatory factor was genetically upregulated using CRISPR, followed by incubation for 72h with αSyn MSA fibrils. The fluorescence stains show pSer129 αSyn inclusions (pSer129/81A yellow), nuclei (DAPI, blue), and cytoplasm and nucleus (whole cell stain, red). NTG (Non-targeting control) indicates plasmids with a scrambled gRNA sequence. Scale bar 50 µm.

| | E3 | | | DUBs | |
|---|---|---|---|---|---|
| **PD** | **DLB** | **MSA** | **PD** | **DLB** | **MSA** |
| DCAF13 | FBXO22 | HACE1 | UCHL1 | WDR48 | WDR48 |
| HUWE1 | HUWE1 | HUWE1 | UCHL5 | UCHL5 | USP15 |
| RANBP2 | SKP1 | IRF2BP1 | USP7 | UCHL1 | USP5 |
| SKP1 | CUL4A | LRRC41 | OTUB1 | USP7 | USP9X |
| UFL1 | BIRC6 | UHRF1 | SENP3 | | VCPIP1 |
| CAND1 | CUL1 | UBR4 | UFSP2 | | OTUB1 |
| DCUN1D1 | TRIM28 | UBE3A | HINT1 | | HINT1 |
| UHRF1 | | RanBP2 | USP13 | | UCHL1 |
| UBR5 | | CACYBP | USP14 | | UCHL5 |
| UBE4A | | CDC23 | USP5 | | USP11 |
| FBXL18 | | CAND1 | | | USP14 |
| CACYBP | | DDB1 | | | USP4 |
| CUL4B | | ELOC | | | EIF3F |
| ELOC | | HLTF | | | |
| PRPF19 | | LRRC41 | | | |
| TRIM24 | | PRPF19 | | | |
| TRIM28 | | TRIM28 | | | |
| UBR4 | | UBR7 | | | |
| UBE3A | | | | | |

**Supplementary Table 1. Lists of E3 ligases and DUBs structurally responding to the disease fibrils after spiking into SH-SY5Y lysate for 15 min.** Hits are coloured (text) based on the significance cut off (black: FC>2, q-val<0.05; magenta: FC>1.5, q-val<0.05).

| Proteases | | |
|---|---|---|
| **PD** | **DLB** | **MSA** |
| COPS5 | COPS5 | AGTPBP1 |
| XPNPEP1 | RNPEP | XPNPEP1 |
| NPEPPS | CAPN1 | RNPEP |
| RNPEP | CPD | CPD |
| BLMH | MIPEP | CTSD |
| CNDP2 | UCHL5 | NUP98 |
| ERAP1 | UCHL1 | TPP2 |
| IDE | USP7 | USP15 |
| PMPCB | AGTPBP1 | USP5 |
| PSMC2 | CLPP | USP9X |
| SCPEP1 | THOP1 | USP9Y |
| UCHL1 | | VCPIP1 |
| UCHL5 | | AFG3L2 |
| USP7 | | OTUB1 |
| AGTPBP1 | | CNDP2 |
| OTUB1 | | DPP3 |
| PARK7 | | EIF3F |
| SENP3 | | HINT1 |
| UFSP2 | | PEPD |
| CAPN1 | | PSMB6 |
| CLPP | | PSMC2 |
| CTSD | | PSMD14 |
| HINT1 | | UCHL1 |
| NLN | | USP11 |
| PSMD14 | | USP14 |
| TPP2 | | USP4 |
| USP13 | | |
| USP14 | | |
| USP5 | | |

**Supplementary Table 2. Lists of proteases structurally responding to αSyn strains after spiking into SH-SY5Y lysate for 15 min.** Hits are coloured (text) based on the significance cut off (black: FC>2, q-val<0.05; magenta: FC>1.5, q-val<0.05).

| Proteases | | |
|---|---|---|
| **PD** | **DLB** | **MSA** |
| ACE | UFSP2 | OTUD6B |
| BLMH | THOP1 | NPEPPS |
| CPD | USP19 | BLMH |
| ERAP1 | USP47 | CAPN1 |
| HINT1 | AFG3L2 | CNDP2 |
| AFG3L2 | CLPP | CASP3 |
| RNPEP | CTSL | CTSA |
| LTA4H | | DPP3 |
| LONP1 | | DPP9 |
| PSMC2 | | IDE |
| USP14 | | NUP98 |
| USP10 | | PREP |
| | | UCHL1 |
| | | USP4 |
| | | ADAM10 |
| | | AFG3L2 |
| | | OTUB1 |
| | | PARK7 |
| | | UFSP2 |
| | | DPP6 |
| | | EIF3F |
| | | HINT1 |
| | | NUP98 |
| | | PSMD14 |
| | | TPP2 |
| | | USP11 |
| | | USP35 |
| | | USP4 |

**Supplementary Table 3. Lists of proteases structurally responding to disease-derived αSyn fibrillar polymorph uptake by SH-SY5Y live cells.** Hits are coloured (text) based on the significance cut off (black: FC>2, q-val<0.05; magenta: FC>1.5, q-val<0.05).

| Wikipathway | DLB hits Count | P value |
|---|---|---|
| WP534:Glycolysis and gluconeogenesis | 10 | 1.05E-05 |
| WP1946:Cori cycle | 7 | 2.54E-05 |
| WP4628:Aerobic glycolysis augmented | 6 | 1.40E-04 |
| WP4629:Aerobic glycolysis | 6 | 1.40E-04 |
| WP3888:VEGFA VEGFR2 signaling | 27 | 2.26E-04 |
| WP4290:Metabolic reprogramming in colon cancer | 9 | 7.85E-04 |
| WP5355:Metabolic epileptic disorders | 11 | 0.0012922 |
| WP383:Striated muscle contraction pathway | 5 | 0.00187809 |
| WP4018:Clear cell renal cell carcinoma pathways | 9 | 0.00242735 |
| WP5193:Cholesterol synthesis disorders | 5 | 0.00657619 |
| WP706:Sudden infant death syndrome SIDS susceptibility pathways | 7 | 0.02174297 |
| WP5426:HDAC6 interactions in the central nervous system | 9 | 0.02179481 |
| WP5333:Enterocyte cholesterol metabolism | 5 | 0.0264029 |
| WP289:Myometrial relaxation and contraction pathways | 7 | 0.02683042 |
| WP197:Cholesterol biosynthesis pathway | 4 | 0.02831878 |
| WP183:Proteasome degradation | 7 | 0.0326534 |
| WP2884:NRF2 pathway | 6 | 0.03926318 |
| WP5329:Cholesterol biosynthesis pathway in hepatocytes | 5 | 0.04039139 |
| WP2359:Parkin ubiquitin proteasomal system pathway | 7 | 0.0592287 |

**Supplementary Table 4.** Pathway enrichment analysis (WikiPathways) for the set of proteins that show structural changes in neurons treated with DLB fibrils (p-val <0.05).

| Wikipathway | MSA hits Count | P value |
|---|---|---|
| WP183:Proteasome degradation | 26 | 1.92E-06 |
| WP5376:17p13 3 YWHAE copy number variation | 11 | 6.21E-04 |
| WP314:Fas ligand pathway and stress induction of heat shock proteins | 13 | 9.72E-04 |
| WP4718:Cholesterol metabolism with Bloch and Kandutsch Russell pathways | 15 | 0.00177423 |
| WP534:Glycolysis and gluconeogenesis | 15 | 0.00177423 |
| WP5193:Cholesterol synthesis disorders | 10 | 0.00178096 |
| WP3888:VEGFA VEGFR2 signaling | 73 | 0.00213816 |
| WP197:Cholesterol biosynthesis pathway | 9 | 0.00261463 |
| WP4629:Aerobic glycolysis | 8 | 0.00381255 |
| WP4190:Mevalonate arm of cholesterol biosynthesis pathway | 8 | 0.00381255 |
| WP4628:Aerobic glycolysis augmented | 8 | 0.00381255 |
| WP5333:Enterocyte cholesterol metabolism | 12 | 0.00392345 |
| WP5124:Alzheimer 39 s disease | 43 | 0.00408537 |
| WP2059:Alzheimer 39 s disease and miRNA effects | 43 | 0.00408537 |
| WP5355:Metabolic epileptic disorders | 24 | 0.00555933 |
| WP4290:Metabolic reprogramming in colon cancer | 17 | 0.00656341 |
| WP4018:Clear cell renal cell carcinoma pathways | 19 | 0.00724847 |
| WP2359:Parkin ubiquitin proteasomal system pathway | 21 | 0.00762465 |
| WP5329:Cholesterol biosynthesis pathway in hepatocytes | 12 | 0.01280461 |
| WP1946:Cori cycle | 8 | 0.01298829 |
| WP5304:Cholesterol metabolism | 14 | 0.02001194 |
| WP4949:16p11 2 proximal deletion syndrome | 15 | 0.02333649 |
| WP5153:N glycan biosynthesis | 12 | 0.02418329 |
| WP3963:Mevalonate pathway | 5 | 0.02520266 |
| WP4804:Cholesterol biosynthesis with skeletal dysplasias | 5 | 0.02520266 |
| WP5114:Nucleotide excision repair in xeroderma pigmentosum | 18 | 0.03186722 |
| WP3925:Amino acid metabolism | 21 | 0.0322771 |
| WP2369:Histone modifications | 14 | 0.03315797 |
| WP698:Glucuronidation | 5 | 0.04852685 |
| WP3871:Valproic acid pathway | 5 | 0.04852685 |

**Supplementary Table 5** Pathway enrichment analysis (WikiPathways) for the set of proteins that show structural changes in neurons treated with MSA fibrils (p-val <0.05).

| Wikipathway | PD hits Count | P value |
|---|---|---|
| WP2267~Synaptic vesicle pathway | 20 | 4.65E-07 |
| WP534~Glycolysis and gluconeogenesis | 13 | 0.00121081 |
| WP706~Sudden infant death syndrome (SIDS) susceptibility pathways | 17 | 0.00249845 |
| WP5352~10q11.21q11.23 copy number variation syndrome | 13 | 0.00549347 |
| WP4290~Metabolic reprogramming in colon cancer | 13 | 0.00709031 |
| WP5233~Arsenic metabolism and reactive oxygen species generation | 6 | 0.0123602 |
| WP78~TCA cycle (aka Krebs or citric acid cycle) | 8 | 0.01556097 |
| WP536~Calcium regulation in cardiac cells | 20 | 0.01605956 |
| WP500~Glycogen synthesis and degradation | 10 | 0.01894123 |
| WP383~Striated muscle contraction pathway | 6 | 0.01976094 |
| WP289~Myometrial relaxation and contraction pathways | 18 | 0.02116647 |
| WP2359~Parkin-ubiquitin proteasomal system pathway | 14 | 0.03152441 |
| WP4932~7q11.23 copy number variation syndrome | 13 | 0.03619362 |
| WP4756~Renin-angiotensin-aldosterone system (RAAS) | 7 | 0.04016533 |
| WP4698~Vitamin D-sensitive calcium signaling in depression | 7 | 0.04016533 |
| WP3298~Melatonin metabolism and effects | 5 | 0.04053488 |
| WP3679~Cell-type dependent selectivity of CCK2R signaling | 5 | 0.04053488 |
| WP2118~Arrhythmogenic right ventricular cardiomyopathy | 11 | 0.04776238 |
| WP5124~Alzheimer's disease | 29 | 0.04855935 |
| WP2059~Alzheimer's disease and miRNA effects | 29 | 0.04855935 |

**Supplementary Table 6.** Pathways enrichment analysis (WikiPathways) for the set of proteins that show structural changes in patient brains afflicted by PD (p-val <0.05).

| Wikipathway | DLB hits Count | P value |
|---|---|---|
| WP534~Glycolysis and gluconeogenesis | 19 | 6.63E-08 |
| WP4290~Metabolic reprogramming in colon cancer | 17 | 6.23E-05 |
| WP706~Sudden infant death syndrome (SIDS) susceptibility pathways | 18 | 0.00127209 |
| WP2267~Synaptic vesicle pathway | 15 | 0.00130544 |
| WP2359~Parkin-ubiquitin proteasomal system pathway | 17 | 0.00282581 |
| WP4018~Clear cell renal cell carcinoma pathways | 16 | 0.00297169 |
| WP4917~Proximal tubule transport | 9 | 0.00302602 |
| WP4629~Aerobic glycolysis | 7 | 0.00718467 |
| WP5049~Glycolysis in senescence | 6 | 0.00824285 |
| WP1946~Cori cycle | 7 | 0.01152649 |
| WP2272~Pathogenic Escherichia coli infection | 13 | 0.01183644 |
| WP289~Myometrial relaxation and contraction pathways | 19 | 0.01389298 |
| WP2059~Alzheimer's disease and miRNA effects | 32 | 0.01567581 |
| WP5124~Alzheimer's disease | 32 | 0.01567581 |
| WP3888~VEGFA-VEGFR2 signaling | 50 | 0.01644492 |
| WP2431~Spinal cord injury | 14 | 0.02023907 |
| WP536~Calcium regulation in cardiac cells | 20 | 0.02257138 |
| WP5344~Cardiomyocyte signaling pathways converging on Titin | 6 | 0.02273813 |
| WP5220~Metabolic reprogramming in pancreatic cancer | 10 | 0.04537814 |

**Supplementary Table 7.** Pathways enrichment analysis (WikiPathways) for the set of proteins that show structural changes in patient brains afflicted by DLB (p-val <0.05).

| Wikipathway | MSA hits Count | P value |
|---|---|---|
| WP706~Sudden infant death syndrome (SIDS) susceptibility pathways | 30 | 4.73E-04 |
| WP4290~Metabolic reprogramming in colon cancer | 22 | 9.36E-04 |
| WP534~Glycolysis and gluconeogenesis | 20 | 0.00106238 |
| WP2267~Synaptic vesicle pathway | 24 | 0.00133559 |
| WP2359~Parkin-ubiquitin proteasomal system pathway | 28 | 0.00141021 |
| WP4148~Splicing factor NOVA regulated synaptic proteins | 20 | 0.00187179 |
| WP4767~FGFR3 signaling in chondrocyte proliferation and terminal differentiation | 8 | 0.00313195 |
| WP5220~Metabolic reprogramming in pancreatic cancer | 19 | 0.00343762 |
| WP3925~Amino acid metabolism | 32 | 0.00396205 |
| WP4629~Aerobic glycolysis | 10 | 0.00480077 |
| WP5114~Nucleotide excision repair in xeroderma pigmentosum | 20 | 0.00506305 |
| WP4018~Clear cell renal cell carcinoma pathways | 25 | 0.00648689 |
| WP2369~Histone modifications | 13 | 0.00953743 |
| WP5049~Glycolysis in senescence | 8 | 0.00990353 |
| WP2453~TCA cycle and deficiency of pyruvate dehydrogenase complex (PDHc) | 11 | 0.01062892 |
| WP106~Alanine and aspartate metabolism | 9 | 0.01071271 |
| WP1946~Cori cycle | 10 | 0.01087119 |
| WP5085~Vasopressin-regulated water reabsorption | 16 | 0.01215565 |
| WP4159~GABA receptor signaling | 14 | 0.01495281 |
| WP78~TCA cycle (aka Krebs or citric acid cycle) | 12 | 0.01810431 |
| WP3871~Valproic acid pathway | 6 | 0.02043411 |
| WP179~Cell cycle | 14 | 0.0237743 |
| WP4549~Fragile X syndrome | 42 | 0.02512852 |
| WP2864~Apoptosis-related network due to altered Notch3 in ovarian cancer | 13 | 0.02667579 |
| WP5046~NAD metabolism in oncogene-induced senescence and mitochondrial dysfunction-associated senescence | 10 | 0.03719724 |
| WP98~Prostaglandin synthesis and regulation | 10 | 0.03719724 |
| WP500~Glycogen synthesis and degradation | 16 | 0.03996587 |
| WP2064~Neural crest differentiation | 9 | 0.04120123 |
| WP3888~VEGFA-VEGFR2 signaling | 98 | 0.04228711 |

**Supplementary Table 8.** Pathways enrichment analysis (WikiPathways) for the set of proteins that show structural changes in patient brains afflicted by MSA (p-val <0.05).

| | E3 | | | DUBs | |
|---|---|---|---|---|---|
| **PD** | **DLB** | **MSA** | **PD** | **DLB** | **MSA** |
| FBXO41 | BRCC3 | COMMD4 | UCHL1 | MINDY1 | MINDY2 |
| HECTD3 | FBXO2 | FBXL16 | **USP14** | OTUB1 | **OTUB1** |
| CUL1 | FBXO41 | FBXO2 | USP5 | HINT1 | **EIF3F** |
| **CUL3** | FBXO44 | FBXO41 | | **UCHL1** | **HINT1** |
| CUL4B | HUWE1 | HACE1 | | USP5 | **UCHL1** |
| **CAND1** | SKP1 | HECW2 | | | UCHL3 |
| DDB1 | UBAC1 | **HUWE1** | | | UCHL5 |
| ITCH | WSB2 | **RANBP2** | | | USP10 |
| MARCHF5 | CUL3 | SKP1 | | | USP14 |
| TRIM2 | CUL4B | SUGT1 | | | USP15 |
| | DDB1 | UBAC1 | | | USP46 |
| | MUL1 | **UFL1** | | | USP47 |
| | **UBR4** | CACYBP | | | USP5 |
| | UBE4A | CUL1 | | | USP9X |
| | **TRIM28** | **CUL2** | | | |
| | | CUL4B | | | |
| | | **CUL5** | | | |
| | | **CAND1** | | | |
| | | DDB1 | | | |
| | | GAN | | | |
| | | RNF123 | | | |
| | | RBX1 | | | |
| | | **TRIM2** | | | |
| | | **TRIM25** | | | |
| | | **TRIM28** | | | |
| | | UBR1 | | | |
| | | **UBR4** | | | |
| | | UBE3C | | | |
| | | UBE4A | | | |

**Supplementary Table 9. Lists of E3 ligases and DUBs structurally altered in brains of patients suffering from PD, DLB, and MSA.** Hits are coloured (text) based on the significance cut off (black: FC>2, q-val<0.05; magenta: FC>1.5, q-val<0.05). Orange, cyan, and red shades highlight the overlapping LiP-MS hits (i.e., proteins that show structural changes) between the cell seeding experiment in SH-SY5Y cells (orange), in neurons (cyan), or in both (red) and the comparison of brain proteomes (q-val< 0.05, FC >1.5).

| Sample | Repl. 1 Fold change mRNA | Repl. 2 Fold change mRNA | Fold change Protein level |
|---|---|---|---|
| TRIM25 | 2.1 | 2.9 | 2.2 |
| UBE3A | 2.0 | 2.1 | 1.6 |
| VCP | 2.4 | 11.0 | 1.5 |
| HUWE1 | 21.4 | 2.6 | 1.1 |
| RNF126 | 2.0 | 1.6 | 1.5 |
| UBR4 | 1.2 | 1.9 | 1.1 |

**Supplementary Table 10. Control of gene activation in CRISPR-edited HEK293 cells.** Fold change of mRNA and protein levels in HEK cells where corresponding gene was activated.
